# Ubiquitin Receptor-Mediated, Ubiquitin-Independent Targeted Protein Degradation via 26S Proteasomes

**DOI:** 10.1101/2025.08.18.670774

**Authors:** Seh Hoon Park, Yejin Jang, Soo-Yeon Lee, Eunseo Kim, Dawon Jeong, Insuk Byun, Jiseong Kim, Jisoo Yang, Chang Han Lee, Dohyun Han, Jae-Hwan Nam, Min Jae Lee

## Abstract

The 26S proteasome engages with ubiquitinated substrates primarily through its constituent ubiquitin (Ub) receptors, which initiates a cascade of proteolytic processes. Leveraging this recognition mechanism, we developed a targeted protein degradation (TPD) strategy that recruits substrates directly to the proteasome, thereby bypassing the ubiquitination step. Our proteasome-targeting chimera, Protea-Tac, is a heterobifunctional protein degrader composed of a Ub receptor and an intracellular antibody. This chimera integrates into 26S proteasomes without altering their functional integrity. Localization of target proteins, including c-Fos, BRD4, Flag-TDP43, HA-tau, and GFP-ODC, to the proteasome via Protea-Tac with cognate antibodies resulted in their induced degradation. We demonstrated that this platform is 1) modular, allowing facile switching between targets; 2) Ub-independent; and 3) highly target-specific. Protea-Tac exhibited *in vivo* anti-tumor efficacy, degrading c-Fos and substantially delaying tumor growth. Overall, these findings identify Protea-Tac as a distinct TPD modality capable of directly degrading intracellular proteins via engineered 26S proteasomes.

**Teaser:** Protea-Tac is a heterobifunctional protein degrader that enables Ub-independent TPD, exhibits high specificity and modularity, and demonstrates *in vivo* applicability.

## Introduction

Targeted protein degradation (TPD) has evolved from a conceptual framework to a emerging therapeutic modality that selectively eliminates disease-related proteins (*1*). Unlike conventional small-molecule drugs that typically rely on high-affinity binding, TPD can accomplish a catalytic mechanism that allows for effective protein turnover at moderate-to-low binding affinity (*2, 3*). Proteolysis-targeting chimera (PROTAC) is at the forefront of TPD therapeutics. It is a synthetic heterobifunctional molecule composed of a target binding ligand and a ligand for a cellular E3 ubiquitin (Ub) ligase. The induced proximity via PROTAC promotes substrate polyubiquitination (hereafter referred to as ubiquitination) and, subsequently, degradation by 26S proteasomes (*4–8*). Many other TPD platforms, including molecular glue degraders (MGDs), employ a similar mode of action to PROTACs (*9–11*). These approaches offer distinct advantages over traditional therapeutics, particularly in targeting proteins previously considered undruggable, such as scaffold proteins and transcription factors (*12, 13*). Additionally, TPD has the potential to provide a durable therapeutic solution for acquired drug resistance, which is often caused by mutations in neoplastic targets (*14–17*).

The key mechanistic basis of these TPD modalities is their exploitation of the endogenous Ub-proteasome system (UPS), the primary proteolytic pathway in all eukaryotes (*18*). In the UPS, E3 Ub ligases, which have over 600 members encoded in the human genome, serve as critical regulator of substrate specificity, mediating the proximity between substrates and Ub moieties (charged by E2 Ub-conjugating enzymes). Both PROTACs and MGDs aim to emulate this biochemical process by forming an induced ternary complex in precise spatial proximity and orientation. Nevertheless, finding suitable ligands for target-E3 ligase engagement remains challenging, particularly for recalcitrant proteins lacking well-defined hydrophobic pockets or possessing intrinsically disordered regions. The development process typically consists of multiple iterative cycles of chemical design, synthesis, functional validation, and optimization. However, the developed small molecules tend to have high molecular weights (>800 Da) and large polar surface areas, resulting in relatively low solubility and limited cell permeability. The efficacy of Ub inducers is further constrained by the availability of lysine residues on the neo-substrates and the cellular abundance of E3 enzymes; at present, two E3s, CRBN and VHL, account for more than 90% of current PROTACs and MGDs (*19–22*).

The proteolytic “processor” in the UPS is the 26S proteasome holoenzyme, which is responsible for the majority of intracellular protein degradation (at least 80% in proliferating cells) (*23*). Given its high cellular abundance (> 200 nM) in most cell types, the proteasome is an ideal degradation machinery for TPD (*24–27*). Moreover, 26S proteasomes appear to have reserved proteolytic capacity for neo-substrates, because a ∼ 30% decline in 26S proteasome levels after *PSMD2* knockdown did not significantly affect global protein turnover (*28–30*). The 26S proteasome is composed of two distinct subcomplexes: the 20S catalytic particle (CP) and the 19S regulatory particle (RP), which can reversibly associate and dissociate in response to cellular stress (*31–33*). The 26S proteasome also exemplifies an evolutionary dichotomy: the ancient CP for proteolysis and the later-evolved RP for Ub recognition and processing (*34*). Most Ub chains are recognized by Ub receptors (UbRs) located in the RP, including PSMD2/Rpn1, PSMD4/Rpn10, and ADRM1/Rpn13. Interaction between substrates and these subunits initiates a cascade of coordinated processes, including “gate” opening of the CP, conformational remodeling of the RP, and alignment of the RP-CP axial channel, allowing efficient substrate translocation (*35–38*). When the substrate’s unstructured region engages with the pore of the PSMC/Rpt ATPase ring, nucleotide hydrolysis generates mechanical force, simultaneously unfolding the substrate and translocating it into the hydrolytic chamber (*39*).

To circumvent the mechanistic limitations of E3-dependent TPD strategies, alternative degraders, some of which hijack the lysosomal or autophagic pathways, have been developed (*40–44*). In parallel, Ub-independent target degradation has largely been achieved by proximity-driven substrate recruitment to the 26S proteasome, as initially demonstrated using inducible recruitment systems and engineered degrons (*45–47*). Such substrate tethering is not restricted to artificial settings: pathogen effectors and endogenous cofactors could engage substrates to the proteasome to promote proteolysis bypassing ubiquitination (*48, 49*). More recently, compounds that recruit neo-substrates directly to UbRs or other RP components have demonstrated the therapeutic potential of Ub-independent TPD (*50–54*). While promising, expanding these chemical approaches to new targets remains dependent on the availability of suitable small-molecule ligands and often requires extensive chemical optimization. Genetically-encoded degraders offer a complementary strategy that can expand the scope of TPD targets. Their modular, protein-based recognition elements may facilitate the targeting of ligand-poor, conformation-selective targets (*55, 56*). At the same time, the translational development of protein-based degraders requires solutions for challenges such as intracellular delivery and potential immunogenicity.

In this study, we exploited the biochemical feature of the proteasome-residing UbRs to develop a genetically-encoded, directly 26S proteasome-targeting chimera (Protea-Tac). By fusing PSMD4 or ADRM1 to a substrate-targeting antibody (tAb; single-chain variable fragment [scFv] or nanobody [VHH]), we created a modular degrader system that integrates into 26S proteasomes while preserving their proteolytic activity. Biochemical analyses demonstrated that Protea-Tac is modular and heterobifunctional and can induce Ub-independent TPD. Tandem mass tag (TMT)- based quantitative proteomics revealed its high selectivity. Additionally, *in vivo* experiments showed that Protea-Tac degraders post-translationally reduced target proteins and inhibited tumor growth across multiple mouse tumor models. This study introduces Protea-Tac as a distinct, Ub-independent TPD platform, broadening the therapeutic potential of TPD.

## Results

### Protea-Tac as a TPD modality that directly engages the 26S proteasome

Based on previous studies, which revealed low micromolar affinity interactions between the UbRs and Ub chains, we hypothesized that intracellular antibodies such as scFv and VHH could serve as alternatives to Ub chain-mediated substrate recruitment to the 26S proteasome (*57, 58*). We designed the initial Protea-Tac platform by fusing a UbR to an scFv targeting the c-Fos oncoprotein, scFv(c-Fos), at either the N- or C-terminus, so as not to sterically impede the 19S subcomplex assembly (Fig. 1A). To assess TPD activity, various Protea-Tac constructs were transiently expressed in A549 cells. We found that PSMD4-scFv(c-Fos) significantly lowered the steady-state levels of co-transfected c-Fos in an expression level-dependent manner (Fig. 1B). In sharp contrast, *c-Fos* mRNA levels remained unchanged regardless of co-expression of PSMD4-scFv(c-Fos), LacZ^V5^, or their combination (Fig. 1C), implying that the reduction in c-Fos protein occurred post-translationally. Chase experiments demonstrated accelerated c-Fos degradation in the presence of its cognate, PSMD4-based Protea-Tac chimera (Fig. 1D).

**Fig. 1.**
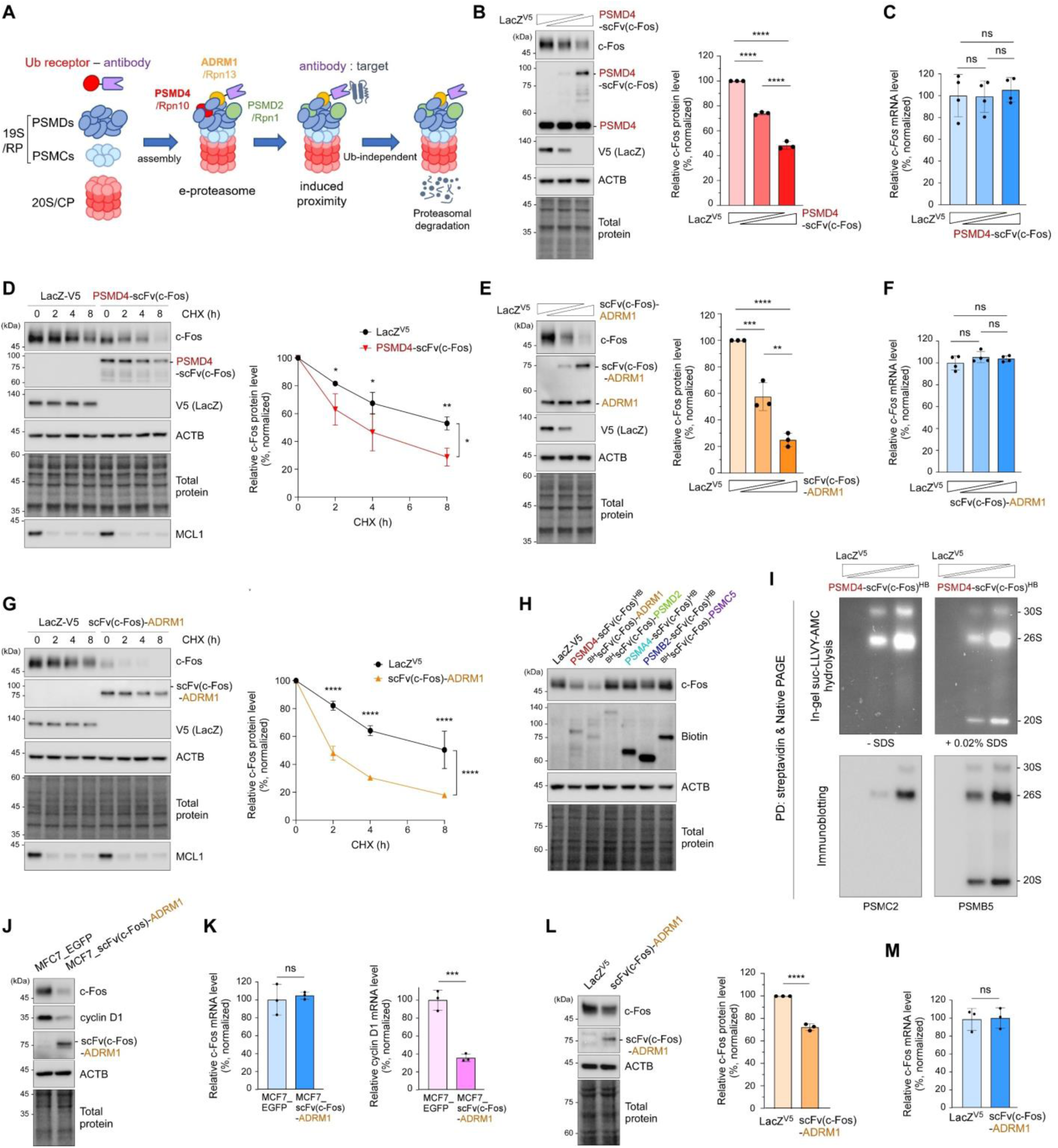
The proteasome-targeting chimera (Protea-Tac) system enables engineered degradation of target proteins through direct proteasome engagement. (**A**) Schematic illustration of targeted protein degradation (TPD) via direct proteasome recruitment by Protea-Tac. (**B**) A549 cells were co-transfected with plasmids expressing PSMD4-scFv(c-Fos) degraders (0, 0.3, and 1 μg) or LacZ^V5^ controls (1, 0.7, and 0 μg), together with c-Fos-expressing plasmids (0.5 μg). Total plasmid DNA amounts and volumes remained constant across all experimental conditions. After 64 h of incubation, whole-cell lysates (WCLs) were analyzed by SDS-PAGE followed by immunoblotting (IB) and total protein staining. *Left*: Representative IB results from three independent experiments are shown. *Right*: Quantification of c-Fos levels normalized to total protein signals. Bars represent the mean ± SD (N = 3). ****p < 0.0001 (one-way ANOVA with Tukey’s post hoc test). (**C**) As in (B) except total RNA was extracted for quantitative RT-PCR (qRT-PCR) using *c-Fos* and *GAPDH* (normalization control) primers. Values represent the mean ± SD of four independent experiments (N = 4). ns, not significant (one-way ANOVA followed by Tukey’s post hoc test). (**D**) Similar to (B) but chase experiments were performed at 2, 4, and 8 h following treatment with 80 μg/mL cycloheximide (CHX) at time zero. c-Fos signals were normalized to the values at 0 h chase (set as 100%) of each group and to total protein levels. Average percentages of remaining c-Fos (mean ± SD) from three independent experiments (N = 3, *p < 0.05, **p < 0.01 from two-way ANOVA followed by Sidak’s post hoc test). MCL1 was used as a physiological proteasome substrate. (**E to G**) The ADRM1-based degrader (scFv(c-Fos)-ADRM1) was evaluated using the same methods as in (B) to (D) (**H**) Various proteasome subunit-based Protea-Tac chimeras were transiently expressed in A549 cells, and c-Fos levels were determined by SDS-PAGE/IB. (**I**) PSMD4-scFv(c-Fos) was C-terminally tagged with a hexahistidine-biotin (HB) tag and transiently expressed for streptavidin affinity purification. Pulldown samples were resolved by native PAGE and analyzed using in-gel suc-LLVY-AMC hydrolysis (*top*) and subsequent IB (*bottom*). The addition of 0.02% SDS activated 20S proteasomes. (**J**) Endogenous c-Fos and cyclin D1 protein levels were compared in MCF7 cells stably expressing EGFP or scFv(c-Fos)-ADRM1. Single-cell clones with the highest expression level were selected for analysis. (**K**) As in (J) except that mRNA levels of endogenous *c-Fos* (*left*) and *cyclin D1* (*right*) were evaluated by qRT-PCR and normalized to *GAPDH*. ****p < 0.0001 (N = 3, two-tailed Student’s *t*-test). (**L**) MCF7 cells were transfected with either LacZ^V5^ or scFv(c-Fos)-ADRM1 for 48 h and analyzed for c-Fos levels by IB. Representative blots (*left*) and quantification (*right*) from three independent experiments are shown. (**M**) Similar to (L) but *c-Fos* mRNA levels were evaluated by qRT-PCR analysis. Values are presented as means ± SD. ****p < 0.0001 (N = 3, two-tailed Student’s *t*-test). ns, not significant. *See* Supplementary Figure 1.

The Protea-Tac degraders with ADRM1 were likewise effective; both ADRM1-scFv(c-Fos) and scFv(c-Fos)-ADRM1 substantially reduced c-Fos protein levels without altering *c-Fos* mRNA (Fig. 1, E and F; fig. S1, A and B). We found that ADRM1-based Protea-Tacs were more effective in c-Fos degradation than PSMD4-based degraders (Fig. 1G; fig. S1C). Notably, fusions of scFv(c-Fos) with PSMD2 or other structural components (either CP or RP subunits) of the 26S proteasome did not significantly affect c-Fos levels (Fig. 1H; fig. S1, D to G). To assess whether and to what extent the cellular 26S proteasome pool is remodeled through Protea-Tac incorporation, we affinity-purified 26S proteasomes and compared the levels of the incorporated chimeras with those of their endogenous UbR counterparts by immunoblotting. Both PSMD4-scFv(c-Fos) and scFv(c-Fos)-ADRM1 were detected in purified proteasomes, albeit at substantially lower levels than endogenous PSMD4 and ADRM1, respectively, suggesting that Protea-Tac is incorporated into only a minor fraction of cellular 26S proteasomes (fig. S1, H and I). We next performed streptavidin-affinity purification using biotin-hexahistidine (BH)-tagged Protea-Tac constructs, revealing structurally intact and enzymatically active 26S proteasomes through non-denaturing (native) electrophoresis and in-gel fluorogenic substrate (suc-LLVY-AMC) hydrolysis (Fig. 1I; fig. S1, J to L). Protea-Tac degraders with a mutant UbR that is incapable of direct proteasomal interaction did not show any TPD activity (fig. S1M). Collectively, these results show that direct substrate tethering to the 26S proteasome can trigger TPD when mediated by permissive UbR configurations, such as PSMD4- and ADRM1-based Protea-Tacs.

Having validated Protea-Tac functionality using transient transfection, we next generated stable cell lines overexpressing scFv(c-Fos)-ADRM1 in human breast cancer (MCF7) and colorectal cancer (HCT116) cells, where c-Fos upregulation is implicated in tumorigenesis (*59, 60*). The selected clones expressing the greatest amounts of scFv(c-Fos)-ADRM1 exhibited marked endogenous c-Fos degradation (72.9% and 58.6% in MCF7 and HCT116, respectively) compared to control cells expressing EGFP (Fig. 1J; fig. S1N). Again, the mRNA levels of endogenous *c-Fos* remained unaffected (Fig. 1K). It was noticeable that protein and mRNA levels of cyclin D1, a direct transcriptional target of c-Fos, were substantially reduced in these stable cells (Fig. 1, J and K; fig. S1O) (*61, 62*). Transient expression of scFv(c-Fos)-ADRM1 in MCF7 and HCT116 cells also reduced endogenous c-Fos levels but to a lesser extent than when stably overexpressed (Fig. 1, L and M; fig. S1, P and Q), likely owing to higher proportion of Protea-Tac-expressing cells and sustained expression across the stable cell populations.

### Mechanistic exploration of Ub-independent, proteasomal degradation by Protea-Tac

To understand the mechanistic basis of Protea-Tac-mediated TPD, we first investigated whether both modules, the tAb and UbR, are necessary and whether their fusion is essential. A panel of control constructs expressing only scFv(c-Fos), UbR alone (PSMD4 or ADRM1), or their co-expression did not affect cellular c-Fos levels, whereas the chimeric Protea-Tac constructs effectively degraded c-Fos (Fig. 2, A and B). We also successfully achieved TPD using VHH as the tAb module, which also demonstrated that both modules are necessary and should be fused for degradation (fig. S2, A and B). To further investigate this heterobifunctionality, we generated constructs that fused scFv(c-Fos) and ADRM1 with or without a self-cleaving T2A sequence at their junction, followed by a P2A-mCherry reporter (Fig. 2C). Flow cytometry analysis of A549 cells stably expressing EGFP-tagged c-Fos (EGFP-c-Fos) indicated significantly lower EGFP signals in the mCherry-positive cell population that expresses the fused scFv(c-Fos)-ADRM1 than those expressing the T2A-separated format (Fig. 2D).

**Fig. 2.**
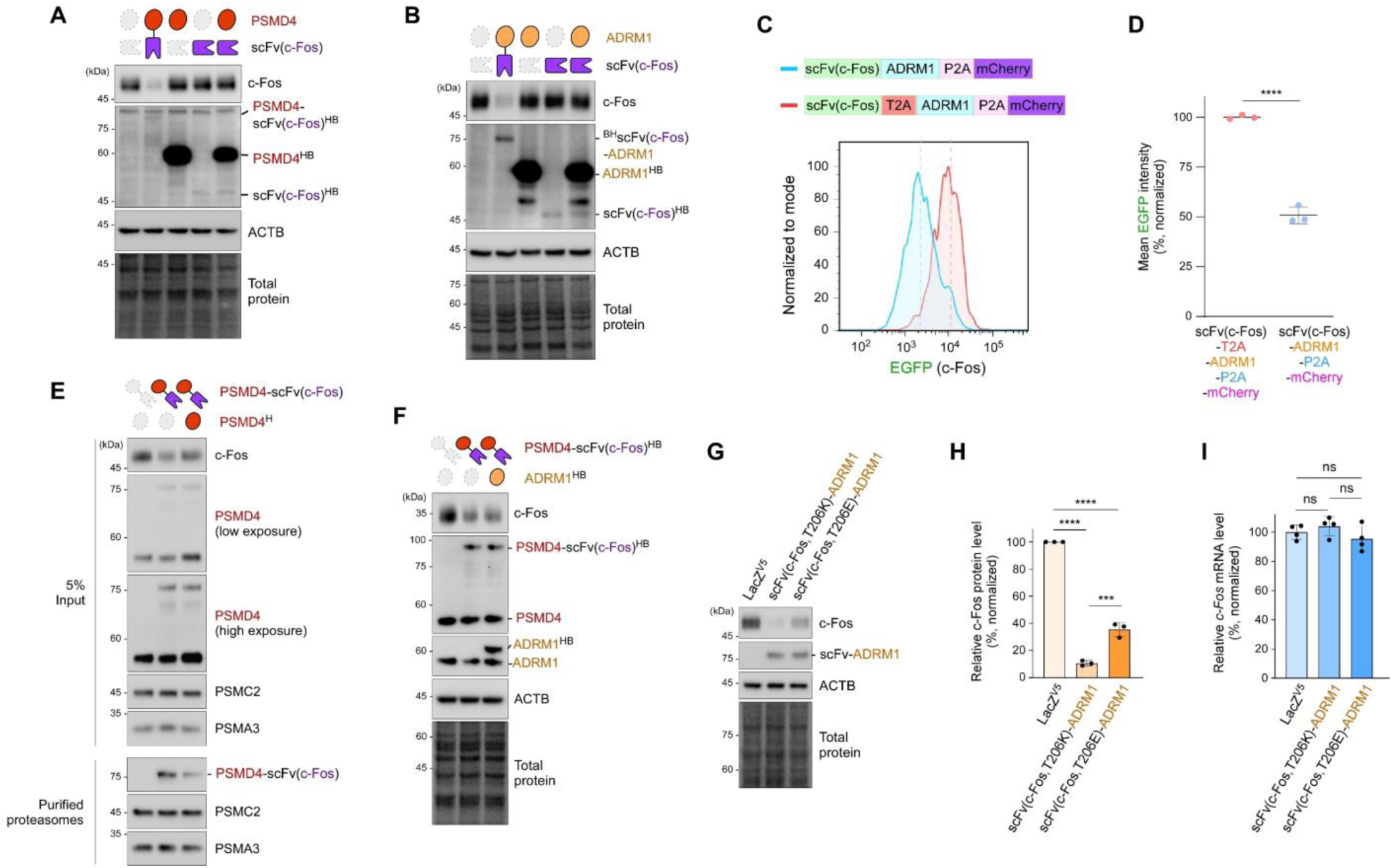
Protea-Tac degraders require both Ub receptor (UbR) and targeting antibody (tAb) modules. (**A**) A549 cells were transfected with plasmids expressing PSMD4, scFv(c-Fos), their combination without fusion, or as a PSMD4-scFv(c-Fos) fusion, along with c-Fos-expressing plasmids. LacZ^V5^ plasmids were added to maintain a total plasmid DNA amount (1.5 μg). WCLs were harvested at 64 h post-transfection and examined using SDS-PAGE/IB, followed by total protein staining. (**B**) Same as (A), but ADRM1 was used as the UbR component. (**C**) Schematic of bicistronic (blue) and tricistronic (red) constructs encoding scFv(c-Fos)-ADRM1 without and with a T2A sequence at the tAb–UbR junction, respectively. mCherry was included downstream of a P2A sequence as a reporter. These constructs were transfected into HEK293T cells stably expressing EGFP-c-Fos for 72 h, and degradation was monitored by flow cytometry. Representative histograms show EGFP-c-Fos fluorescence in mCherry-positive cells. (**D**) Mean fluorescence intensity (MFI) of EGFP in mCherry-positive cells. Dots indicate the average MFI (normalized) per experiment, with bars showing the mean ± SD. ****p < 0.0001 (N = 3, two-tailed Student’s t test). (**E**) HEK293 cells stably expressing biotin-tagged PSMB2 were transfected as in (A), but PSMD4^H^ was co-expressed with PSMD4-scFv(c-Fos) to assess competitive incorporation into the proteasome and target degradation efficiency. Affinity purification of proteasome was conducted and samples were examined by denaturing SDS-PAGE/IB using antibodies against c-Fos and proteasome subunits. (**F**) PSMD4-scFv(c-Fos) was co-expressed with non-cognate ADRM1 to assess its effect on c-Fos degradation. (**G**) scFv(c-Fos) variants with lower (T206K) or higher (T206E) binding affinities were evaluated as the tAb component in ADRM1-based degraders. HEK293T cells were co-transfected with scFv(c-Fos)-ADRM1 (T206K or T206E, 1 μg) or LacZ^V5^ (1 μg) with c-Fos plasmids (0.5 μg). WCLs were analyzed using SDS-PAGE/IB. Representative IB results from three independent experiments are shown. (**H**) Quantification of c-Fos band intensities from (G), normalized to total protein. Bars indicate the mean ± SD (N = 3). **p < 0.01, ***p < 0.001, ****p < 0.0001 (One-way ANOVA followed by Tukey’s multiple comparison test). (**I**) As in (G) except total RNA was extracted for qRT-PCR using *c-Fos* and *GAPDH* (normalization control) primers. Values represent the mean ± SD of four independent experiments (N = 4). ns, not significant (one-way ANOVA followed by Tukey’s post hoc test). *See* Supplementary Figure 2.

To further dissect the mechanism and functionality of Protea-Tac, we co-expressed free PSMD4 or ADRM1 alongside Protea-Tac degraders with the cognate UbR module. Co-expression of the cognate free UbR competitively reduced the incorporation of Protea-Tac into 26S proteasomes and attenuated its TPD activity (Fig. 2E; fig. S2C). In contrast, excess PSMD4 or ADRM1 had no effect on Protea-Tacs with non-cognate UbRs (Fig. 2F; fig. S2D), indicating that Protea-Tac-mediated degradation is primarily associated with its incorporation into the 26S proteasome holoenzyme rather than with stand-alone UbRs. Similarly, co-expression of cognate free UbRs attenuated the TPD activity of VHH-based degraders (fig. S2, E and F). Protea-Tac chimeras with non-cognate antibodies did not affect c-Fos levels (fig. S2, G and H). When we substituted the targeting antibody’s CDR region with that of IgG, Protea-Tac activity was completely abolished (fig. S2, I to K). We then investigated the impact of tAb affinity on degradation efficiency, using scFv(c-Fos) variants with varying affinities for recombinant c-Fos (fig. S2L). We found that the lower-affinity T206K mutant improved post-translational c-Fos degradation, whereas the higher-affinity T206E variant was associated with greater retention of c-Fos in purified proteasomes and induced less efficient degradation (Fig. 2, G to I; fig. S2M). These findings indicate that Protea-Tac degradation efficiency is not simply proportional to tAb binding affinity and that maximal affinity may not necessarily optimal for productive target turnover.

Because Protea-Tac relies on direct substrate localization to the 26S proteasome rather than ubiquitination, we next sought to investigate its dependency on substrate ubiquitination and proteasomal degradation. The induced degradation of c-Fos and EGFP-c-Fos by PSMD4- or ADRM1-based Protea-Tacs was strongly abrogated by MG132 (a proteasome inhibitor), but not by bafilomycin A1 (an autophagic flux inhibitor) or MLN7243 (an E1 Ub-activating enzyme inhibitor) (Fig. 3, A and B; fig. S3, A to D). Similarly, VHH(GFP)-ADRM1-mediated degradation of GFP-ODC was sensitive to MG132 but not to MLN7243 (Fig. 3, C and D). In chase experiments, treatment with MLN7243 had little effect on EGFP-c-Fos turnover by Protea-Tac but MG132 quickly halted it (Fig. 3, E to H). Flow cytometric analysis of EGFP-c-Fos stable cells further showed that MG132, but not MLN7243, attenuated Protea-Tac-mediated target degradation (fig. S3E). Importantly, both PSMD4- and ADRM1-based Protea-Tacs substantially reduced the protein levels of a lysine-less c-Fos mutant (c-Fos (K0)) without altering its mRNA levels, further supporting a ubiquitination-independent mode of target degradation (fig. S3, F to H). Together, these findings support a proteasome-dependent, ubiquitination-independent mode of Protea-Tac-mediated degradation.

**Fig. 3.**
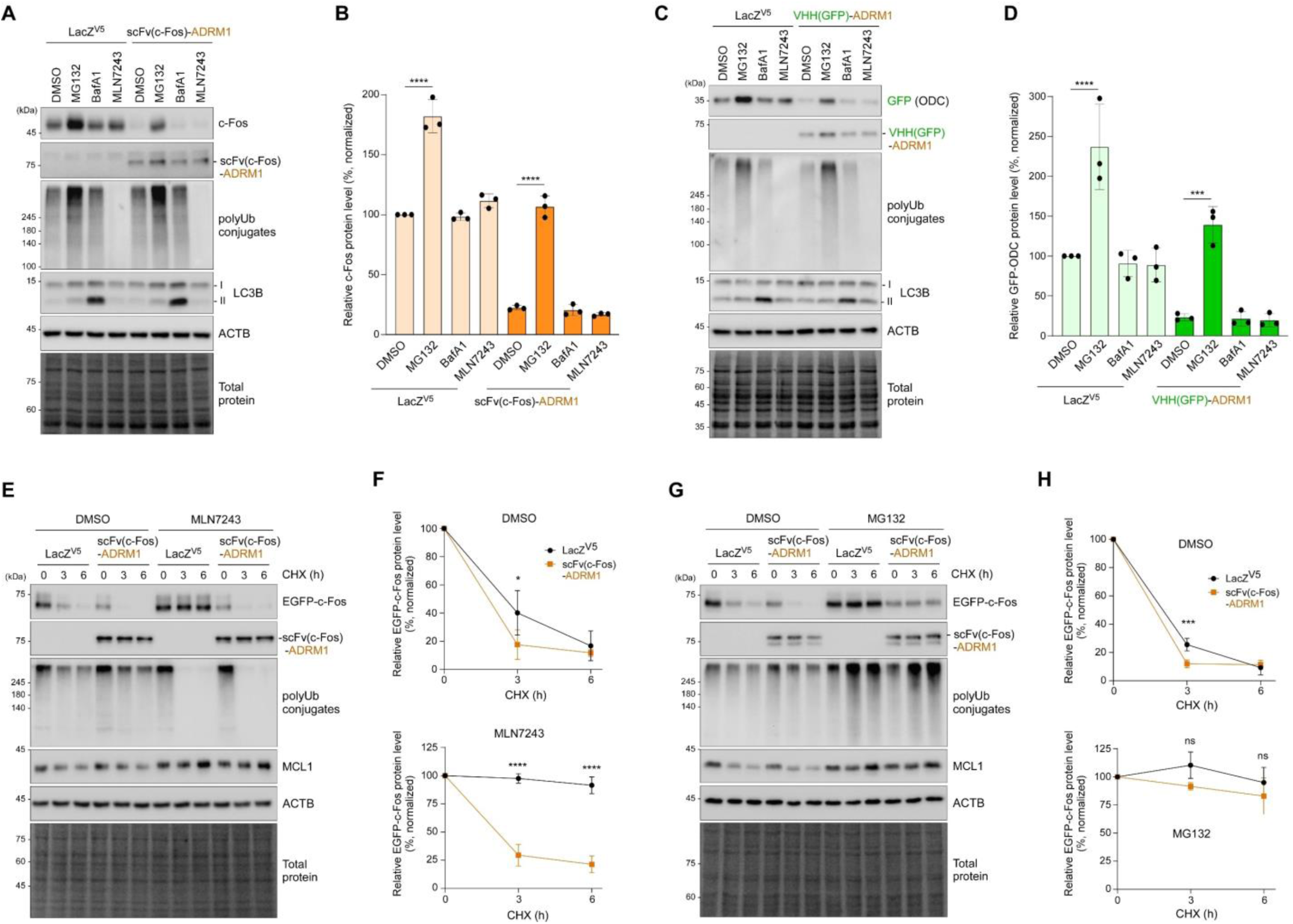
The Protea-Tac system enables Ub-independent and proteasome-mediated TPD. (**A**) scFv(c-Fos)-ADRM1-mediated c-Fos degradation was examined in the presence of DMSO, the proteasome inhibitor MG132 (10 μM), the vacuolar type H^+^-ATPase inhibitor bafilomycin A1 (BafA1; 200 nM), or the E1 Ub-activating enzyme inhibitor MLN7243 (1 μM) for 6 h. (**B**) The steady-state levels of c-Fos detected across conditions in (A) were quantified and normalized to total protein. Each bar indicates the mean ± SD (N = 3). ****p < 0.0001 (One-way ANOVA followed by Tukey’s multiple comparison test). (**C and D**) As in (A) and (B), except that VHH(GFP)-ADRM1 was used as a degrader and GFP-ODC was a neo-substrate. (**E**) Chase experiments were performed to track the degradation kinetics of EGFP-tagged version of c-Fos (EGFP-c-Fos) in the presence of DMSO or MLN7243. (**F**) Quantification of EGFP-c-Fos band intensities in (E), normalized to those of β-actin (ACTB) and to the values at 0 h post-CHX treatment (set as 100%), under DMSO (*top)* or MLN7243 (*bottom*) treatment conditions. Average percentages of remaining EGFP-c-Fos (mean ± SD) from three independent experiments (N = 3); *p < 0.05, ****p < 0.0001 (two-way ANOVA followed by Sidak’s post hoc test). (**G and H**) Chase experiments and quantification using EGFP-c-Fos as in panel (E) and (F) except they were performed in the presence of either DMSO or MG132. Data represent the mean ± SD from three independent experiments (N = 3). ***p < 0.001 (two-way ANOVA followed by Sidak’s post hoc test). ns, not significant. *See* Supplementary Figure 3.

### Versatility and specificity of modular Protea-Tac degraders

To assess the versatility of the Protea-Tac system in targeting diverse neo-substrates, as well as its modular flexibility allowing component substitution, we first replaced the tAb module with scFvs or VHHs that recognize common epitopes such as Flag, HA, or GFP tags (Fig. 4A). When expressed in A549 cells, both PSMD4-scFv(Flag) and scFv(HA)-ADRM1 degraders effectively reduced the levels of co-expressed Flag-TDP43 and HA-Tau proteins, respectively, in both the RIPA-soluble and RIPA-insoluble fractions, while there was no change in their mRNA levels (Fig. 4, B to E; fig. S4, A and B). To assess the effects of Protea-Tac on distinct biochemical pools of tau, we analyzed HEK293 cells stably expressing HA–tau (0N4R) by RIPA fractionation. While scFv(HA)-ADRM1 expression substantially reduced RIPA-soluble monomeric tau levels, it had little effect on high-molecular-weight tau (HMW) and insoluble species. These results suggest that Protea-Tac primarily reduces the soluble, monomeric tau pool under the conditions tested, rather than directly clearing soluble HMW or RIPA-insoluble tau species (fig. S4, C and D). In addition, a VHH-based chimera in which VHH(GFP) was fused to PSMD4 showed robust TPD activity toward GFP-tagged ornithine decarboxylase (ODC) (Fig. 4, F and G). Next, we set out to target BRD4, a protein whose inhibition or depletion has been shown to be therapeutically relevant as an anti-cancer intervention. The ADRM1-scFv(BRD4) chimera substantially lowered endogenous BRD4 protein levels without affecting *BRD4* mRNA expression, accelerating BRD4 turnover as well (Fig. 4, H and I; fig. S4F). Collectively, our results indicate that Protea-Tac degraders consistently elicit efficient target degradation by fusing intracellular tAbs to PSMD4 or ADRM1, with varied efficacy depending on target and module. Furthermore, the system displayed high substrate specificity from immunoblotting analyses: scFv(c-Fos) degraders did not affect the protein levels of Flag-TDP43, HA-Tau, or GFP-ODC, whereas scFv(Flag) or scFv(HA) degraders did not alter c-Fos levels (Fig. 4, J and K; fig. S2, G and H; fig. S4E).

**Fig. 4.**
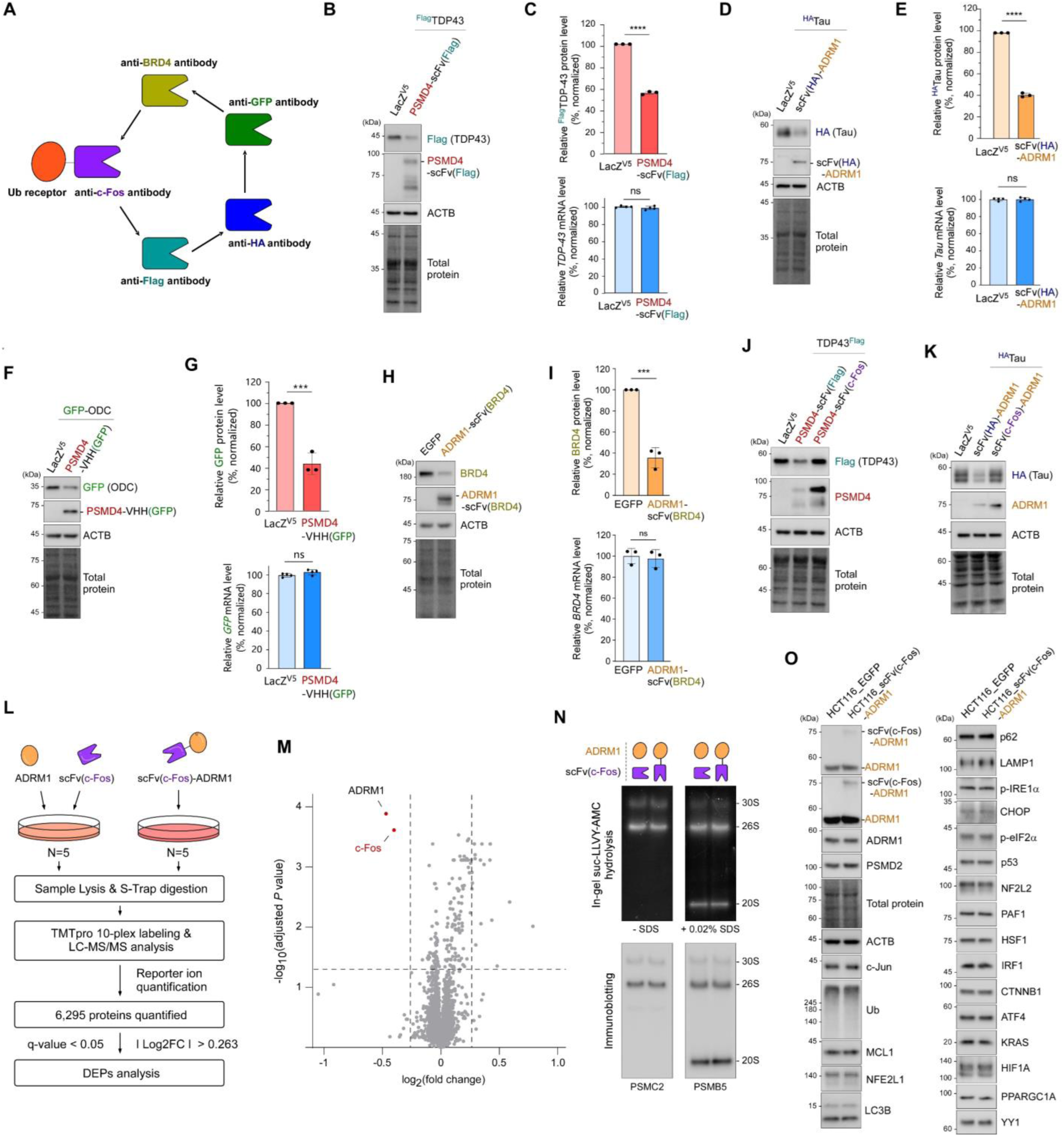
Protea-Tac-mediated degradation is highly selective and modular via UbR and tAb exchanges. (**A**) Schematic illustrating the modular design of Protea-Tac degraders with interchangeable tAb units. (**B**) A549 cells were co-transfected with ^Flag^TDP43 and either LacZ^V5^ or the Flag degrader (PSMD4-scFv(Flag)). After 72 h, WCLs were examined using SDS-PAGE/IB, followed by total protein staining. Representative results from three independent experiments are shown. (**C**) *Top*: quantification of ^Flag^TDP43 protein levels normalization to total protein signals. Bars indicate the mean ± SD (N = 3). ****p < 0.0001 (two-tailed Student’s *t*-test). *Bottom*: *^Flag^TDP43* mRNA levels were measured using qRT-PCR and normalized to *GAPDH.* ns, not significant (N = 4, two-tailed Student’s *t*-test). (**D** & **E**) Same as (B & C), except using ^HA^Tau as the target and scFv(HA)-ADRM1 as the degrader. (**F** & **G**) As in (B & C), except that a nanobody (VHH)-conjugated PSMD4 chimera (PSMD4-VHH(GFP)) was evaluated for degradation of GFP-tagged ODC. (**H**) A549 cells were treated with EGFP or anti-BRD4 Protea-Tac (ADRM1-scFv(BRD4)^HB^) encoding lenti virus for 72 h. WCLs were examined using SDS-PAGE/IB, followed by total protein staining. (**I**) *Top*: quantification of endogenous BRD4 protein levels normalized to total protein signals. Bars indicate the mean ± SD (N = 3). ***p < 0.001 (two-tailed Student’s *t*-test). *Bottom*: endogenous *BRD4* mRNA levels were measured using qRT-PCR and normalized to *GAPDH.* ns, not significant (N = 3, two-tailed Student’s *t*-test). (**J** & **K**) Specificity of Protea-Tac was assessed using mismatched target-degrader pairs. c-Fos-directed Protea-Tac showed no TPD activity toward TDP43^Flag^ (J) and ^HA^Tau (K). (**L**) Overview of the TMT-LC-MS/MS method for profiling global proteomes. Out of 6,295 identified proteins, 16 differentially expressed proteins (DEPs; 2 downregulated, 14 upregulated) were found in A549 cells stably expressing c-Fos treated with ADRM1-scFv(c-Fos) versus controls (ADRM1 + scFv(c-Fos)), using cutoffs of |log₂FC| > 0.263 and q < 0.05. (**M**) Volcano plot of DEPs with dotted lines representing fold-change (log₂) and p-value thresholds. Significantly downregulated DEPs (i.e., c-Fos and ADRM1) are marked in red; upregulated and non-DEPs in gray. (**N**) Native PAGE analysis of 26S proteasomes, followed by in-gel suc-LLVY-AMC hydrolysis visualization (*top*) and subsequent IB with antibodies against the CP (PSMB5) and RP (PSMC2) (*bottom*). The addition of 0.02% SDS activated 20S proteasomes. A549 cells were transfected with either scFv(c-Fos)-ADRM1 plasmid (4 μg) or individual scFv(c-Fos) and ADRM1 plasmids (2 μg each), as in (L). (**O**) Levels of various endogenous proteins in HCT116_EGFP and HCT116_scFv(c-Fos)-ADRM1 stable cell lines were compared using SDS-PAGE/IB. *See* Supplementary Figure 4.

To evaluate the global impact of Protea-Tac degraders, we performed a quantitative proteomic study using tandem mass tag (TMT)-based quantitative proteomics. Protein samples from five independent cultures of A549 cells overexpressing c-Fos transfected with scFv(c-Fos)-ADRM1 or individual modules (free ADRM1 and scFv(c-Fos) alone) were labeled with isobaric TMT reagents, pooled, fractionated via high-pH reverse-phase HPLC, and analyzed by LC-MS/MS methods, identifying a total of 6,295 proteins (Fig. 4L). The Principal component analysis showed separation between the control and Protea-Tac groups (fig. S4H). Applying fold-change thresholds of >1.2 or <0.83 and a q-value of <0.05, only c-Fos and ADRM1 were significantly reduced, while the broader proteome remained largely unchanged (Fig. 4M). With the same criteria, 14 proteins (0.22%) were identified to be upregulated but could not be enriched within any hallmark or ontology gene set. The pronounced reduction in c-Fos protein levels, as confirmed by both biochemical and TMT-MS assays, underscores high selectivity of the Protea-Tac system, driven by tAb specificity. The observed reduction in ADRM1 abundance likely reflects the faster turnover of the scFv(c-Fos)-ADRM1 fusion compared with ADRM1 expressed separately with scFv(c-Fos). Additionally, the absence of detectable changes in cyclin D1 and other downstream proteins likely reflects the artificial c-Fos-overexpression context used for TMT-MS/MS to ensure reliable quantification of on-target c-Fos depletion (fig. S4G). Even after stable overexpression of Protea-Tac chimeras, we detected no marked changes in native proteasome profiles, in-gel proteasome activity, proteasome abundance, or selected markers of ubiquitin-proteasome system, autophagy–lysosomal pathways, and ER stress (Fig. 4, N and O; fig. S4, I to K). Taken together, these results indicated that modifying proteasomal UbRs with tAb is not only well-tolerable by both the 26S proteasome and the global proteome but also supports the feasibility of selective target degradation with limited broader proteome perturbation under the conditions tested.

### *In vivo* target degradation and anti-tumor efficacy of Protea-Tac

To assess whether Protea-Tac-mediated c-Fos depletion suppresses tumor growth in vivo, we generated xenografts using stable MCF7 and HCT116 cell lines overexpressing scFv(c-Fos)-ADRM1 and their corresponding EGFP-expressing controls (Fig. 1J; fig. S1N), and subcutaneously injected equivalent quantities of these stable cells into six-week-old male NOD/SCID/IL-2γ-receptor-null (NSG) mice. Tumor volumes and body weights were monitored every three days for five to seven weeks. At the experimental endpoint (days 32 or 50), mice with Protea-Tac-expressing tumors exhibited substantially delayed tumor growth than controls (978.1 ± 285.5 mm^3^ in MCF7_EGFP tumors vs. 92.4 ± 18.9 mm^3^ in MCF7_scFv(c-Fos)-ADRM1 tumors; 1244.8 ± 459.8 mm^3^ in HCT116_EGFP tumors vs. 176.3 ± 44.8 mm^3^ in HCT116_scFv(c-Fos)-ADRM1 tumors) (Fig. 5, A and B). In stark contrast, there were no significant differences in body weights between the groups with MCF7 and HCT116 tumors (Fig. 5C; fig. S5A).

**Fig. 5.**
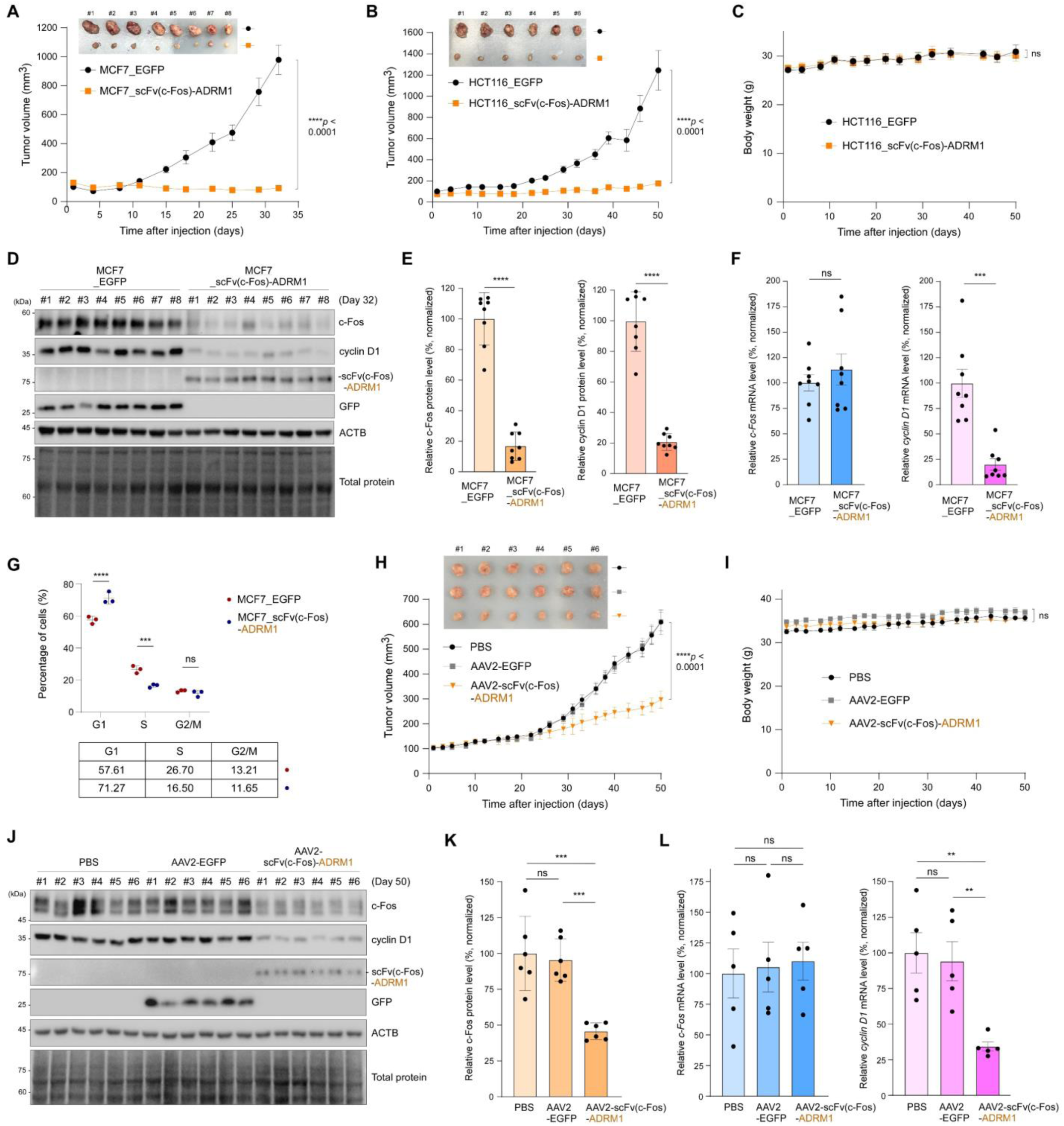
Protea-Tac-induced c-Fos degradation leads to anti-tumor efficacy *in vivo*. (**A**) After subcutaneous inoculation of 5 × 10^6^ MCF7 cells stably expressing either EGFP or scFv(c-Fos)-ADRM1 into six-week-old male NOD/SCID/IL-2γ-receptor null (NSG) mice, tumor volumes were measured twice a week for approximately 5 weeks post-implantation. Data are presented as means ± SEM (N = 8 per group); Mann-Whitney *U* test (*p* < 0.0001). *Insets* are image of tumors harvested (day 32). (**B**) As in (A), Tumors harvested on day 50 from HCT116_EGFP or HCT116_scFv(c-Fos)-ADRM1 xenografts. (**C**) Body-weight changes during the HCT116 xenograft experiment. (**D**) IB analysis of xenografted tumor lysates showing effective *in vivo* degradation of endogenous c-Fos by the Protea-Tac degrader. (**E**) Quantification of c-Fos (*left*) and cyclin D1 (*right*) protein levels, normalized to total protein signals. Bars represent the mean ± SD (N = 8). \*\*\*\**p* < 0.0001 (two-tailed Student’s t test). (**F**) As in (E), but mRNA levels of *c-Fos* and *cyclin D1* were determined using qRT-PCR and normalized to *GAPDH*. Bars represent the mean ± SEM (N = 8). \*\*\**p* < 0.001 (two-tailed Student’s t test). ns, not significant. (**G**) Cell cycle distribution in the MCF7 stable cell lines used for xenograft assays, analyzed by flow cytometry. DNA contents were quantified using DAPI staining. \*\*\**p* < 0.001, \*\*\*\**p* < 0.0001 between EGFP-and Protea-Tac-expressing tumors, based on two-way ANOVA followed by Bonferroni’s multiple-comparisons test (N = 3 each). ns, not significant. (**H**) MCF7-xenografted NSG mice were intratumorally injected with PBS, AAV2-EGFP, or AAV2-scFv(c-Fos)-ADRM1 (2 × 10^9^ vg each) three times per week, and tumor growth curves were measured three times per week for 50 days. Data are presented as mean ± SEM (N = 6 for each group); two-way ANOVA followed by Tukey’s multiple-comparisons test (\*\*\*\**p* < 0.0001). (**I**) Body weight changes during the AAV2-delivery experiment. (**J**) IB analysis of whole-tumor lysates with indicated antibodies. (**K**) Quantification of c-Fos protein levels, normalized to total protein signals. Bars represent mean ± SD (N = 6). ***p < 0.001 (one-way ANOVA with Tukey’s post hoc test). ns, not significant. (**L**) Quantitative RT-PCR analysis of c-Fos and cyclin D1 mRNA levels. Bars represent mean ± SEM (N = 5). **p < 0.01 (one-way ANOVA with Tukey’s post hoc test). *See* Supplementary Figures 5 & 6.

Immunoblotting examination of dissected tumor lysates revealed marked depletion of c-Fos protein in scFv(c-Fos)-ADRM1-expressing tumors but not in EGFP-expressing controls (Fig. 5D; fig. S5B), demonstrating *in vivo* TPD activity of the Protea-Tac system. Furthermore, cyclin D1 was also significantly reduced, but there were little change in PARP cleavage (an apoptotic marker) and ERK1/2 and p38 phosphorylation (c-Fos upstream markers) (Fig. 5E; fig. S5, C to E). Analysis of mRNA levels from the tumor tissues revealed a significant downregulation of cyclin D1 expression, a marked upregulation of Protea-Tac mRNA expression, and no change in endogenous c-Fos mRNA levels (Fig. 5F; fig. S5, F and G). Flow cytometry of stable cell lines used in xenograft assays further confirmed a pronounced delay in cell cycle progression, particularly at the G1–S transition, in tumors expressing the c-Fos-targeting Protea-Tac degraders (Fig. 5G; fig. S5H). These findings demonstrate that Protea-Tac degraders have substantial anti-tumor actions *in vivo*, which are mediated predominantly through cell cycle arrest rather than apoptosis activation.

To evaluate the therapeutic potential of Protea-Tac, we used adeno-associated virus (AAV)-based expression and MCF7 xenografts in NSG mice. Intratumoral injection with AAV2 encoding either EGFP or scFv(c-Fos)-ADRM1 revealed that mice receiving Protea-Tac exhibited a significant delay in tumor progression compared to both PBS- and AAV2-EGFP-injected controls (608.5 ± 121.9 mm^3^ for PBS vs. 610.2 ± 92.4 mm^3^ in AAV2-EGFP vs. 296.9 ± 34.1 mm^3^ for AAV2-scFv(c-Fos)-ADRM1; \*\*\*\**p* < 0.0001) (Fig. 5H). Body weights remained comparable across all groups, with no evident treatment-associated weight loss (Fig. 5I). IB analyses of tumor lysates mirrored the molecular profiles observed in stable cell-derived xenografts, showing robust expression of AAV2-scFv(c-Fos)-ADRM1, efficient degradation of c-Fos, and a marked reduction in cyclin D1 protein levels (Fig. 5, J and K; fig. S5I). Unlike *c-Fos* mRNA, which was unchanged in the AAV2-Protea-Tac group, *cyclin D1* mRNA expression was markedly reduced (Fig. 5L).

To further evaluate AAV2-mediated Protea-Tac delivery in an immunocompetent setting, we established a syngeneic mouse tumor model by subcutaneously implanting CT26 cells into BALB/c mice. When tumors reached approximately 150 mm³, mice were intratumorally administered PBS or AAV2 encoding EGFP or scFv(c-Fos)-ADRM1 once weekly for a total of three injections. AAV2-mediated expression of scFv(c-Fos)-ADRM1 significantly suppressed tumor growth compared with the PBS- and AAV2-EGFP-treated groups (fig. S6, A and B). Immunoblotting analyses of tumor lysates showed reduced c-Fos and cyclin D1 protein levels in the AAV2-scFv(c-Fos)-ADRM1 group (fig. S6, D and E). Body weights remained comparable across the groups, and serum biochemical analyses revealed no significant differences in ALT, AST, ALP, UA, creatinine, or BUN levels at the experimental endpoint (fig. S6, C, F, and G). Serum IL-6 levels were also not significantly elevated at the endpoint (fig. S6H).

### Non-viral mRNA-LNP delivery of Protea-Tac induces target degradation and suppresses tumor growth *in vivo*

To further explore delivery strategies beyond AAV-based expression, we next evaluated whether Protea-Tac could be delivered as mRNA formulated in lipid nanoparticles (LNPs). Although ADRM1-based Protea-Tacs were used in our preceding *in vivo* studies, PSMD4-based Protea-Tac also exhibited robust degradation activity and was therefore selected for evaluation in the context of mRNA delivery. Consistent with our earlier findings, stable expression of PSMD4-scFv(c-Fos), but not scFv(c-Fos) alone, reduced endogenous c-Fos protein levels in HCT116 cells without altering c-Fos mRNA levels. This reduction was accompanied by decreased cyclin D1 protein and mRNA levels (fig. S7, A and B). Similarly, transfection of A549 cells with mRNA encoding PSMD4-scFv(c-Fos) reduced endogenous c-Fos protein levels while leaving c-Fos mRNA levels unchanged. Cyclin D1 expression was also reduced under these conditions (fig. S7, C to E).

We next evaluated mRNA-LNP-mediated Protea-Tac delivery in an A549 xenograft model. Mice bearing A549 tumors received subcutaneous injections of PBS or LNP formulations containing mRNA encoding EGFP, scFv(c-Fos), or PSMD4-scFv(c-Fos) at the indicated time points. Mice treated with LNP formulations containing mRNA encoding PSMD4-scFv(c-Fos) exhibited significantly delayed tumor growth compared with mice receiving PBS or LNP formulations containing mRNA encoding EGFP or scFv(c-Fos) (Fig. 6, A and B).

**Fig. 6.**
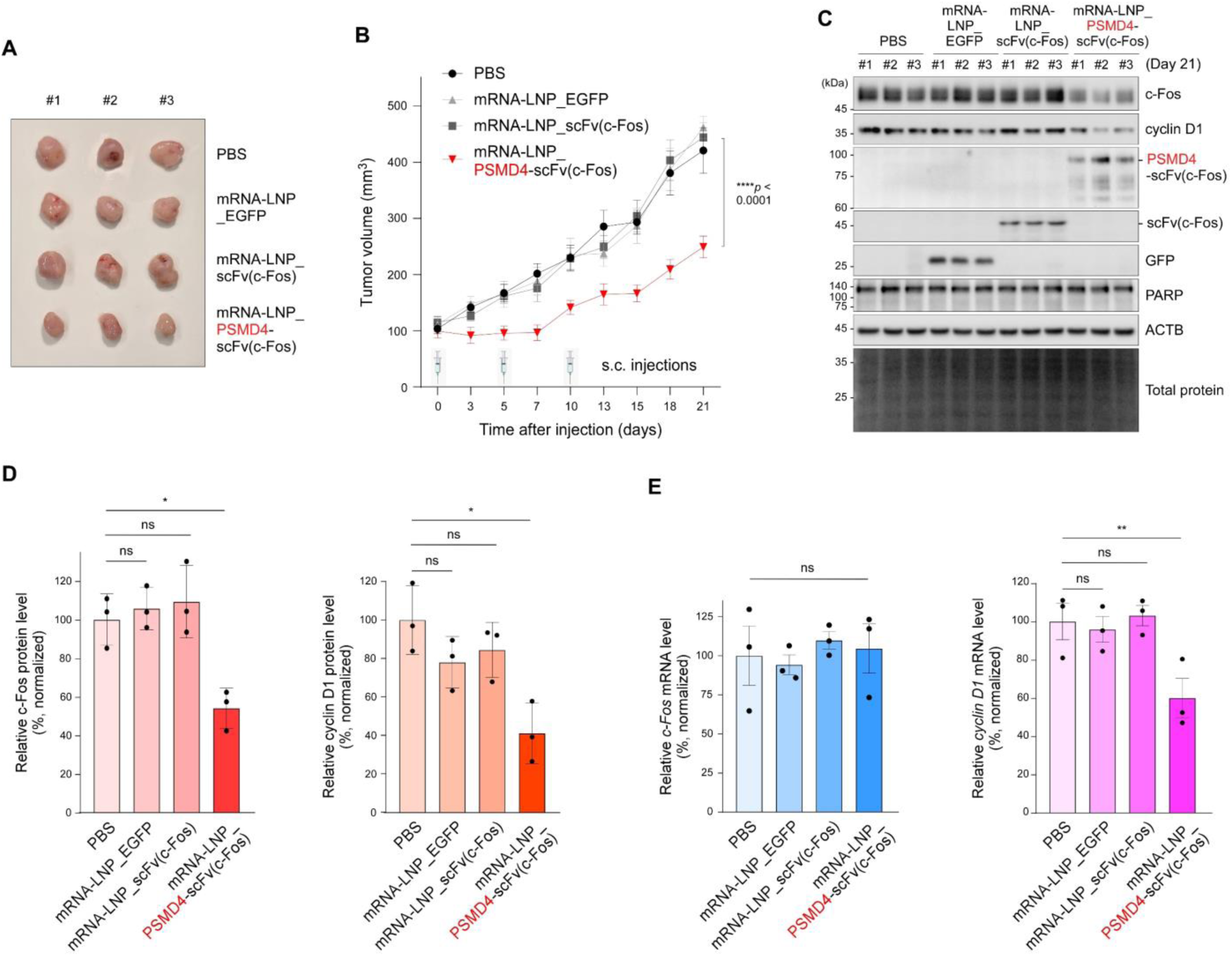
mRNA-LNP-mediated Protea-Tac expression inhibits tumor growth *in vivo*. (**A**) A549-xenografted 6-week-old female BALB/cAnN-Foxn1nu/Crl mice received subcutaneous injections of PBS or LNP formulations containing mRNA encoding EGFP, scFv(c-Fos), or PSMD4-scFv(c-Fos). Representative image of tumors harvested is shown. (**B**) Tumor volumes were measured twice a week for 21 days. LNP formulations were administered subcutaneously every 5 days for a total of three doses. Each formulation contained 40 μg of mRNA per dose. Data are presented as means ± SEM (N = 6 per group); two-way ANOVA followed by Tukey’s multiple-comparisons test (p < 0.0001). (**C**) IB analysis of whole-tumor lysates with indicated antibodies. (**D**) Quantification of c-Fos (*left*) and cyclin D1 (*right*) protein levels, normalized to total protein signals. Bars represent the mean ± SD (N = 3). ns, not significant, \**p* < 0.05 (one-way ANOVA with Tukey’s post hoc test). (**E**) As in (D), but mRNA levels of *c-Fos* and *cyclin D1* were determined using qRT-PCR and normalized to *GAPDH*. Bars represent the mean ± SEM (N = 3). ns, not significant, \*\**p* < 0.01 (one-way ANOVA with Tukey’s post hoc test).

Immunoblotting analyses of tumor lysates confirmed expression of the encoded proteins and revealed marked reductions in c-Fos and cyclin D1 protein levels specifically in tumors treated with PSMD4-scFv(c-Fos) encoding mRNA-LNPs, without evident PARP cleavage (Fig. 6, C and D). Analysis of tumor mRNA levels showed that c-Fos mRNA remained unchanged, whereas cyclin D1 mRNA was significantly reduced (Fig. 6E). These results are consistent with post-translational depletion of c-Fos and the consequent suppression of cyclin D1 expression. Serum biochemical analyses revealed no significant differences in alanine aminotransferase (ALT), aspartate aminotransferase (AST), blood urea nitrogen (BUN), or creatinine (CREA) levels among all groups (fig. S7F), indicating no detectable hepatic or renal toxicity under the conditions tested. Together with our AAV-mediated experiments, these findings provide proof-of-concept that Protea-Tac can be delivered through both viral and non-viral modalities to show anti-tumor efficacy *in vivo*.

## Discussion

The 26S proteasome is the sole ATP-dependent protease in the eukaryotic cytoplasm and nucleus, which uses Ub chains for the recognition of to-be-degraded substrates. Ub chains are primarily used to localize substrates to the 26S proteasome. Unless a substrate lacks a flexible, unstructured region (the “initiation site”) that triggers unfolding and translation via the AAA+ PSMC pore loop, this tethering alone is usually sufficient to cause spontaneous substrate hydrolysis in the 20S core particle (*63–65*). Based on this mechanism, we developed Protea-Tac by fusing a proteasomal Ub receptor to an intracellular targeting antibody, thereby replacing Ub-chain-mediated substrate recruitment with direct antibody-guided engagement of the 26S proteasome. This modular architecture enabled degradation of endogenous c-Fos and BRD4 and further supported generalizability of the platform in assays using cognate tag-directed chimeras against tagged TDP-43, tau, and ODC model substrates. Its activity against endogenous c-Fos and the comparatively stable endogenous BRD4 further indicates that Protea-Tac is not restricted to rapidly turned-over substrates. By promoting Ub-independent proteasomal degradation, Protea-Tac bypasses the substrate-ubiquitination and E3-ligase-recruitment steps required by conventional ubiquitination-inducing degraders, including PROTACs and many MGDs. This feature may be advantageous in settings where the engaged ubiquitin-transfer machinery is compromised, a recognized mechanism of resistance to conventional degraders (*66, 67*).

Recent direct-to-proteasome small-molecule degraders have established that chemically induced recruitment of neo-substrates to UbRs or other RP components can induce Ub-independent TPD (*50–54*). These approaches offer important advantages inherent to small-molecule modalities, including cell permeability and opportunities for pharmacological optimization. Protea-Tac should therefore be viewed as a complementary protein-based strategy rather than a competing replacement. Its distinction lies not simply in its molecular format, but in the source of target recognition: whereas chemical degraders depend on the availability of a suitable small-molecule ligand for each neo-substrate, Protea-Tac encodes target recognition in a modular intracellular binder. This does not eliminate the need for optimization, but shifts an important early bottleneck from small-molecule warhead discovery toward the identification and engineering of binders with suitable specificity and geometry. Such a framework may be particularly useful for ligand-poor proteins and for applications in which epitope- or conformation-selective engagement is desirable (*68–71*).

Beyond expanding the range of accessible targets, the proteasome-centered design of Protea-Tac may offer additional practical advantages. Protea-Tac may benefit from the substantial cellular abundance of 26S proteasomes, as robust target degradation was observed despite the apparent remodeling of only a minor subpopulation of cellular proteasomes (*24, 25*). This limited extent of remodeling may help preserve global proteasome activity and minimize nonspecific perturbation of endogenous protein turnover, while precise antibody-guided recognition contributes to the observed target selectivity. Our affinity-tuning analyses further suggest that maximal substrate-binding affinity is not necessarily required for efficient Protea-Tac activity; rather, a dynamic interaction with the recruited substrate may favor productive turnover in permissive configurations. Together with ongoing advances in antibody and de novo mini-binder engineering, the modularity of this platform may facilitate the development of a broader repertoire of Protea-Tac degraders for disease-associated proteins that remain difficult to engage with small-molecule ligands (*72*).

At the same time, the modularity of Protea-Tac does not imply that every configuration will be equally productive. Protea-Tac activity was also dependent on the proteasomal subunit used to construct the chimera. Whereas PSMD4- and ADRM1-based Protea-Tacs efficiently reduced c-Fos, chimeras constructed with PSMD2, PSMA4, PSMB2, or PSMC5 did not induce appreciable target degradation. Moreover, extending the flexible linker did not restore degradation activity in the inactive configurations tested. Although the structural basis for these differences remains to be determined, these findings are consistent with the concept that recruitment to the proteasome is necessary but not invariably sufficient for productive degradation (*50*). Proteasomal processing requires an accessible initiation region that can engage the AAA+ ATPase pore, and the spatial relationship between a proteasome-binding element and the initiation region can influence whether a recruited substrate is efficiently degraded (*73*). Substrate engagement is then coupled to conformational transitions of the 26S proteasome that support unfolding and translocation into the 20S core particle (*74*). Thus, the efficiency of a Protea-Tac degrader may depend on how the recruited substrate is positioned relative to the proteasome. A productive subunit–linker configuration may be required for the substrate engagement to the ATPase pore and subsequently undergo processive translocation. Because the position and accessibility of initiation regions are likely to vary among neo-substrates, future development of Protea-Tac may benefit from substrate-specific, structure-guided optimization of the proteasomal subunit, linker architecture, and targeting binder.

Despite inherent challenges related to intracellular delivery and expression, Protea-Tac represents a distinct and promising protein-based TPD strategy. Our in vivo studies further demonstrate that this platform can be implemented using both viral and non-viral delivery formats. Using AAV2-mediated intratumoral delivery, we observed target degradation and anti-tumor efficacy in both an MCF7 xenograft model and a CT26 syngeneic mouse tumor model, indicating that Protea-Tac can remain active in an immunocompetent setting. AAV vectors have become an established platform for clinical gene delivery, with multiple recombinant AAV-based products receiving regulatory approval (*75*). In parallel, subcutaneous administration of mRNA-LNPs encoding PSMD4-scFv(c-Fos) reduced c-Fos protein levels and attenuated tumor growth in A549 xenografts. Given the clinical validation of LNPs as vehicles for mRNA delivery, these findings provide proof of concept for non-viral delivery of Protea-Tac-encoding mRNA (*76*). Meanwhile, the immunogenicity of the delivery vehicles and the Protea-Tac protein itself will also require dedicated investigation, including humoral and cellular responses and the consequences of repeated administration. This consideration is particularly important because recombinant AAV vectors can elicit innate and adaptive immune responses, while LNP composition can influence immune activation and the efficacy of mRNA therapeutics (*77, 78*). Thus, the present study does not establish a clinically optimized therapeutic modality; rather, it defines a modular framework for direct, antibody-guided, Ub-independent degradation through engineered 26S proteasomes and provides in vivo proof of concept using both viral and non-viral delivery formats. Its mechanistic basis, target selectivity, and modularity support further development of Protea-Tac as a complementary strategy for expanding the range of proteins accessible to TPD.

## Material and Methods

### Antibodies and Reagents

The following primary antibodies were used: anti-c-Fos (Cell Signaling Technology [CST], 1:2,000 for IB), anti-PSMD4 (Thermo Fisher Scientific [TFS], 1:1,000), anti-ADRM1 (TFS, 1:1,000), anti-PSMA4 (Enzo Life Sciences, 1:1,000), anti-V5 (TFS, 1:3,000), anti-ACTB (MilliporeSigma, 1:3,000), anti-GAPDH (Santa Cruz Biotechnology [SCBT], 1:3,000), anti-MCL1 (CST, 1:2,000), Streptavidin-HRP (MilliporeSigma, 1:1,000), anti-cyclin D1 (CST, 1:2,000), anti-His (CST, 1:1,000), anti-VCP (TFS, 1:3,000), anti-Flag (TFS, 1:1,000), anti-HA (MilliporeSigma, 1:3,000), anti-GFP (SCBT, 1:3,000), anti-PSMC2 (SCBT, 1:1,000), anti-PSMB5 (TFS, 1:1,000), anti-PSMB6 (TFS, 1:1,000), anti-PSMA3 (Enzo Life Sciences, 1:1,000), anti-PSMD1 (SCBT, 1:1,000), anti-LC3B (MilliporeSigma, 1:1,000), anti-Ub (SCBT, 1:3,000), anti-NFE2L1 (TFS, PA5-90023, 1:3,000), anti-p62 (Abcam, ab56416, 1:3,000), anti-LAMP1 (SCBT, sc-20011, 1:1,000), anti-p-IRE1α (Ser724, TFS, PA1-16927, 1:1,000), anti-CHOP (TFS, MA1-250, 1:1,000), anti-p-eIF2a (Abclonal, AP0342, 1:1,000), anti-c-Jun (CST, 9165T, 1:2,000), anti-p53 (TFS, MA5-12557, 1:1000), anti-PAF1 (Abnova, 551-650, 1:1000), anti-HSF1 (CST, 12972S, 1:1000), anti-IRF-1 (CST, 8478T, 1:1000), anti-CTNNB1 (CST, 8480S, 1:1000), anti-ATF4 (TFS, PA5-19521, 1:1000), anti-RAS (CST, 3965, 1:1000), anti-HIF-1 (Novus, NB100-105, 1:1,000), anti-YY1 (Abclonal, A19569, 1:1,000), anti-NFE2L2 (abcam, ab62352, 1:1000), anti-PPARGC1A (Abclonal, A12348, 1:1000), anti-PARP (TFS, 436400, 1:1,000), anti-ERK1/2 (CST, 4695S, 1:1,000), anti-p-ERK1/2 (CST, 4370S, 1:1,000), anti-p38 (CST, 8690T, 1:1,000), and anti-p-p38 (CST, 4511T, 1:1,000). Secondary antibodies included HRP-conjugated anti-mouse IgG, anti-rabbit IgG, and anti-rat IgG (all from TFS, used at 1:10,000 for IB). Sources of major biochemical reagents are as follows: MG132 (AG Scientific), MLN7243 (Cayman Chemical), Bafilomycin A1 (Cayman Chemical), suc-LLVY-AMC (Bachem), and No-Stain™ Protein Labeling Reagent (TFS). All scFvs and VHHs were derived from plasmids deposited at Addgene (182090, 198303, 198302, and 166518).

### Mammalian cell culture and transient expression

Mammalian cell lines used in this study, including A549 (KCLB No : 10185), HCT116 (KCLB No : 10247), MCF7 (KCLB No : 30022), and their derivatives stably expressing EGFP, scFv(c-Fos)-ADRM1, were cultured in RPMI (Wellgene), whereas HEK293T cells were cultured in DMEM (Wellgene), supplemented with 10% heat-inactivated FBS (Gibco), 100 U/mL penicillin, 100 μg/mL streptomycin, and 2 mM L-glutamine. Cells were maintained at 37 °C in a humidified atmosphere containing 5% CO₂. For transient overexpression, cells were seeded to reach approximately 70% confluence and transfected with indicated plasmid DNA using Lipofectamine 3000 (TFS) according to the manufacturer’s instructions. The total amount of plasmid DNA was kept constant within each experimental set (2 or 4 μg of total plasmid DNA in a six-well plate). After incubation, cells were washed with PBS and lysed in RIPA buffer (50 mM Tris-HCl [pH 8.0], 1% NP-40, 0.5% deoxycholate, 0.1% sodium dodecyl sulfate [SDS], and 150 mM NaCl) supplemented with protease inhibitor cocktails, or other indicated buffers. WCLs were clarified by centrifugation at 13,000 rpm for 15 min at 4 °C, mixed with 2 × SDS sample buffer, and denatured at 85 °C for 15 min prior to SDS-PAGE and subsequent analyses. For chase analysis, cells were treated with 80 μg/mL cycloheximide at time zero, and samples were collected at indicated chase points.

### RNA isolation and quantitative RT-PCR (qRT-PCR)

Total RNA from cultured cells was extracted using TRIzol (Favorgen), followed by additional purification with RNeasy Mini Columns (Qiagen) with on-column DNase I treatment. cDNA synthesis was performed using an RT-PCR premix (Bioneer) and qRT-PCR was conducted using 1/20-diluted cDNA, SYBR qPCR Master Mix (Bioneer), and 10 µM primers. The target gene-specific primer sequences were as follows: for GAPDH, forward 5’-AGGGCCCTGACAACTCTTTT-3’ and reverse 5’-AGGGGTCTACATGGCAACTG-3’; for c-Fos, forward 5’-CACTCCAAGCGGAGACAGAC-3’ and reverse 5’-AGGTCATCAGGGATCTTGCAG-3’; for cyclin D1, forward 5’-GCTGCGAAGTGGAAACCATC-3’ and reverse 5’-CCTCCTTCTGCACACATTTGAA-3’; for c-Fos (K0), forward 5’-TACCCCTACGACGTGCCCG-3’ and reverse 5’-GAGTCTGCGGGTGAGTGGTAG -3’; for BRD4, forward 5’-ACCTCCAACCCTAACAAGCC-3’ and reverse 5’-TTTCCATAGTGTCTTGAGCACC-3’; for TDP-43, forward 5’-GGGTAACCGAAGATGAGAACG-3’ and reverse 5’-CTGGGCTGTAACCGTGGAG-3’; for tau, forward 5’-CCAAGTGTGGCTCATTAGGCA-3’ and reverse 5’-CCAATCTTCGACTGGACTCTGT-3’; for GFP, forward 5’-AAGCAGAAGAACGGCATCAA-3’ and reverse 5’-GGGGGTGTTCTGCTGGTAGT-3’; and for scFv(c-Fos)-ADRM1, forward 5’-CCGAAGATCTGGGAGTGTACT-3’ and reverse 5’-CACCAAGTACTTGTTGGAGGC-3’, where the scFv primers were designed to specifically amplify the scFv region, avoiding endogenous ADRM1. Gene expression levels were normalized to *GAPDH*, and statistical significance was evaluated using a two-tailed Student’s t-test with p < 0.05 considered significant.

### Soluble/insoluble fractionation

Cells expressing Protea-Tac degraders were washed with ice-cold PBS and lysed in RIPA buffer supplemented with protease inhibitor cocktails. WCLs were centrifuged at 16,000 × g for 30 min at 4 °C to separate supernatants (RIPA-soluble fractions), which were boiled at 85 °C for 15 min in 2 × SDS sample buffer. Pellets (RIPA-insoluble fractions) were washed with lysis buffer, resuspended in SDS sample buffer, and boiled at 100 °C for 15 min.

### Affinity purification and Native PAGE analysis of 26S proteasomes

For affinity purification using biotin tags, cells were harvested in lysis buffer (25 mM Tris-HCl [pH 7.5], 10% glycerol, 5 μM MgCl2, 1 mM ATP, 1 mM DTT, and protease inhibitor cocktails) and homogenized with a Dounce homogenizer. The lysates were incubated with streptavidin-conjugated agarose beads overnight at 4 °C. The beads were washed five times with wash buffer (20 mM Tris [pH 7.5], 15 % glycerol, 1 mM EDTA, 150 mM KCl, 0.05 % NP-40, 1 mM DTT, and 0.2 mM PMSF). Next, the precipitated proteins were eluted from the resin by incubating in TEV cleavage buffer (50 mM Tris-HCl [pH 7.5], 1 mM ATP, 10% glycerol, and TEV protease) for 3 h at 30 °C. Then, the eluted proteins were concentrated with Amicon Ultra-0.5 centrifugal filter units (MilliporeSigma). Proteins were resuspended in Tris-Glycine Native Sample Buffer (TFS) and resolved on 3–8% NuPAGE Tris-Acetate native gels (TFS) at 150 V for 4–5 h. Gels were incubated in activity assay buffer (20 mM Tris, 1 mM ATP, 5 mM MgCl₂) containing 100 μM suc-LLVY-AMC to visualize proteasome complexes. To activate 20S proteasomes in native gels, 0.02% SDS was added. After fluorescence-based in-gel activity assay, separated samples in the gel were transferred to PVDF membranes for IB analysis.

### Flow cytometry for c-Fos degradation and cell cycle distribution

EGFP-c-Fos stable cells were transfected with either tricistronic scFv(c-Fos)-T2A-ADRM1-P2A-mCherry or bicistronic scFv(c-Fos)-ADRM1-P2A-mCherry plasmids and incubated for 72 h. Cells were harvested, washed, and resuspended in FACS buffer (0.2 % of BSA, 0.1 % of sodium azide, and 2 mM of EDTA in PBS). Flow cytometry was performed on a BD LSRFortessa X-20 using FITC and PE channels. Data were analyzed using Flowjo software (ver. 10.9.0). For DNA content analysis, EGFP or scFv(c-Fos)-ADRM1 expressing cells were fixed in 70% ethanol overnight, washed, and stained with DAPI/Triton X-100 at RT for 30 min in the dark. Flow cytometry was performed using LSRFortessa (BD Bioscience) with UV excitation at 340 to 380 nm and analyzed using FlowJo to quantify the distribution across G0/G1, S, and G2/M phases.

### Affinity tuning of scFv(c-Fos) and affinity measurement

To change the binding affinity of scFv(c-Fos), random mutagenesis of six central residues within the CDR3 region was performed using primers containing NNK degenerate codons. The resulting mutant libraries were cloned into the pMopac12 vector and transformed into *E. coli* Jude1 cells. Individual colonies (N = 596) were cultured in 96-deep-well plates and induced with 0.5 mM IPTG at 20°C for 20 h. Secreted scFv(c-Fos) variants were purified from culture supernatants using Ni-NTA affinity chromatography. To express recombinant GST-tagged c-Fos, *E. coli* BL21(DE3) cells harboring the expression plasmids were induced with 0.5 mM IPTG, followed by incubation at 30°C for 16 h. Cells were lysed, and clarified lysates were applied to a GST-affinity column, eluted with 20 mM reduced glutathione, and buffer-exchanged into PBS using Amicon Ultra centrifugal filters (10 kDa MWCO). Protein purity and molecular weight were verified by SDS-PAGE and protein staining, and concentrations were determined by absorbance at 280 nm using a NanoDrop spectrophotometer.

To assess antibody affinity, purified scFvs (200 ng/well) were coated onto 96-well immunoplates and incubated overnight at 4°C. After blocking with 3% BSA in PBS, serial dilutions of GST-c-Fos (ranging from 5 µM to 0.32 nM) were added to the plates and incubated for 1 h at RT. Plates were then washed with PBST and incubated with HRP-conjugated anti-GST antibodies for additional 1 h at RT. TMB-ELISA substrates (TFS) were then applied for 20 min, and the reaction was stopped by the addition of 2 M H₂SO₄. Absorbance at 450 nm was measured using an Infinite 200 PRO NanoQuant microplate reader. Dose-response curves were generated, and half-maximal inhibitory concentrations (IC₅₀) were calculated from normalized values using GraphPad Prism (ver. 10.4.1).

### TMT-MS analysis

Protein samples were prepared from A549 cells after transfection with either ADRM1 and scFv(c-Fos) (control) or scFv(c-Fos)-ADRM1 degraders (Protea-Tac) for 64 h in biological quintuplicates (N = 5 per group). Cells were washed three times with ice-cold PBS, snap-frozen in liquid nitrogen, lysed in freshly prepared MS lysis buffer (4% SDS and 0.1 M Tris-HCl [pH 8.0]), heated at 95°C for 20 min, and sonicated. Lysates were cleared by centrifugation at 15,000 rpm for 20 min at room temperature (RT), and protein concentrations were determined using a BCA assay (TFS). An aliquot containing 40 µg of total protein per sample was precipitated in pre-chilled acetone at −20°C overnight and collected by centrifugation at 14,000 rpm for 10 min at 4°C. After air-drying, pellets were resuspended in denaturing buffer (5% SDS, 10 mM TCEP, 50 mM CAA, and 50 mM HEPES [pH 8.5]), reduced, alkylated, and then acidified with phosphoric acid to pH ≤1. Following additional centrifugation, the samples were processed with S-Trap micro columns (Protifi), washed with methanol-containing buffer, and digested on-column with trypsin (1:25 enzyme-to-protein ratio) at 47°C for 2 h. Peptides were eluted sequentially using three buffers (50 mM HEPES; 0.2% formic acid in water; and 0.2% formic acid in 50% acetonitrile), pooled, and dried under vacuum. The resulting peptides were labeled using TMTpro 16plex reagents (TFS; Lot #XL348283) according to the manufacturer’s instructions. After labeling for 15 min and quenching with 5% hydroxylamine, labeled peptides were combined in equal amounts across all channels, desalted using Sep-Pak tC18 cartridges (Waters), and dried again by vacuum centrifugation. To reduce ratio compression and systematic bias, the TMT channels were systematically assigned rather than randomized: HEK293T reference (channel 126), controls (127N–131C), and Protea-Tac groups (132N–134N). Pooled peptides were fractionated under high-pH conditions using an Agilent 1260 HPLC system equipped with a ZORBAX 300Extend-C18 column (4.6 mm × 150 mm, 3.5 μm; Agilent). Peptides were separated using a 55-min gradient, increasing buffer B (15 mM ammonium hydroxide in 90% acetonitrile [pH 10.0]) from 5% to 35% over 43 min at a flow rate of 0.2–1.0 mL/min. Ninety-six fractions were collected and concatenated into 24 fractions, which were vacuum-dried and stored at −80°C until MS analysis.

For LC-MS/MS analysis, each fraction was reconstituted in 10 μL of 0.1% formic acid/2% acetonitrile solution, and 2 μL were injected onto a trap column (75 µm I.D. × 2 cm, 3 µm PepMap100 C18 beads) before separation on an EASY-Spray analytical C18 column (75 µm × 50 cm, 2 μm particle size; TFS) using an Ultimate 3000 UHPLC system coupled to a Q-Exactive HF-X mass spectrometer (TFS). A 130-min linear gradient ramping solvent B (0.1% formic acid in acetonitrile) from 8% to 60% was applied at a flow rate of 300 nL/min. MS1 scans were acquired in positive ion mode over a mass range of m/z 350–1,800 at a resolution of 120,000 (FWHM at m/z 200), with an AGC target of 3×10⁶ and maximum injection time of 25 ms. The top 15 most intense precursor ions were selected for higher-energy collisional dissociation fragmentation using a 0.7 Th isolation window, normalized collision energy of 32%, and MS2 scans acquired at 30,000 resolution, with an AGC target of 1×10⁵ and maximum injection time of 80 ms. Raw MS data were analyzed using Proteome Discoverer ver. 3.1 (TFS), employing the Sequest HT search engine and Percolator algorithm to control the false discovery rate (FDR) at 1% at both the peptide and protein levels. The MS/MS spectra were searched against the UniProt Swiss-Prot human canonical protein sequence database (retrieved on March 2024; 20,360 entries), appended with common contaminants. Precursor mass tolerance was set at ±10 ppm, and fragment mass tolerance at ±0.02 Da. Peptide-spectrum matches were filtered to include only those with Sequest HT Xcorr ≥1, high Percolator confidence, and reporter ion signal-to-noise ratios >10, with spectra showing isolation interference >70% excluded. Quantification was performed using both unique and razor peptides, applying isotopic impurity correction factors provided by the manufacturer, and proteins identified with fewer than three unique peptides were excluded from further quantitative analysis. The TMT-MS proteomics data have been deposited in the ProteomeXchange Consortium via the PRIDE partner repository with the data set identifier PXD063133.

### Tumor xenograft/allograft mouse model

All experiments related to animals were performed according to the protocols accredited by the Institutional Animal Care and Use Committee (IACUC) of Seoul National University (IACUC no. SNU-240510-3-2), the Catholic University of Korea (IACUC no. CUK-IACUC-2026-015) and SML biopharm (IACUC no. SML-BP-IACUC-2025-11). Mice were housed in a pathogen-free facility fully approved by the Institutional Animal Care and Use Committee (IACUC). EGFP or scFv(c-Fos)-ADRM1 stably expressing cells were suspended in a 1:1 mixture of PBS and Matrigel (Corning, USA), and 100 µL of the suspension (5 × 10^6^ cells) was subcutaneously injected into the right flank region of a 6-week-old immunodeficient NOD.Cg-PrkdcscidIl2rgtm1Wjl/SzJ (NSG) mice (The Jackson Laboratory) using a 26-gauge syringe. In case of CT26 allograft, CT26 cells were resuspended in PBS and 150 µL (5 × 10^6^ cells) of the suspension was subcutaneously injected into the right flank of 5-week-old female BALB/cAnNCrl (Orient Bio, Seongnam, Korea) using a 26-gauge syringe. To establish the A549 xenograft model, A549 cells were resuspended in a 1:1 mixture of PBS and Matrigel. A total volume of 100 µL (3 × 10^6^ cells) was injected subcutaneously into the right flank of 6-week-old female BALB/cAnN-Foxn1nu/Crl mice (Orient Bio, Seongnam, Korea) using a 26-gauge syringe. Tumor size and body weight were monitored twice a week using a caliper. Tumor volume was calculated using the formula (length × width²) / 2. When the tumor of EGFP expressing cells reached approximately 1000 mm^3^, every mouse was sacrificed and the tumor tissues were removed for subsequent biochemical analyses.

### *In vivo* Protea-Tac delivery using AAV

When the MCF7 xenograft tumors reached a volume of approximately 100 mm³, AAV serotype 2 (AAV2) encoding surviving-driven EGFP or scFv(c-Fos)-ADRM1 was administered intratumorally three times per week (2 × 10⁹ vg in PBS). To ensure tumor-specific expression, all transgenes were placed under the control of the 269-bp survivin core promoter. When tumors in the PBS- or AAV2-survivin-EGFP–treated groups reached approximately 600 mm³, mice were euthanized, and tumor tissues were collected for subsequent biochemical analyses. In case of CT26 allograft assay, when the CT26 allograft tumors reached a volume of approximately 150 mm³, AAV2 encoding survivin-driven EGFP or scFv(c-Fos)-ADRM1 was administered intratumorally one time per week (4 × 10⁹ vg in PBS). When tumors in the PBS- or AAV2-survivin-EGFP–treated groups reached approximately 1200 mm³, mice were euthanized, and tumor tissues were collected for subsequent biochemical analyses.

### Serum biochemical and IL-6 analyses

At the experimental endpoint, blood samples were collected and serum was isolated for subsequent analyses. Serum levels of alanine aminotransferase (ALT), aspartate aminotransferase (AST), alkaline phosphatase (ALP), uric acid (UA), creatinine (CREA), and blood urea nitrogen (BUN) were analyzed by KP&T (South Korea). Serum IL-6 concentrations were determined using a Mouse IL-6 ELISA Kit (Invitrogen, KMC0061) according to the manufacturer’s instructions. Absorbance was measured at 450 nm using a microplate reader, and IL-6 concentrations were calculated from a standard curve.

### Preparation of mRNA

To generate mRNA, a DNA template includes the 5ʹ-untranslated region (UTR), 3ʹ-UTR, and poly-A tails. The DNA template featured multiple cloning sites containing four distinct restriction sequences, facilitating the precise insertion of target genes, such as EGFP, scFv(c-Fos), PSMD4-scFv(c-Fos) and scFv(c-Fos)-ADRM1. Before in vitro transcription (IVT), the plasmid DNA was completely linearized using the Not Ⅰ (Enzynomics, Daejeon, South Korea) restriction enzyme. IVT was then conducted at 37 °C for 4 h using the T7 RNA polymerase (Enzynomics, Daejeon, South Korea) and its associated buffer system to produce mRNA from DNA templates. Uridine was entirely replaced with N1-methyl-pseudouridine (Trilink, San Diego, CA, USA) to minimize innate immune activation, and a Cap 1 analog was used (SMARTCAP®, ST PHARM, Seoul, South Korea) during the transcription process according to the manufacturer’s instructions. Following the reaction, the template DNA was digested with DNase Ⅰ (Enzynomics, Daejeon, South Korea) at 37 °C for 30 min. The synthesized mRNA was precipitated via lithium chloride and cellulose-based purification to remove double-stranded RNA (dsRNA) contaminants. The final mRNA products were characterized for integrity using 1% denaturing agarose gel electrophoresis, and their concentration and purity were assessed using a NanoDrop-2000 spectrophotometer (Thermo Fisher Scientific, Waltham).

### Formulation of mRNA-LNP

To formulate mRNA-encapsulated lipid nanoparticles (LNP), we utilized lipids including ALC-0315, 1,2-distearoyl-sn-glycero-3-phosphocholine (DSPC), ALC-0159 and cholesterol (Avanti Polar Lipids, USA). For mRNA-LNP formulation, the lipid phase was dissolved in absolute ethanol and mixed with an acidic mRNA aqueous phase (50 mM citrate, pH 4.0) maintaining an N/P ratio of 6. NanoAssemblr® IgniteTM (Precision Nanosystems Inc., Canada) was used for formulating mRNA-LNP. The solutions were mixed at a 1:3 volume ratio (organic to aqueous solutions) and a total flow rate of 10 mL/min. To ensure complete solvent exchange and particle stabilization, the synthesized mRNA-LNPs were first subjected to a two-step dialysis process. The formulations were dialyzed against Tris Buffer for 2 h at room temperature, followed by an overnight incubation at 4°C. In accordance with the manufacturer’s recommendations, the final mRNA-LNPs were concentrated using an Amicon® Ultra-15 Centrifugal Filter (Merck Millipore, Germany). Dynamic light scattering (DLS) was performed on a Zetasizer Nano ZS (Malvern Panalytical, Malvern, Worcestershire, United Kingdom) to determine particle size with all samples diluted in Tris buffer. Furthermore, the encapsulation efficiency was quantified through a Ribo-green assay using Quant-it™ Ribo-Green RNA Assay Kit (Thermo Fisher, Waltham, MA, USA). Total RNA was released via Triton X-100 treatment, enabling the determination of encapsulation efficiency by comparing internalized versus free RNA.

### *in vivo* Protea-Tac delivery using mRNA-LNP

When the average tumor volume reached approximately 100 mm³, the mice were randomly assigned to treatment groups. The mice were treated with a subcutaneous injection of 40 μg of mRNA per dose formulated in LNPs encoding EGFP, scFv(c-Fos), PSMD4-scFv(c-Fos) in a total volume of 50 µL, while the control group was injected with 50 µL of PBS via the same route. Treatments were administered every 5 days for a total of three doses. Tumor growth was monitored every 2 to 3 days. 11 days after the final injection, the mice were euthanized via CO2 asphyxiation. Blood was collected from the caudal vena cava for serum biochemical analysis, and tumor tissues were harvested for further analyses.

### Densitometry and Statistical analysis

Band intensities from immunoblots were quantified by densitometric analysis using iBright Analysis Software (v5.4.0). Statistical significance between groups was assessed using an unpaired t-test, one-way ANOVA, or two-way ANOVA followed by Tukey’s and Sidak’s multiple comparison tests as post hoc analyses (GraphPad Prism, ver. 10.4.1). A p-value of less than 0.05 was considered statistically significant. For TMT-MS, protein abundance data were processed in Perseus (ver. 1.6.15.0). Reporter ion intensities were log₂-transformed, and width adjustment normalization was applied to equalize interquartile ranges across samples. Principal component analysis was conducted to visualize overall sample variation and to assess the separation between controls and experimental groups. Student’s t-test was performed to identify differentially expressed proteins, applying a Benjamini-Hochberg FDR threshold < 0.05 and a fold-change (FC) cutoff of >1.2 or <0.833, corresponding to a 20% increase or decrease in expression (|log₂FC| > 0.263).

## Acknowledgements

This work was supported by the grants from the National Research Foundation of Korea (RS-2025-16652968 and RS-2026-25477259 to M.J.L.) and the Samsung Research Funding & Incubation Center of Samsung Electronics (SRFC-MA2202-06 to C.H.L., J-.H.N. and M.J.L.).

## Author Contributions

S.H.P. and Y.J. designed and carried out most experiments and data analyses unless otherwise stated. S.L. and J.N. prepared mRNA, formulated mRNA-LMP and conducted xenograft experiments. E.K. and D.H. performed TMT-MS analysis. I.B., J.K., and D.J. conducted the xenograft experiments, VHH-based Protea-Tac-related works, and FACS analysis, respectively. J.Y. and C.H.L engineered the c-Fos-targeting scFv antibodies. S.Y.L. and J.H.N. performed the *in vivo* mRNA-LNP delivery experiments. S.H.P., Y.J. and M.J.L. prepared the manuscript. M.J.L. conceived the study and supervised the project.

## Data, Materials, and Software availability

The raw TMT-MS/MS proteomics data have been deposited at the ProteomeXchange Consortium via the PRIDE partner repository (PXD063133). All other data supporting the findings of this study are available from the corresponding author on request.

