## Supplementary figures and legends for "Ubiquitin Receptor-Mediated, Ubiquitin-Independent Targeted Protein Degradation via 26S Proteasomes"

**This PDF file includes:**

**Supplementary Figures 1 to 7**
**Supplementary Table 1**

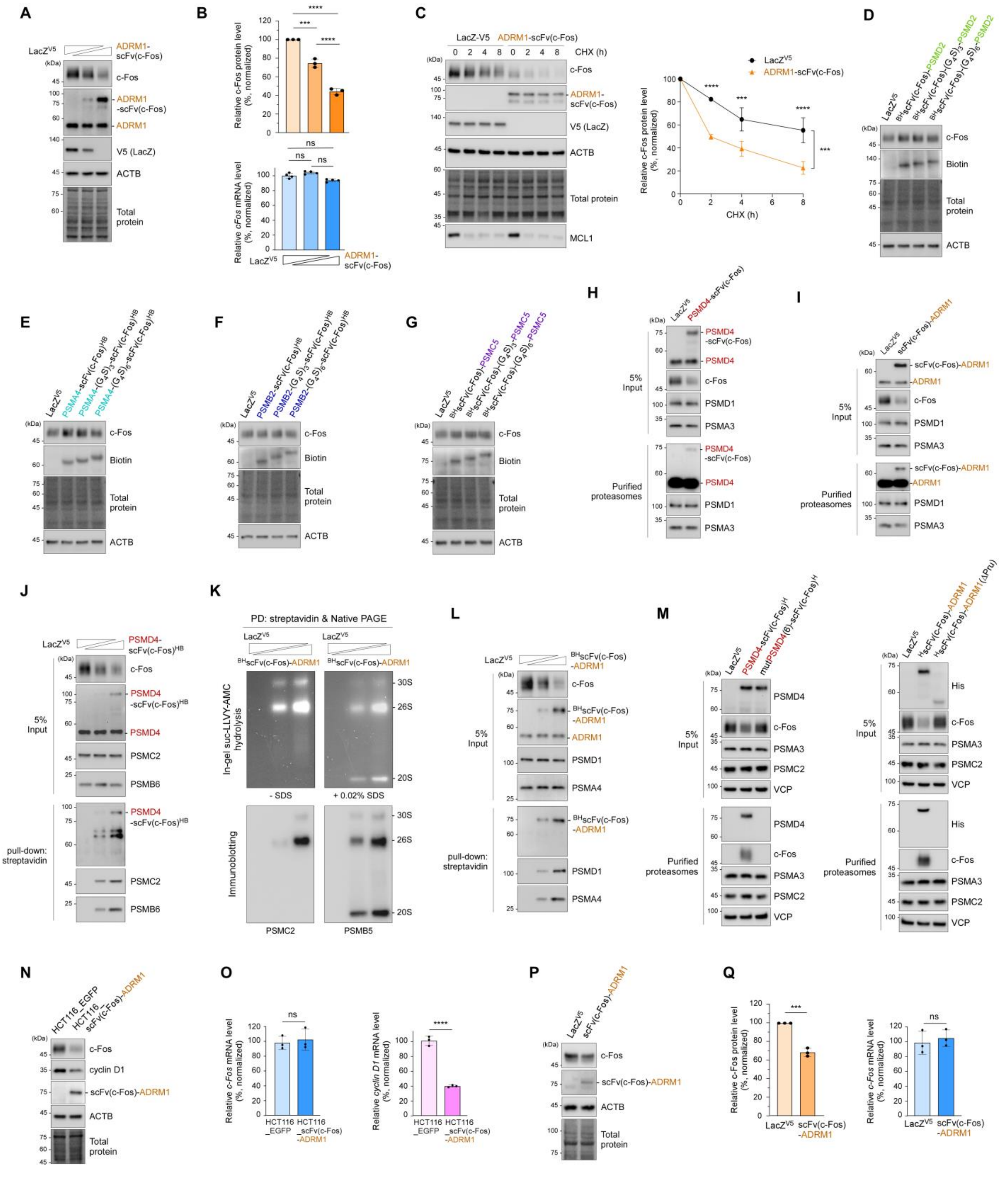

**Fig. S1. Proteasome-targeting chimera (Protea-Tac), composed of a Ub receptor (UbR) and**

**a targeting antibody (tAb), promotes efficient degradation of target proteins, including c-Fos oncoprotein. Related to Figure 1.**

(**A**) A549 cells were co-transfected with plasmids expressing LacZ$^{V5}$ controls (1, 0.7, and 0 μg) or ADRM1-scFv(c-Fos) degraders (0, 0.3, and 1 μg), along with plasmids expressing target protein c-Fos (0.5 μg). After 64 h, whole-cell lysates (WCLs) were prepared and analyzed by SDS-PAGE/immunoblotting (IB) with indicated antibodies, followed by total protein staining. Representative IB results from three independent experiments are shown. (**B**) (*top*) Quantification of c-Fos band intensities from (A), normalized to total protein signals. Data represent mean ± SD (N = 3). ***p < 0.001, ****p < 0.0001 (one-way ANOVA with Tukey's post hoc test). (*bottom*) Total RNA was extracted and analyzed by quantitative RT-PCR (qRT-PCR) using primers targeting *c-Fos* and *GAPDH* (as a normalization control) mRNA. Data represent the mean ± SD from four independent experiments (N = 4). ns, not significant (one-way ANOVA with Tukey's multiple comparison test). (**C**) Same as (A), except that chase experiments were performed by adding 80 μg/mL cycloheximide (CHX) and harvesting samples at 2, 4, or 8 h. c-Fos levels were normalized to total protein and to the values at 0 h post-CHX treatment (set as 100%) of each group (LacZ$^{V5}$ controls / ADRM1-scFv(c-Fos) degraders). Average values of normalized c-Fos levels (mean ± SD) from three independent experiments are shown (N = 3, ***p < 0.001, ****p < 0.0001 from two-way ANOVA followed by Sidak's multiple comparison test). (**D**–**G**) PSMD2 (D), PSMA4 (E), PSMB2 (F), and PSMC5 (G) were fused to scFv(c-Fos) using linkers of varying lengths and evaluated for TPD activity. (**H**) HEK293 cells stably expressing biotin-tagged PSMB2 were transfected with either LacZ$^{V5}$ or PSMD4-scFv(c-Fos) together with c-Fos-expressing plasmids for 64 h. Affinity purification of 26S proteasomes was conducted, and the samples were analyzed by IB. (**I**) The ADRM1-based degrader (scFv(c-Fos)-ADRM1) was evaluated in the same methods as in (H). (**J**) PSMD4-scFv(c-Fos) degraders with C-terminal hexahistidine-biotin (HB) tags were transiently expressed for streptavidin affinity purification. Pulldown samples and WCLs were examined by denaturing SDS-PAGE/IB using antibodies against c-Fos and proteasome subunits. (**K**) Similar to (J), except that N-terminally biotin-hexahistidine (BH)-tagged scFv(c-Fos)-ADRM1 was evaluated. Samples were analyzed by non-denaturing (native) PAGE, followed by in-gel suc-LLVY-AMC hydrolysis visualization (*top*) and IB with antibodies against the CP (PSMB5) and RP (PSMC2) subcomplexes (*bottom*). (**L**) Similar to (J), except N-terminally biotin-hexahistidine (BH)-tagged scFv(c-Fos)-ADRM1 was analyzed. (**M**) Protea-Tac degraders harboring mutant PSMD4 (where six key residues in its VWA domain were substituted with alanine; D11A, S13A, T15A, I84A, G115A, and S116A) and Pru-domain-truncated ADRM1 were compared with their cognate Protea-Tac degraders in terms of c-Fos degradation and proteasomal incorporation. (**N**) Endogenous levels of c-Fos and cyclin D1 were compared in HCT116 cells stably overexpressing EGFP or scFv(c-Fos)-ADRM1. (**O**) As in (N), except mRNA levels of endogenous *c-Fos* (top) and *cyclin D1* (bottom) were measured by qRT-PCR and normalized to *GAPDH*. ****p < 0.0001 (N = 3, two-tailed Student's *t*-test). (**P**) HCT116 cells were transiently transfected with either LacZ$^{V5}$ or scFv(c-Fos)-ADRM1 for 48 h, followed by IB analysis to determine c-Fos levels. Representative IB results from three independent experiments are shown. (**Q**) (left) Quantification of c-Fos protein levels in (P), normalized to total protein signals. Data represent the mean ± SD from three independent experiments. ***p < 0.001 (N = 3, two-tailed Student's t-test). (right) As in (P), but c-Fos mRNA levels were quantified by qRT-PCR and normalized to GAPDH. Data are presented as the mean ± SD. ns, not significant (N = 3, two-tailed Student's t-test).

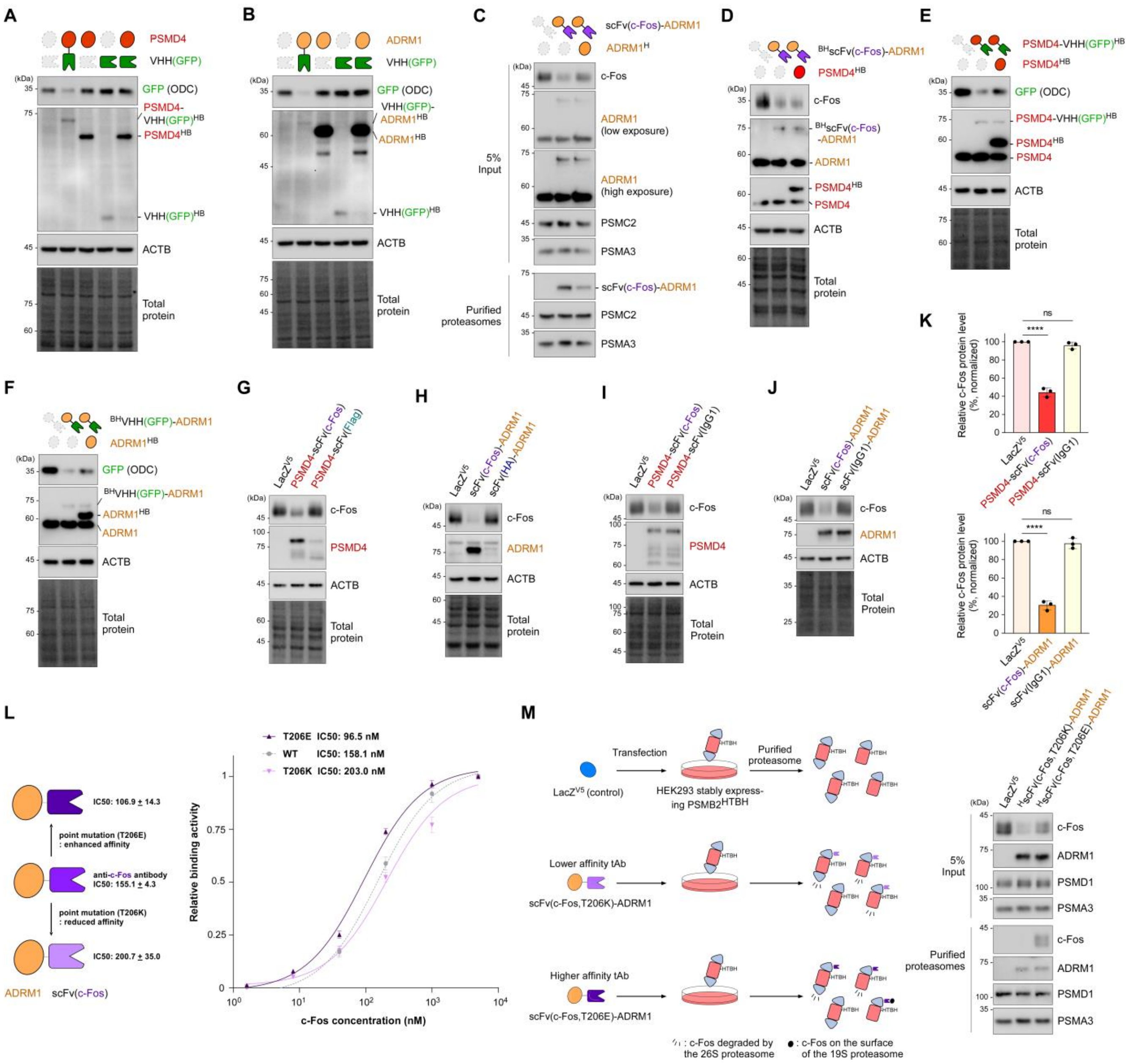


**fig. S2. Protea-Tac is a modular system requiring fused UbR and tAb components. Related to Figure 2.**

(**A**) A549 cells were co-transfected with plasmids expressing PSMD4, VHH(GFP), either individually, or as a PSMD4-VHH (GFP) chimera, along with plasmids expressing GFP-ODC. LacZ$^{V5}$-expressing plasmids were used as a control to maintain a total of 1.5 μg plasmid DNA per condition. After 64 h, WCLs were prepared and subjected to SDS-PAGE/IB with the indicated antibodies, followed by total protein staining. (**B**) As in (A), except that ADRM1 was used as the UbR rather than PSMD4. (**C**) HEK293 cells stably expressing biotin-tagged PSMB2 were transfected with scFv(c-Fos)-ADRM1 and c-Fos-expressing plasmids in the presence or absence of hexahistidine-tagged ADRM1 ($ADRM1^H$) to assess competitive incorporation into proteasomes and target degradation efficiency. Proteasomes were affinity-purified, and the samples were analyzed by denaturing SDS-PAGE/IB using antibodies against c-Fos and proteasome subunits. (**D**) As in (C), except that PSMD4 (non-cognate UbR) was co-expressed with scFv(c-Fos)-ADRM1 in A549 cells. (**E** and **F**) Similar to (D), but PSMD4 or ADRM1 was co-expressed with their corresponding Protea-Tac constructs (PSMD4-VHH(GFP) or VHH(GFP)-ADRM1) to evaluate their competitive effects on GFP-ODC degradation. (**G** and **H**) Protea-Tac degraders with scFv(Flag) and scFv(HA) were compared with PSMD4-scFv(c-Fos) and scFv(c-Fos)-ADRM1 chimeras, respectively, in terms of c-Fos degradation. (**I** and **J**) Protea-Tac chimeras of PSMD4 and ADRM1 fused to a mutant scFv construct whose CDR regions were substituted with that of human IgG1 were examined. (**K**) (*top*) Quantification of c-Fos band intensities from (I), normalized to total protein. (*bottom*) c-Fos signal quantification from (J). Three independent experiments are shown. Bars represent the mean ± SD (N = 3). ****$p < 0.0001$ (one-way ANOVA with Tukey's post hoc test). (**L**) (*left*) Diagram of scFv(c-Fos) variants with reduced and increased binding affinities toward recombinant c-Fos, obtained through specific point mutations (T206K and T206E, respectively; *see* Methods for a detailed screening process). (*right*) Dose-response curves displaying the binding affinities of recombinant c-Fos-targeting scFvs for GST-tagged c-Fos. Immobilized scFvs were incubated with four-fold serial dilutions of GST-c-Fos (5000 – 0.32 nM) in an ELISA format. $IC_{50}$ values were calculated from four-parameter-logistics equations (GraphPad Prism, ver. 10.4.1). The mean values were determined from three independent experiments, with error bars representing standard deviation (N = 3 each group). (**M**) (*left*) Schematic illustration of the experimental design assessing ternary complex stability among the Protea-Tac degrader, target substrate (c-Fos), and the 26S proteasome as a function of the tAb module's binding affinity. (*right*) HEK293T cells stably expressing biotin- and hexahistidine-tagged PSMB2 (PSMB2-HTBH) were transiently transfected with either control $LacZ^{V5}$, low-affinity scFv(c-Fos, T206K)-ADRM1, or high-affinity scFv(c-Fos, T206E)-ADRM1 constructs. Following streptavidin pull-down under native conditions, c-Fos and various proteasome subunits were analyzed by SDS-PAGE/IB with the indicated antibodies.

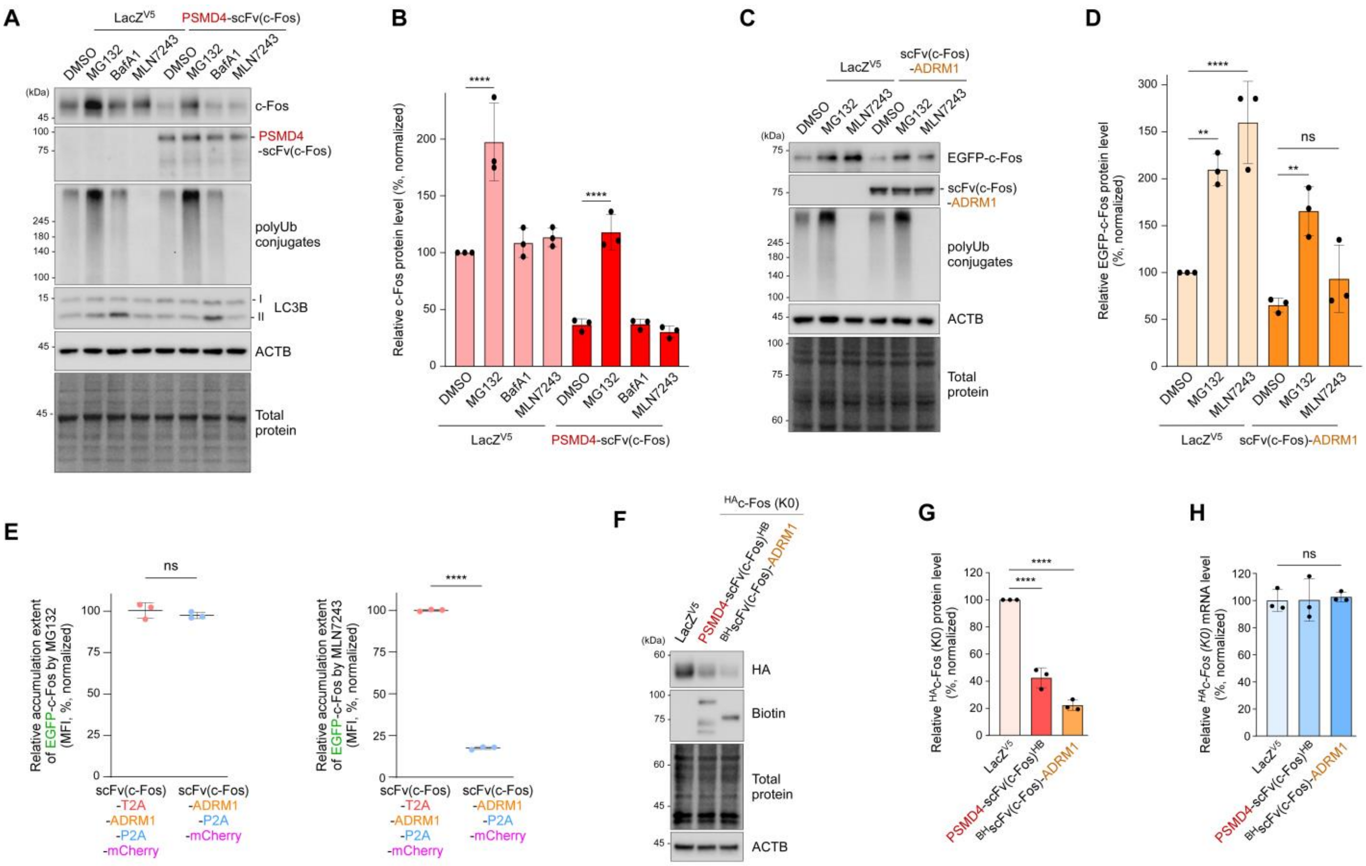


**fig. S3. The Protea-Tac degraders induce Ub-independent and proteasome-mediated TPD. Related to Figure 3.**

(**A**) The PSMD4-scFv(c-Fos)-mediated c-Fos degradation assay was performed with DMSO, the proteasome inhibitor MG132 (10 μM), the autophagic flux inhibitor bafilomycin A1 (BafA1; 200 nM), or the E1 Ub-activating enzyme inhibitor MLN7243 (1 μM) for 6 h to determine pathway dependence. (**B**) Quantification of the steady-state levels of c-Fos from (A), normalized to total protein. Bars represent the mean ± SD (N = 3). ****$p < 0.0001$ (one-way ANOVA with Tukey's post hoc test). (**C**) Similar to (A), but here the scFv(c-Fos)-ADRM1 was tested in the presence of MG132 (10 μM for 6 h) or MLN7243 (1 μM for 6 h) to evaluate its TPD activity toward EGFP-c-Fos. (**D**) Quantification of (C), normalized to total protein. Bars represent the mean ± SD (N = 3). **$p < 0.01$, ****$p < 0.0001$; ns, not significant (one-way ANOVA with Tukey's post hoc test). (**E**) HEK293T cells stably expressing EGFP-c-Fos were transfected with the tricistronic control construct (red) or the bicistronic Protea-Tac construct (blue). After DMSO or MG132 (10 μM for 6 h) or MLN7243 (1 μM for 6 h) treatment, mean MFI value from EGFP-c-Fos in mCherry-positive cells were determined by flow cytometry. Extent of MFI value accumulation compared to DMSO group are shown (*left*: MG132, *right*: MLN7243). Bars indicate the mean ± SD (N = 3). ****$p < 0.0001$; ns, not significant (two-tailed Student's t test). (**F**) A549 cells were co-transfected with lysine-less $^{HA}$c-Fos (K0) and either LacZ$^{V5}$ or the indicated anti-c-Fos Protea-Tac constructs for 64 h. WCLs were analyzed by SDS-PAGE followed by IB and total protein staining.

Representative IB results from three independent experiments are shown. (**G**) Quantification of $^{HA}$c-Fos (K0) levels normalized to total protein signals. Bars represent the mean ± SD (N = 3). ****$p < 0.0001$ (one-way ANOVA with Tukey's post hoc test). (**H**) As in (F) except total RNA was extracted for quantitative RT-PCR using $^{HA}$c-Fos (K0) and GAPDH (normalization control) primers. Values represent the mean ± SD of three independent experiments (N = 3). ns, not significant (one-way ANOVA with Tukey's post hoc test).

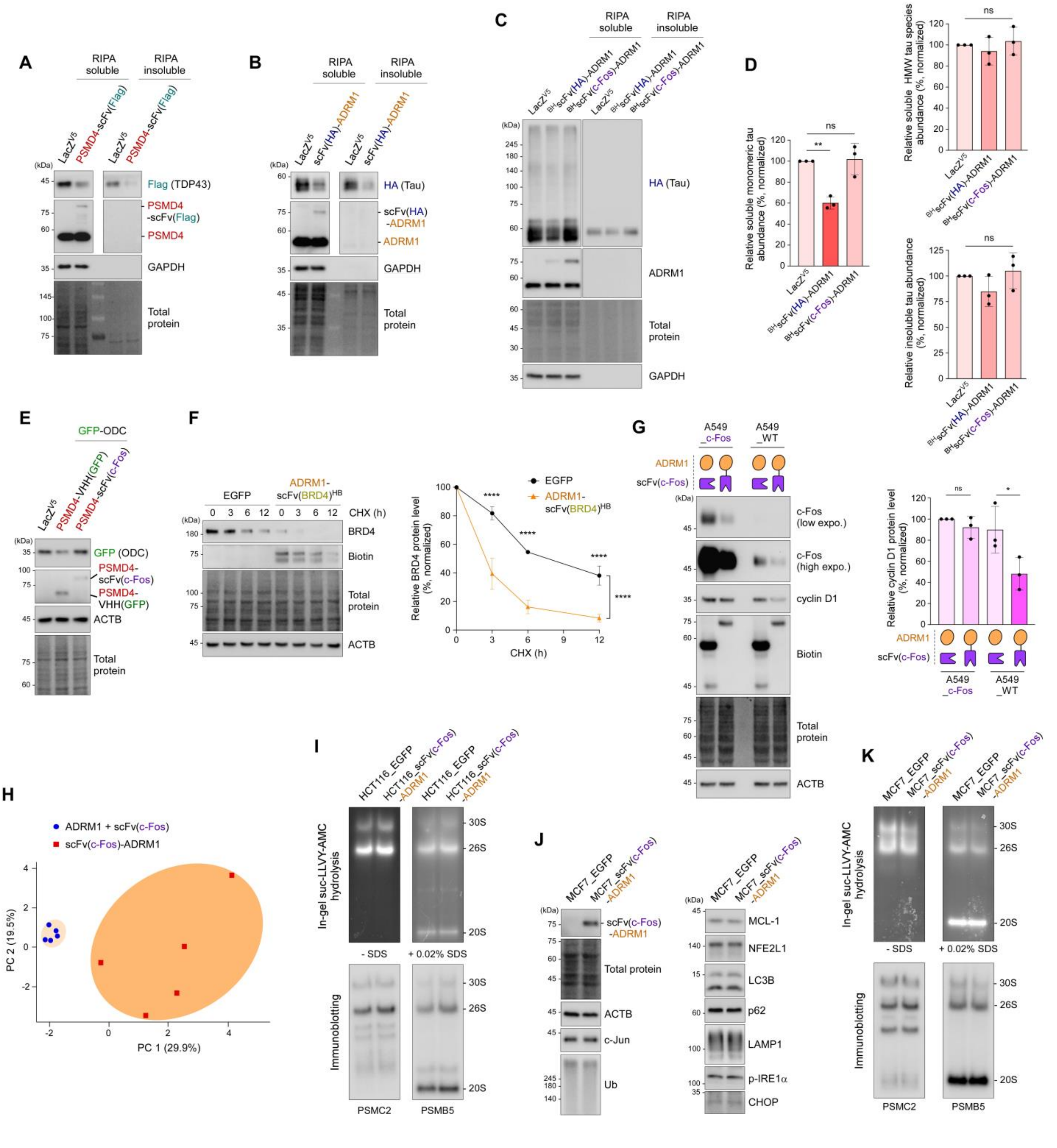


**fig. S4. The Protea-Tac chimeras allow for specific TPD of diverse proteins without affecting the global proteome. Related to Figure 4.**

(**A**) A549 cells were co-transfected with $^{Flag}$TDP43 (target protein; 0.5 μg) and either PSMD4-

scFv(Flag) (Flag degrader; 1 μg) or $LacZ^{V5}$ (transfection control; 1 μg) for 72 h. WCLs were separated into RIPA-soluble and -insoluble (pellet) fractions and examined using SDS-PAGE/IB and total protein staining. (**B**) Same as (A), except $^{HA}$Tau and scFv(HA)-ADRM1 were used as a target and degrader, respectively. (**C**) HEK293 cells stably expressing HA-tau(0N4R) were transfected with either $LacZ^{V5}$ or $^{BH}$scFv(HA)-ADRM1 or $^{BH}$scFv(c-Fos)-ADRM1 for 64 h. Cells were lysed in RIPA buffer, whole cell lysates were separated into supernatant and pellet fractions and subjected to non-reducing (supernatants) and reducing (pellets) SDS-PAGE/IB. Representative IB results from three independent experiments are shown. (**D**) Left: quantification of RIPA-soluble, monomeric HA-tau(0N4R) levels normalized to total protein signals. *Top right*: levels of soluble, high-molecular-weight HA-tau(0N4R) species were quantified, normalized to total protein signals. *Bottom right*: RIPA-insoluble HA-tau(0N4R) levels were quantified, normalized to total protein signals. Bars represent the mean ± SD (N = 3). $^{**}p < 0.01$, ns, not significant (one-way ANOVA with Tukey's post hoc test). (**E**) Specificity of Protea-Tac was assessed using mismatched target-degrader pairs. (**F**) A549 cells were treated with EGFP or anti-BRD4 Protea-Tac (ADRM1-scFv$(BRD4)^{HB}$) encoding lenti virus for 72 h and chase experiments were performed at 3, 6, and 12 h following treatment with 80 μg/mL cycloheximide (CHX) at time zero. BRD4 signals were normalized to the values at 0 h chase (set as 100%) of each group and to total protein levels. Average percentages of remaining BRD4 (mean ± SD) from three independent experiments (N = 3, $^{****}p < 0.0001$ from two-way ANOVA followed by Sidak's post hoc test). (**G**) A549 cells stably expressing c-Fos and A549 WT cells were transfected with either $ADRM1^{HB}$ plus scFv$(c\text{-}Fos)^{HB}$ or $^{BH}$scFv(c-Fos)-ADRM1, for 64 h. WCLs were analyzed by SDS-PAGE followed by IB. *Left*: Representative IB results from three independent experiments are shown. *Right*: Quantification of cyclin D1 levels normalized to total protein signals. Bars represent the mean ± SD (N = 3). ns, not significant, $^{*}p < 0.05$ (one-way ANOVA with Tukey's post hoc test). (**H**) Principal component analysis (PCA) plots depicting replicates from the control (N = 5; blue circles) and Protea-Tac-treated groups (N = 5; red rectangles). (**I**) Analysis of 26S proteasomes in HCT116 cells stably expressing EGFP or scFv(c-Fos)-ADRM1 using native PAGE, followed by in-gel suc-LLVY-AMC hydrolysis visualization (*top*) and subsequent IB with antibodies against the CP (PSMB5) and the RP (PSMC2) (*bottom*). 20S proteasomes were activated by 0.02% SDS. (**J**) Levels of various endogenous proteins were compared in MCF7_EGFP and MCF7_scFv(c-Fos)-ADRM1 stable cell lines by SDS-PAGE/IB with the appropriate antibodies. (**K**) As in (I), except that MCF7_EGFP and MCF7_scFv(c-Fos)-ADRM1 stable cell lines were analyzed.

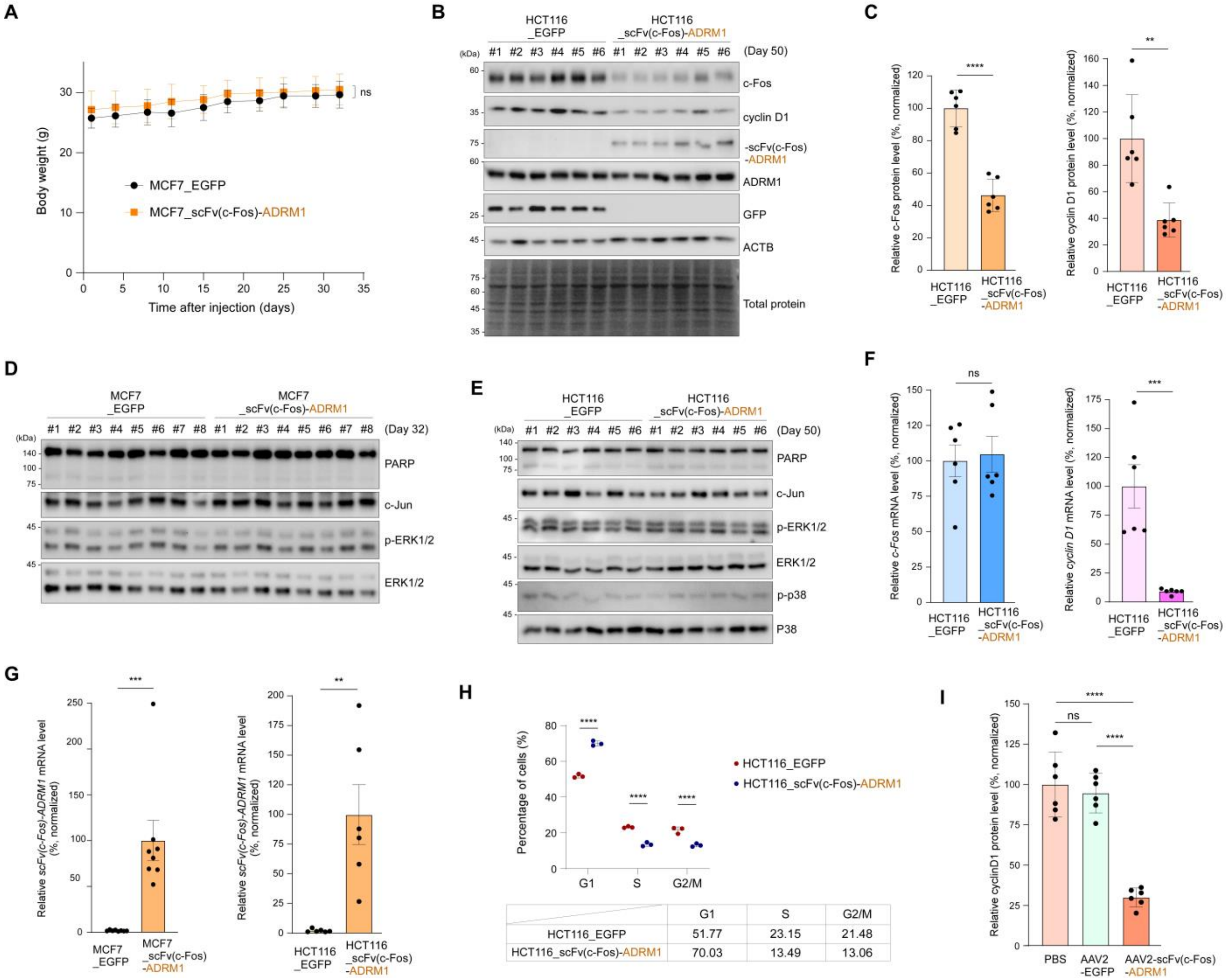


| | G1 | S | G2/M |
|---|---|---|---|
| HCT116_EGFP | 51.77 | 23.15 | 21.48 |
| HCT116_scFv(c-Fos)-ADRM1 | 70.03 | 13.49 | 13.06 |

**fig. S5. Protea-Tac-induced c-Fos degradation leads to anti-tumor efficacy *in vivo*. Related to Figure 5.**

(**A**) Body weight changes of NOD/SCID/IL-2γ-receptor null (NSG) mice (male, six-week-old) xenografted with MCF7 cells stably overexpressing either EGFP or scFv(c-Fos)-ADRM1 (N = 8 each) throughout the 32-day experiment period. (**B**) Whole lysates of HCT116 xenograft tumors were subjected to SDS-PAGE/IB using the indicated antibodies. (**C**) Quantification of c-Fos and cyclin D1 protein levels in (B), normalized to total protein signals. Bars represent the mean ± SD (N = 6). $^{**}p < 0.01$ and $^{****}p < 0.0001$ (two-tailed Student's t-test). (**D** and **E**) IB analysis of xenograft tumor lysates from MCF7 (D) and HCT116 (E) stable cells, probed for apoptosis markers and upstream regulators of c-Fos. (**F**) qRT-PCR analysis of *c-Fos* and *cyclin D1* transcript levels in HCT116 xenograft tumors (N = 6). Quantitative normalization was performed using *GAPDH* mRNA expression. (**G**) As in (F), except that mRNA levels of *scFv(c-Fos)-ADRM1* were determined in biological triplicates. (**H**) Cell cycle distribution in the HCT116 stable cell lines used for xenograft assays, analyzed by flow cytometry. DNA contents were quantified using DAPI staining. ****$p$<0.0001 between EGFP- and Protea-Tac-expressing tumors, two-way ANOVA followed by Bonferroni's multiple-comparisons test (N = 3 each). (**I**) Quantification of cyclin D1 protein levels from the IB analysis in Fig. 5J. Data are shown as mean ± SD (N = 6). ****$p$ < 0.0001 (one-way ANOVA with Tukey's post hoc test). ns, not significant.

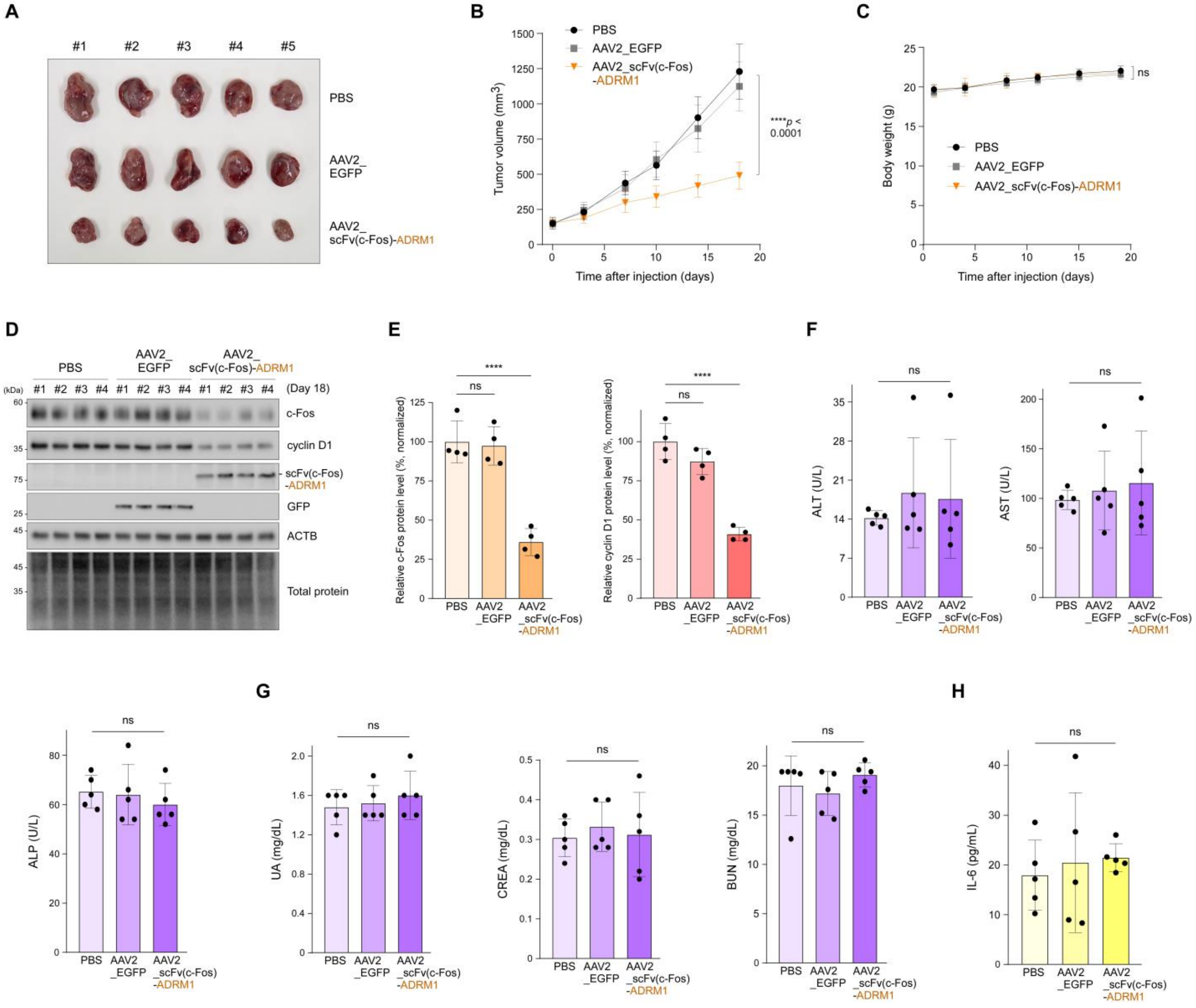


**fig. S6. AAV2-mediated Protea-Tac expression suppresses tumor growth in an immunocompetent syngeneic tumor model. Related to Figure 5.**

(**A**) CT26 cells were subcutaneously inoculated into five-week-old female BALB/cAnNCrl mice. When tumors reached approximately 150 mm³, mice were intratumorally injected with PBS or AAV2 encoding EGFP or scFv(c-Fos)-ADRM1 ($4 \times 10^9$ vg each) once weekly for a total of three injections. Representative images of tumors harvested on day 18 after the first injection are shown. (**B**) Tumor volumes were measured at the indicated time points for 18 days. Data are presented as means ± SEM (N = 5 per group). ****$p < 0.0001$ (two-way ANOVA followed by Tukey's post hoc test). (**C**) Body weight changes during the experiment. Data are presented as means ± SEM (N = 5 per group). ns, not significant (two-way ANOVA followed by Tukey's post hoc test). (**D**) IB analysis of whole-tumor lysates with the indicated antibodies. (**E**) Quantification of c-Fos (left) and cyclin D1 (right) protein levels in (D), normalized to total protein signals. Bars represent the mean ± SD (N = 4). ns, not significant; ****$p < 0.0001$ (one-way ANOVA followed by Tukey's post hoc test). (**F and G**) Serum samples collected at the experimental endpoint were analyzed for alanine aminotransferase (ALT; F), aspartate aminotransferase (AST; F), alkaline phosphatase (ALP; F), uric acid (UA; G), creatinine (CREA; G), and blood urea nitrogen (BUN; G) levels. Bars represent the mean ± SD (N = 5). ns, not significant (one-way ANOVA followed by Tukey's post hoc test). (H) Serum IL-6 concentrations were measured at the experimental endpoint by ELISA. Bars represent the mean ± SD (N = 5). ns, not significant (one-way ANOVA followed by Tukey's post hoc test).

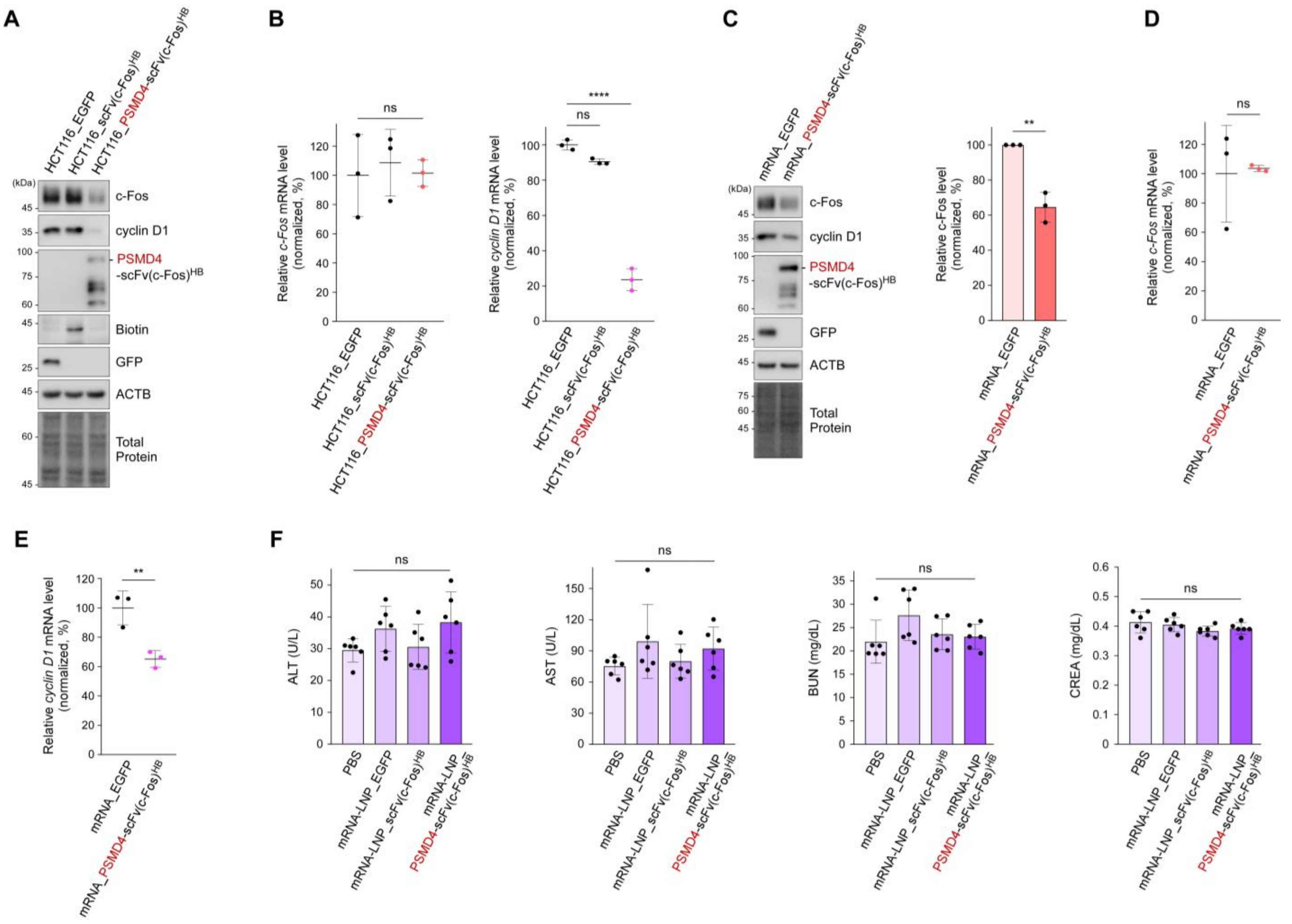


**fig. S7. Endogenous substrate degradation and mRNA mediated delivery of PSMD4-based Protea-Tac. Related to Figure 6.**

(**A**) Endogenous c-Fos and cyclin D1 protein levels were compared in HCT116 cells stably expressing EGFP, scFv(c-Fos)$^{HB}$ or PSMD4-scFv(c-Fos)$^{HB}$. Single-cell clones with the highest expression level were selected for analysis. (**B**) As in (A) except that mRNA levels of endogenous *c-Fos* (*left*) and *cyclin D1* (*right*) were evaluated by qRT-PCR and normalized to GAPDH. ns, not significant; ****p < 0.0001 (one-way ANOVA with Tukey's post hoc test). (**C**) A549 cells were transfected with EGFP or PSMD4-scFv(c-Fos)$^{HB}$ encoding mRNA for 36 h. WCLs were analyzed by SDS-PAGE and IB. *Left*: Representative IB results from three independent experiments are shown. *Right*: Quantification of c-Fos levels normalized to total protein signals. Bars represent the mean ± SD (N = 3). **p < 0.01 (two-tailed Student's *t*-test). (**D**) As in (C) except that mRNA levels of endogenous *c-Fos* were evaluated by qRT-PCR and normalized to GAPDH. Values are presented as means ± SD. ns, not significant (N = 3, two-tailed Student's t-test). (**E**) As in (C) except that mRNA levels of endogenous *cyclin D1* were evaluated by qRT-PCR and normalized to GAPDH. Values are presented as means ± SD. **p < 0.01 (N = 3, two-tailed Student's t-test). (**F**) Serum samples collected from the mice used in the xenograft experiment shown in Figure 6 at the experimental endpoint were analyzed for alanine aminotransferase (ALT), aspartate aminotransferase (AST), blood urea nitrogen (BUN), and creatinine (CREA) levels. Bars represent the mean ± SD (N = 6). ns, not significant (one-way ANOVA followed by Tukey's post hoc test).

### Supplementary Table 1. Coding sequences of chimeras.

**Color code : Proteasome subunit – Linker – scFv/VHH**

| | |
|---|---|
| PSMD4-<br>scFv(c-Fos) | ATGGTGTTGGAAAGCACTATGGTGTGTGTGGACAACAGTGAGTATATGCGGAATGGAGACTTCTTACCCACCAGGC<br>TGCAGGCCCAGCAGGATGCTGTCAACATAGTTTGTCATTCAAAGACCCGCAGCAACCCTGAGAACAACGTGGGCC<br>TTATCACACTGGCTAATGACTGTGAAGTGCTGACCACACTCACCCCAGACACTGGCCGTATCCTGTCCAAGCTACA<br>TACTGTCCAACCCAAGGGCAAGATCACCTTCTGCACGGGCATCCGCGTGGCCCATCTGGCTCTGAAGCACCGACA<br>AGGCAAGAATCACAAGATGCGCATCATTGCCTTTGTGGGAAGCCCAGTGGAGGACAATGAGAAGGATCTGGTGAA<br>ACTGGCTAAACGCCTCAAGAAGGAGAAAGTAAATGTTGACATTATCAATTTTGGGGAAGAGGAGGTGAACACAGA<br>AAAGCTGACAGCCTTTGTAAACACGTTGAATGGCAAAGATGGAACCGGTTCTCATCTGGTGACAGTGCCTCCTGG<br>GCCCAGTTTGGCTGATGCTCTCATCAGTTCTCCGATTTTGGCTGGTGAAGGTGGTGCCATGCTGGGTCTTGGTGCC<br>AGTGACTTTGAATTTGGAGTAGATCCCAGTGCTGATCCTGAGCTGGCCTTGGCCCTTCGTGTATCTATGGAAGAGC<br>AGCGGCAGCGGCAGGAGGAGGAGGCCCGGCGGGCAGCTGCAGCTTCTGCTGCTGAGGCCGGGATTGCTACGACT<br>GGGACTGAAGACTCAGACGATGCCCTGCTGAAGATGACCATCAGCCAGCAAGAGTTTGGCCGCACTGGGCTTCCT<br>GACCTAAGCAGTATGACTGAGGAAGAGCAGATTGCTTATGCCATGCAGATGTCCCTGCAGGGAGCAGAGTTTGGC<br>CAGGCGGAATCAGCAGACATTGATGCCAGCTCAGCTATGGACACATCTGAGCCAGCCAAGGAGGAGGATGATTAC<br>GACGTGATGCAGGACCCCGAGTTCCTTCAGAGTGTCCTAGAGAACCTCCCAGGTGTGGATCCCAACAATGAAGCC<br>ATTCGAAATGCTATGGGCTCCCTGGCCTCCCAGGCCACCAAGGACGGCAAGAAGGACAAGAAGGAGGAAGACAA<br>GAAGCGACCCAAGCTTGGTACCGAGCTCGGATCCATGGAAGTGCAGCTGCAGCAGAGCGGAGCCGAGCTGGTGC<br>GGAGCGGGGCCTCTGTGAAACTGTCTTGCACCGCTTCTGGCTTCAACATCAAGGACTACTACATGCACTGGGTCA<br>AGCAGAGACCTGAGCAGGGCCTGGAATGGATCGGCTGGATGGACCCCAAGAACGGCGATACCGAGTACGCCCCTA<br>AGTTCCAGGGCAAGGCCACCATGACCGCCGACACCAGCAGCAATACCGCCTACCTGCAGCTGAGCAGCCTTACAA<br>GCGACGACACAGCCGTGTACTACTGCCACGACGCCATCACCACGCCCACCAGAAACTTCGACGTGTGGGGCGCCG<br>GCACAACCGTGACCGTGTCTTCTGGCGGCGGAGGCAGCGGCGGAGGCGGCAGCGGCGGCGGCGGCAGCGGAGG<br>CGGCTCTGATGTGGTGATGACCCAGACACCTCTGAGCCTGCCTGTGTCCCTGGGCGACCAGGCCAGCATCAGCTG<br>TAGAAGCTCTCAGAGCCTGGTGCACAGCAACGGCAACACCTACCTGCACTGGTACCTGCAAAAGCCCGGCCAGTC<br>TCCAAAGCTGCTGATGTACAAGGTGTCCAATAGATTCAGCGGTGTTCCTGATAGATTTAGCGGCAGCGGGAGCGGA<br>ACCGACTTCACCCTGAAGATCAGCAGAGTGGAAGCCGAAGATCTGGGAGTGTACTTCTGCAGCCAGTCAACACAC<br>GTGCCTTTCACCTTCGGCTCCGGCACCAAGCTCGAGATCAAG |
| scFv(c-Fos)-<br>ADRM1 | ATGGAAGTGCAGCTGCAGCAGAGCGGAGCCGAGCTGGTGCGGAGCGGGGCCTCTGTGAAACTGTCTTGCACCGC<br>TTCTGGCTTCAACATCAAGGACTACTACATGCACTGGGTCAAGCAGAGACCTGAGCAGGGCCTGGAATGGATCGG<br>CTGGATGGACCCCAAGAACGGCGATACCGAGTACGCCCCTAAGTTCCAGGGCAAGGCCACCATGACCGCCGACAC<br>CAGCAGCAATACCGCCTACCTGCAGCTGAGCAGCCTTACAAGCGACGACACAGCCGTGTACTACTGCCACGACGC<br>CATCACCACGCCCACCAGAAACTTCGACGTGTGGGGCGCCGGCACAACCGTGACCGTGTCTTCTGGCGGCGGAG<br>GCAGCGGCGGAGGCGGCAGCGGCGGCGGCGGCAGCGGAGGCGGCTCTGATGTGGTGATGACCCAGACACCTCTG<br>AGCCTGCCTGTGTCCCTGGGCGACCAGGCCAGCATCAGCTGTAGAAGCTCTCAGAGCCTGGTGCACAGCAACGGC<br>AACACCTACCTGCACTGGTACCTGCAAAAGCCCGGCCAGTCTCCAAAGCTGCTGATGTACAAGGTGTCCAATAGAT<br>TCAGCGGTGTTCCTGATAGATTTAGCGGCAGCGGGAGCGGAACCGACTTCACCCTGAAGATCAGCAGAGTGGAAG<br>CCGAAGATCTGGGAGTGTACTTCTGCAGCCAGTCAACACACGTGCCTTTCACCTTCGGCTCCGGCACCAAGCTCG<br>AGATCAAGGAATTCATGACGACCTCAGGCGCGCTCTTTCCAAGCCTGGTGCCAGGCTCTCGGGGCGCCTCCAACA<br>AGTACTTGGTGGAGTTTCGGGCGGGAAAGATGTCCCTGAAGGGGACCACCGTGACTCCGGATAAGCGGAAAGGG<br>CTGGTGTACATTCAGCAGACGGACGACTCGCTTATTCACTTCTGCTGGAAGGACAGGACGTCCGGGAACGTGGAA<br>GACGACTTGATCATCTTCCCTGACGACTGTGAGTTCAAGCGGGTGCCGCAGTGCCCCAGCGGGAGGGTCTACGTG<br>CTGAAGTTCAAGGCAGGGTCCAAGCGGCTTTTCTTCTGGATGCAGGAACCCAAGACAGACCAGGATGAGGAGCAT<br>TGCCGGAAAGTCAACGAGTATCTGAACAACCCCCCGATGCCTGGGGCGCTGGGGGCCAGCGGAAGCAGCGGCCA<br>CGAACTCTCTGCGCTAGGCGGTGAGGGTGGCCTGCAGAGCCTGCTGGGAAACATGAGCCACAGCCAGCTCATGCA<br>GCTCATCGGACCAGCCGGCCTTGGAGGACTGGGTGGGCTGGGGGCCCTGACTGGACCTGGCCTGGCCAGCTTACT<br>GGGGAGCAGTGGGCCTCCAGGGAGCAGCTCCTCCTCCAGCTCCCGGAGCCAGTCGGCAGCGGTCACCCCGTCAT<br>CCACCACCTCTTCCACCCGTGCCACCCCAGCCCCTTCTGCTCCAGCAGCTGCCTCAGCAACTAGCCCGAGCCCCGC<br>GCCCAGTTCCGGGAATGGAGCCAGCACAGCAGCCAGCCCGACCCAGCCCATCCAGCTGAGCGACCTCCAGAGCA<br>TCCTGGCCACGATGAACGTACCAGCCGGGCCAGCAGGCGGCCAGCAAGTGGACCTGGCCAGTGTGCTGACGCCG<br>GAGATAATGGCTCCCATCCTCGCCAACGCGGATGTCCAGGAGCGCCTGCTTCCCTACTTGCCATCTGGGGAGTCGC<br>TGCCGCAGACCGCGGATGAGATCCAGAATACCCTGACCTCGCCCCAGTTCCAGCAGGCCCTGGGCATGTTCAGCG<br>CAGCCTTGGCCTCGGGGCAGCTGGGCCCCCTCATGTGCCAGTTCGGTCTGCCTGCAGAGGCTGTGGAGGCCGCCA<br>ACAAGGGCGATGTGGAAGCGTTTGCCAAAGCCATGCAGAACAACGCCAAGCCCGAGCAGAAAGAGGGCGACAC<br>GAAGGACAAGAAGGACGAAGAGGAGGACATGAGCCTGGAC |
| ADRM1-<br>scFv(c-Fos) | ATGACGACCTCAGGCGCGCTCTTTCCAAGCCTGGTGCCAGGCTCTCGGGGCGCCTCCAACAAGTACTTGGTGGAG<br>TTTCGGGCGGGAAAGATGTCCCTGAAGGGGACCACCGTGACTCCGGATAAGCGGAAAGGGCTGGTGTACATTCAG<br>CAGACGGACGACTCGCTTATTCACTTCTGCTGGAAGGACAGGACGTCCGGGAACGTGGAAGACGACTTGATCATC<br>TTCCCTGACGACTGTGAGTTCAAGCGGGTGCCGCAGTGCCCCAGCGGGAGGGTCTACGTGCTGAAGTTCAAGGCA<br>GGGTCCAAGCGGCTTTTCTTCTGGATGCAGGAACCCAAGACAGACCAGGATGAGGAGCATTGCCGGAAAGTCAA<br>CGAGTATCTGAACAACCCCCCGATGCCTGGGGCGCTGGGGGCCAGCGGAAGCAGCGGCCACGAACTCTCTGCGCT<br>AGGCGGTGAGGGTGGCCTGCAGAGCCTGCTGGGAAACATGAGCCACAGCCAGCTCATGCAGCTCATCGGACCAG<br>CCGGCCTTGGAGGACTGGGTGGGCTGGGGGCCCTGACTGGACCTGGCCTGGCCAGCTTACTGGGGAGCAGTGGG |

| | |
|---|---|
| | CCTCCAGGGAGCAGCTCCTCCTCCAGCTCCCGGAGCCAGTCGGCAGCGGTCACCCCGTCATCCACCACCTCTTCC<br>ACCCGTGCCACCCCAGCCCCTTCTGCTCCAGCAGCTGCCTCAGCAACTAGCCCGAGCCCCGCGCCCAGTTCCGGG<br>AATGGAGCCAGCACAGCAGCCAGCCCGACCCAGCCCATCCAGCTGAGCGACCTCCAGAGCATCCTGGCCACGATG<br>AACGTACCAGCCGGGCCAGCAGGCGGCCAGCAAGTGGACCTGGCCAGTGTGCTGACGCCGGAGATAATGGCTCC<br>CATCCTCGCCAACGCGGATGTCCAGGAGCGCCTGCTTCCCTACTTGCCATCTGGGGAGTCGCTGCCGCAGACCGCG<br>GATGAGATCCAGAATACCCTGACCTCGCCCCAGTTCCAGCAGGCCCTGGGCATGTTCAGCGCAGCCTTGGCCTCGG<br>GGCAGCTGGGCCCCCTCATGTGCCAGTTCGGTCTGCCTGCAGAGGCTGTGGAGGCCGCCAACAAGGGCGATGTGG<br>AAGCGTTTGCCAAAGCCATGCAGAACAACGCCAAGCCCGAGCAGAAAGAGGGCGACACGAAGGACAAGAAGGA<br>CGAAGAGGAGGACATGAGCCTGGACAAGCTTGGTACCGAGCTCGGATCCATGGAAGTGCAGCTGCAGCAGAGCG<br>GAGCCGAGCTGGTGCGGAGCGGGGCCTCTGTGAAACTGTCTTGCACCGCTTCTGGCTTCAACATCAAGGACTACT<br>ACATGCACTGGGTCAAGCAGAGACCTGAGCAGGGCCTGGAATGGATCGGCTGGATGGACCCCAAGAACGGCGAT<br>ACCGAGTACGCCCCTAAGTTCCAGGGCAAGGCCACCATGACCGCCGACACCAGCAGCAATACCGCCTACCTGCAG<br>CTGAGCAGCCTTACAAGCGACGACACAGCCGTGTACTACTGCCACGACGCCATCACCACGCCCACCAGAAACTTC<br>GACGTGTGGGGCGCCGGCACAACCGTGACCGTGTCTTCTGGCGGCGGAGGCAGCGGCGGAGGCGGCAGCGGCG<br>GCGGCGGCAGCGGAGGCGGCTCTGATGTGGTGATGACCCAGACACCTCTGAGCCTGCCTGTGTCCCTGGGCGACC<br>AGGCCAGCATCAGCTGTAGAAGCTCTCAGAGCCTGGTGCACAGCAACGGCAACACCTACCTGCACTGGTACCTGC<br>AAAAGCCCGGCCAGTCTCCAAAGCTGCTGATGTACAAGGTGTCCAATAGATTCAGCGGTGTTCCTGATAGATTTAG<br>CGGCAGCGGGAGCGGAACCGACTTCACCCTGAAGATCAGCAGAGTGGAAGCCGAAGATCTGGGAGTGTACTTCT<br>GCAGCCAGTCAACACACGTGCCTTTCACCTTCGGCTCCGGCACCAAGCTCGAGATCAAG |
| scFv(c-Fos)-<br>PSMD2 | ATGGAAGTGCAGCTGCAGCAGAGCGGAGCCGAGCTGGTGCGGAGCGGGGCCTCTGTGAAACTGTCTTGCACCGC<br>TTCTGGCTTCAACATCAAGGACTACTACATGCACTGGGTCAAGCAGAGACCTGAGCAGGGCCTGGAATGGATCGG<br>CTGGATGGACCCCAAGAACGGCGATACCGAGTACGCCCCTAAGTTCCAGGGCAAGGCCACCATGACCGCCGACAC<br>CAGCAGCAATACCGCCTACCTGCAGCTGAGCAGCCTTACAAGCGACGACACAGCCGTGTACTACTGCCACGACGC<br>CATCACCACGCCCACCAGAAACTTCGACGTGTGGGGCGCCGGCACAACCGTGACCGTGTCTTCTGGCGGCGGAG<br>GCAGCGGCGGAGGCGGCAGCGGCGGCGGCGGCAGCGGAGGCGGCTCTGATGTGGTGATGACCCAGACACCTCTG<br>AGCCTGCCTGTGTCCCTGGGCGACCAGGCCAGCATCAGCTGTAGAAGCTCTCAGAGCCTGGTGCACAGCAACGGC<br>AACACCTACCTGCACTGGTACCTGCAAAAGCCCGGCCAGTCTCCAAAGCTGCTGATGTACAAGGTGTCCAATAGAT<br>TCAGCGGTGTTCCTGATAGATTTAGCGGCAGCGGGAGCGGAACCGACTTCACCCTGAAGATCAGCAGAGTGGAAG<br>CCGAAGATCTGGGAGTGTACTTCTGCAGCCAGTCAACACACGTGCCTTTCACCTTCGGCTCCGGCACCAAGCTCG<br>AGATCAAGGAATTCATGGAGGAGGGAGGCCGGGACAAGGCGCCGGTGCAGCCCCAGCAGTCTCCAGCGGCGGCC<br>CCCGGCGGCACGGACGAGAAGCCGAGCGGCAAGGAGCGGCGGGATGCCGGGGACAAGGACAAAGAACAGGAG<br>CTGTCTGAAGAGGATAAACAGCTTCAAGATGAACTGGAGATGCTCGTGGAACGACTAGGGGAGAAGGATACATCC<br>CTGTATCGACCAGCGCTGGAGGAATTGCGAAGGCAGATTCGTTCTTCTACAACTTCCATGACTTCAGTGCCCAAGC<br>CTCTCAAATTTCTGCGTCCACACTATGGCAAACTGAAGGAAATCTATGAGAACATGGCCCCTGGGGAGAATAAGCG<br>TTTTGCTGCTGACATCATCTCCGTTTTGGCCATGACCATGAGTGGGGAGCGTGAGTGCCTCAAGTATCGGCTAGTG<br>GGCTCCCAGGAGGAATTGGCATCATGGGGTCATGAGTATGTCAGGCATCTGGCAGGAGAAGTGGCTAAGGAGTGG<br>CAGGAGCTGGATGACGCAGAGAAGGTCCAGCGGGAGCCTCTGCTCACTCTGGTGAAGGAAATCGTCCCCTATAAC<br>ATGGCCCACAATGCAGAGCATGAGGCTTGCGACCTGCTTATGGAAATTGAGCAGGTGGACATGCTGGAGAAGGAC<br>ATTGATGAAAATGCATATGCAAAGGTCTGCCTTTATCTCACCAGTTGTGTGAATTACGTGCCTGAGCCTGAGAACTC<br>AGCCCTACTGCGTTGTGCCCTGGGTGTGTTCCGAAAGTTTAGCCGCTTCCCTGAAGCTCTGAGATTGGCATTGATG<br>CTCAATGACATGGAGTTGGTAGAAGACATCTTCACCTCCTGCAAGGATGTGGTAGTACAGAAACAGATGGCATTCA<br>TGCTAGGCCGGCATGGGGTGTTCCTGGAGCTGAGTGAAGATGTCGAGGAGTATGAGGACCTGACAGAGATCATGT<br>CCAATGTACAGCTCAACAGCAACTTCTTGGCCTTAGCTCGGGAGCTGGACATCATGGAGCCCAAGGTGCCTGATG<br>ACATCTACAAAACCCACCTAGAGAACAACAGGTTTGGGGGCAGTGGCTCTCAGGTGGACTCTGCCCGCATGAACC<br>TGGCCTCCTCTTTTGTGAATGGCTTTGTGAATGCAGCTTTTGGCCAAGACAAGCTGCTAACAGATGATGGCAACAA<br>ATGGCTTTACAAGAACAAGGACCACGGAATGTTGAGTGCAGCTGCATCTCTTGGGATGATTCTGCTGTGGGATGTG<br>GATGGTGGCCTCACCCAGATTGACAAGTACCTGTACTCCTCTGAGGACTACATTAAGTCAGGAGCTCTTCTTGCCT<br>GTGGCATAGTGAACTCTGGGGTCCGGAATGAGTGTGACCCTGCTCTGGCACTGCTCTCAGACTATGTTCTCCACAA<br>CAGCAACACCATGAGACTTGGTTCCATCTTTGGGCTAGGCTTGGCTTATGCTGGCTCAAATCGTGAAGATGTCCTA<br>ACACTGCTGCTGCCTGTGATGGGAGATTCAAAGTCCAGCATGGAGGTGGCAGGTGTCACAGCTTTAGCCTGTGGA<br>ATGATAGCAGTAGGGTCCTGCAATGGAGATGTAACTTCCACTATCCTTCAGACCATCATGGAGAAGTCAGAGACTG<br>AGCTCAAGGATACTTATGCTCGTTGGCTTCCTCTTGGACTGGGTCTCAACCACCTGGGGAAGGGTGAGGCCATCGA<br>GGCAATCCTGGCTGCACTGGAGGTTGTGTCAGAGCCATTCCGCAGTTTTGCCAACACACTGGTGGATGTGTGTGCA<br>TATGCAGGCTCTGGGAATGTGCTGAAGGTGCAGCAGCTGCTCCACATTTGTAGCGAACACTTTGACTCCAAAGAG<br>AAGGAGGAAGACAAAGACAAGAAGGAAAAGAAAGACAAGGACAAGAAGGAAGCCCCTGCTGACATGGGAGCA<br>CATCAGGGAGTGGCTGTTCTGGGGATTGCCCTTATTGCTATGGGGGAGGAGATTGGTGCAGAGATGGCATTACGAA<br>CCTTTGGCCACTTGCTGAGATATGGGGAGCCTACACTCCGGAGGGCTGTACCTTTAGCACTGGCCCTCATCTCTGTT<br>TCAAATCCACGACTCAACATCCTGGATACCCTAAGCAAATTCTCTCATGATGCTGATCCAGAAGTTTCCTATAACTC<br>CATTTTTGCCATGGGCATGGTGGGCAGTGGTACCAATAATGCCCGTCTGGCTGCAATGCTGCGCCAGTTAGCTCAAT<br>ATCATGCCAAGGACCCAAACAACCTCTTCATGGTGCGCTTGGCACAGGGCCTGACACATTTAGGGAAGGGCACCC<br>TTACCCTCTGCCCCTACCACAGCGACCGGCAGCTTATGAGCCAGGTGGCCGTGGCTGGACTGCTCACTGTGCTTGT<br>CTCTTTCCTGGATGTTCGAAACATTATTCTAGGCAAATCACACTATGTATTGTATGGGCTGGTGGCTGCCATGCAGCC<br>CCGAATGCTGGTTACGTTTGATGAGGAGCTGCGGCCATTGCCAGTGTCTGTCCGTGTGGGCCAGGCAGTGGATGTG<br>GTGGGCCAGGCTGGCAAGCCGAAGACTATCACAGGGTTCCAGACGCATACAACCCCAGTGTTGTTGGCCCACGGG<br>GAACGGGCAGAATTGGCCACTGAGGAGTTTCTTCCTGTTACCCCCATTCTGGAAGGTTTTGTTATCCTTCGGAAGA<br>ACCCCAATTATGATCTC |
| scFv(c-Fos)- | ATGGAAGTGCAGCTGCAGCAGAGCGGAGCCGAGCTGGTGCGGAGCGGGGCCTCTGTGAAACTGTCTTGCACCGC<br>TTCTGGCTTCAACATCAAGGACTACTACATGCACTGGGTCAAGCAGAGACCTGAGCAGGGCCTGGAATGGATCGG |

| | |
|---|---|
| (GGGGS)$_3$-<br>PSMD2 | CTGGATGGACCCCAAGAACGGCGATACCGAGTACGCCCCTAAGTTCCAGGGCAAGGCCACCATGACCGCCGACAC<br>CAGCAGCAATACCGCCTACCTGCAGCTGAGCAGCCTTACAAGCGACGACACAGCCGTGTACTACTGCCACGACGC<br>CATCACCACGCCCACCAGAAACTTCGACGTGTGGGGCGCCGGCACAACCGTGACCGTGTCTTCTGGCGGCGGAG<br>GCAGCGGCGGAGGCGGCAGCGGCGGCGGCGGCAGCGGAGGCGGCTCTGATGTGGTGATGACCCAGACACCTCTG<br>AGCCTGCCTGTGTCCCTGGGCGACCAGGCCAGCATCAGCTGTAGAAGCTCTCAGAGCCTGGTGCACAGCAACGGC<br>AACACCTACCTGCACTGGTACCTGCAAAAGCCCGGCCAGTCTCCAAAGCTGCTGATGTACAAGGTGTCCAATAGAT<br>TCAGCGGTGTTCCTGATAGATTTAGCGGCAGCGGGAGCGGAACCGACTTCACCCTGAAGATCAGCAGAGTGGAAG<br>CCGAAGATCTGGGAGTGTACTTCTGCAGCCAGTCAACACACGTGCCTTTCACCTTCGGCTCCGGCACCAAGCTCG<br>AGATCAAGGAATTCGGAGGAGGAGGGTCCGGTGGGGGAGGCTCTGGAGGGGGTGGGTCTGAATTCATGGAGGAG<br>GGAGGCCGGGACAAGGCGCCGGTGCAGCCCCAGCAGTCTCCAGCGGCGGCCCCCGGCGGCACGGACGAGAAGC<br>CGAGCGGCAAGGAGCGGCGGGATGCCGGGGACAAGGACAAAGAACAGGAGCTGTCTGAAGAGGATAAACAGCT<br>TCAAGATGAACTGGAGATGCTCGTGGAACGACTAGGGGAGAAGGATACATCCCTGTATCGACCAGCGCTGGAGGA<br>ATTGCGAAGGCAGATTCGTTCTTCTACAACTTCCATGACTTCAGTGCCCAAGCCTCTCAAATTTCTGCGTCCACACT<br>ATGGCAAACTGAAGGAAATCTATGAGAACATGGCCCCTGGGGAGAATAAGCGTTTTGCTGCTGACATCATCTCCGT<br>TTTGGCCATGACCATGAGTGGGGAGCGTGAGTGCCTCAAGTATCGGCTAGTGGGCTCCCAGGAGGAATTGGCATC<br>ATGGGGTCATGAGTATGTCAGGCATCTGGCAGGAGAAGTGGCTAAGGAGTGGCAGGAGCTGGATGACGCAGAGA<br>AGGTCCAGCGGGAGCCTCTGCTCACTCTGGTGAAGGAAATCGTCCCCTATAACATGGCCCACAATGCAGAGCATG<br>AGGCTTGCGACCTGCTTATGGAAATTGAGCAGGTGGACATGCTGGAGAAGGACATTGATGAAAATGCATATGCAA<br>AGGTCTGCCTTTATCTCACCAGTTGTGTGAATTACGTGCCTGAGCCTGAGAACTCAGCCCTACTGCGTTGTGCCCT<br>GGGTGTGTTCCGAAAGTTTAGCCGCTTCCCTGAAGCTCTGAGATTGGCATTGATGCTCAATGACATGGAGTTGGTA<br>GAAGACATCTTCACCTCCTGCAAGGATGTGGTAGTACAGAAACAGATGGCATTCATGCTAGGCCGGCATGGGGTGT<br>TCCTGGAGCTGAGTGAAGATGTCGAGGAGTATGAGGACCTGACAGAGATCATGTCCAATGTACAGCTCAACAGCA<br>ACTTCTTGGCCTTAGCTCGGGAGCTGGACATCATGGAGCCCAAGGTGCCTGATGACATCTACAAAACCCACCTAGA<br>GAACAACAGGTTTGGGGGCAGTGGCTCTCAGGTGGACTCTGCCCGCATGAACCTGGCCTCCTCTTTTGTGAATGG<br>CTTTGTGAATGCAGCTTTTGGCCAAGACAAGCTGCTAACAGATGATGGCAACAAATGGCTTTACAAGAACAAGGA<br>CCACGGAATGTTGAGTGCAGCTGCATCTCTTGGGATGATTCTGCTGTGGGATGTGGATGGTGGCCTCACCCAGATT<br>GACAAGTACCTGTACTCCTCTGAGGACTACATTAAGTCAGGAGCTCTTCTTGCCTGTGGCATAGTGAACTCTGGGG<br>TCCGGAATGAGTGTGACCCTGCTCTGGCACTGCTCTCAGACTATGTTCTCCACAACAGCAACACCATGAGACTTGG<br>TTCCATCTTTGGGCTAGGCTTGGCTTATGCTGGCTCAAATCGTGAAGATGTCCTAACACTGCTGCTGCCTGTGATGG<br>GAGATTCAAAGTCCAGCATGGAGGTGGCAGGTGTCACAGCTTTAGCCTGTGGAATGATAGCAGTAGGGTCCTGCA<br>ATGGAGATGTAACTTCCACTATCCTTCAGACCATCATGGAGAAGTCAGAGACTGAGCTCAAGGATACTTATGCTCG<br>TTGGCTTCCTCTTGGACTGGGTCTCAACCACCTGGGGAAGGGTGAGGCCATCGAGGCAATCCTGGCTGCACTGGA<br>GGTTGTGTCAGAGCCATTCCGCAGTTTTGCCAACACACTGGTGGATGTGTGTGCATATGCAGGCTCTGGGAATGTG<br>CTGAAGGTGCAGCAGCTGCTCCACATTTGTAGCGAACACTTTGACTCCAAAGAGAAGGAGGAAGACAAAGACAA<br>GAAGGAAAAGAAAGACAAGGACAAGAAGGAAGCCCCTGCTGACATGGGAGCACATCAGGGAGTGGCTGTTCTG<br>GGGATTGCCCTTATTGCTATGGGGGAGGAGATTGGTGCAGAGATGGCATTACGAACCTTTGGCCACTTGCTGAGAT<br>ATGGGGAGCCTACACTCCGGAGGGCTGTACCTTTAGCACTGGCCCTCATCTCTGTTTCAAATCCACGACTCAACAT<br>CCTGGATACCCTAAGCAAATTCTCTCATGATGCTGATCCAGAAGTTTCCTATAACTCCATTTTTGCCATGGGCATGGT<br>GGGCAGTGGTACCAATAATGCCCGTCTGGCTGCAATGCTGCGCCAGTTAGCTCAATATCATGCCAAGGACCCAAAC<br>AACCTCTTCATGGTGCGCTTGGCACAGGGCCTGACACATTTAGGGAAGGGCACCCTTACCCTCTGCCCCTACCACA<br>GCGACCGGCAGCTTATGAGCCAGGTGGCCGTGGCTGGACTGCTCACTGTGCTTGTCTCTTTCCTGGATGTTCGAAA<br>CATTATTCTAGGCAAATCACACTATGTATTGTATGGGCTGGTGGCTGCCATGCAGCCCCGAATGCTGGTTACGTTTG<br>ATGAGGAGCTGCGGCCATTGCCAGTGTCTGTCCGTGTGGGCCAGGCAGTGGATGTGGTGGGCCAGGCTGGCAAGC<br>CGAAGACTATCACAGGGTTCCAGACGCATACAACCCCAGTGTTGTTGGCCCACGGGGAACGGGCAGAATTGGCCA<br>CTGAGGAGTTTCTTCCTGTTACCCCCATTCTGGAAGGTTTTGTTATCCTTCGGAAGAACCCCAATTATGATCTC |
| scFv(c-Fos)-<br>(GGGGS)$_6$-<br>PSMD2 | ATGGAAGTGCAGCTGCAGCAGAGCGGAGCCGAGCTGGTGCGGAGCGGGGCCTCTGTGAAACTGTCTTGCACCGC<br>TTCTGGCTTCAACATCAAGGACTACTACATGCACTGGGTCAAGCAGAGACCTGAGCAGGGCCTGGAATGGATCGG<br>CTGGATGGACCCCAAGAACGGCGATACCGAGTACGCCCCTAAGTTCCAGGGCAAGGCCACCATGACCGCCGACAC<br>CAGCAGCAATACCGCCTACCTGCAGCTGAGCAGCCTTACAAGCGACGACACAGCCGTGTACTACTGCCACGACGC<br>CATCACCACGCCCACCAGAAACTTCGACGTGTGGGGCGCCGGCACAACCGTGACCGTGTCTTCTGGCGGCGGAG<br>GCAGCGGCGGAGGCGGCAGCGGCGGCGGCGGCAGCGGAGGCGGCTCTGATGTGGTGATGACCCAGACACCTCTG<br>AGCCTGCCTGTGTCCCTGGGCGACCAGGCCAGCATCAGCTGTAGAAGCTCTCAGAGCCTGGTGCACAGCAACGGC<br>AACACCTACCTGCACTGGTACCTGCAAAAGCCCGGCCAGTCTCCAAAGCTGCTGATGTACAAGGTGTCCAATAGAT<br>TCAGCGGTGTTCCTGATAGATTTAGCGGCAGCGGGAGCGGAACCGACTTCACCCTGAAGATCAGCAGAGTGGAAG<br>CCGAAGATCTGGGAGTGTACTTCTGCAGCCAGTCAACACACGTGCCTTTCACCTTCGGCTCCGGCACCAAGCTCG<br>AGATCAAGGAATTCGGCGGAGGTGGCAGCGGTGGTGGCGGATCTGGCGGAGGTGGCTCTGGAGGAGGCGGTTCT<br>GGCGGAGGCGGCTCAGGTGGAGGTGGCAGCGAATTCATGGAGGAGGGAGGCCGGGACAAGGCGCCGGTGCAGC<br>CCCAGCAGTCTCCAGCGGCGGCCCCCGGCGGCACGGACGAGAAGCCGAGCGGCAAGGAGCGGCGGGATGCCGG<br>GGACAAGGACAAAGAACAGGAGCTGTCTGAAGAGGATAAACAGCTTCAAGATGAACTGGAGATGCTCGTGGAAC<br>GACTAGGGGAGAAGGATACATCCCTGTATCGACCAGCGCTGGAGGAATTGCGAAGGCAGATTCGTTCTTCTACAAC<br>TTCCATGACTTCAGTGCCCAAGCCTCTCAAATTTCTGCGTCCACACTATGGCAAACTGAAGGAAATCTATGAGAAC<br>ATGGCCCCTGGGGAGAATAAGCGTTTTGCTGCTGACATCATCTCCGTTTTGGCCATGACCATGAGTGGGGAGCGTG<br>AGTGCCTCAAGTATCGGCTAGTGGGCTCCCAGGAGGAATTGGCATCATGGGGTCATGAGTATGTCAGGCATCTGGC<br>AGGAGAAGTGGCTAAGGAGTGGCAGGAGCTGGATGACGCAGAGAAGGTCCAGCGGGAGCCTCTGCTCACTCTGG<br>TGAAGGAAATCGTCCCCTATAACATGGCCCACAATGCAGAGCATGAGGCTTGCGACCTGCTTATGGAAATTGAGCA<br>GGTGGACATGCTGGAGAAGGACATTGATGAAAATGCATATGCAAAGGTCTGCCTTTATCTCACCAGTTGTGTGAAT<br>TACGTGCCTGAGCCTGAGAACTCAGCCCTACTGCGTTGTGCCCTGGGTGTGTTCCGAAAGTTTAGCCGCTTCCCTG<br>AAGCTCTGAGATTGGCATTGATGCTCAATGACATGGAGTTGGTAGAAGACATCTTCACCTCCTGCAAGGATGTGGT |

| | |
|---|---|
| | AGTACAGAAACAGATGGCATTCATGCTAGGCCGGCATGGGGTGTTCCTGGAGCTGAGTGAAGATGTCGAGGAGTA<br>TGAGGACCTGACAGAGATCATGTCCAATGTACAGCTCAACAGCAACTTCTTGGCCTTAGCTCGGGAGCTGGACATC<br>ATGGAGCCCAAGGTGCCTGATGACATCTACAAAACCCACCTAGAGAACAACAGGTTTGGGGGCAGTGGCTCTCAG<br>GTGGACTCTGCCCGCATGAACCTGGCCTCCTCTTTTGTGAATGGCTTTGTGAATGCAGCTTTTGGCCAAGACAAGC<br>TGCTAACAGATGATGGCAACAAATGGCTTTACAAGAACAAGGACCACGGAATGTTGAGTGCAGCTGCATCTCTTG<br>GGATGATTCTGCTGTGGGATGTGGATGGTGGCCTCACCCAGATTGACAAGTACCTGTACTCCTCTGAGGACTACATT<br>AAGTCAGGAGCTCTTCTTGCCTGTGGCATAGTGAACTCTGGGGTCCGGAATGAGTGTGACCCTGCTCTGGCACTGC<br>TCTCAGACTATGTTCTCCACAACAGCAACACCATGAGACTTGGTTCCATCTTTGGGCTAGGCTTGGCTTATGCTGGC<br>TCAAATCGTGAAGATGTCCTAACACTGCTGCTGCCTGTGATGGGAGATTCAAAGTCCAGCATGGAGGTGGCAGGT<br>GTCACAGCTTTAGCCTGTGGAATGATAGCAGTAGGGTCCTGCAATGGAGATGTAACTTCCACTATCCTTCAGACCAT<br>CATGGAGAAGTCAGAGACTGAGCTCAAGGATACTTATGCTCGTTGGCTTCCTCTTGGACTGGGTCTCAACCACCTG<br>GGGAAGGGTGAGGCCATCGAGGCAATCCTGGCTGCACTGGAGGTTGTGTCAGAGCCATTCCGCAGTTTTGCCAAC<br>ACACTGGTGGATGTGTGTGCATATGCAGGCTCTGGGAATGTGCTGAAGGTGCAGCAGCTGCTCCACATTTGTAGCG<br>AACACTTTGACTCCAAAGAGAAGGAGGAAGACAAAGACAAGAAGGAAAAGAAAGACAAGGACAAGAAGGAAG<br>CCCCTGCTGACATGGGAGCACATCAGGGAGTGGCTGTTCTGGGGATTGCCCTTATTGCTATGGGGGAGGAGATTGG<br>TGCAGAGATGGCATTACGAACCTTTGGCCACTTGCTGAGATATGGGGAGCCTACACTCCGGAGGGCTGTACCTTTA<br>GCACTGGCCCTCATCTCTGTTTCAAATCCACGACTCAACATCCTGGATACCCTAAGCAAATTCTCTCATGATGCTGA<br>TCCAGAAGTTTCCTATAACTCCATTTTTGCCATGGGCATGGTGGGCAGTGGTACCAATAATGCCCGTCTGGCTGCAA<br>TGCTGCGCCAGTTAGCTCAATATCATGCCAAGGACCCAAACAACCTCTTCATGGTGCGCTTGGCACAGGGCCTGAC<br>ACATTTAGGGAAGGGCACCCTTACCCTCTGCCCCTACCACAGCGACCGGCAGCTTATGAGCCAGGTGGCCGTGGC<br>TGGACTGCTCACTGTGCTTGTCTCTTTCCTGGATGTTCGAAACATTATTCTAGGCAAATCACACTATGTATTGTATGG<br>GCTGGTGGCTGCCATGCAGCCCCGAATGCTGGTTACGTTTGATGAGGAGCTGCGGCCATTGCCAGTGTCTGTCCGT<br>GTGGGCCAGGCAGTGGATGTGGTGGGCCAGGCTGGCAAGCCGAAGACTATCACAGGGTTCCAGACGCATACAAC<br>CCCAGTGTTGTTGGCCCACGGGGAACGGGCAGAATTGGCCACTGAGGAGTTTCTTCCTGTTACCCCCATTCTGGAA<br>GGTTTTGTTATCCTTCGGAAGAACCCCAATTATGATCTC |
| PSMA4-<br>scFv(c-Fos) | ATGTCTCGAAGATATGACTCCAGGACCACTATATTTTCTCCAGAAGGTCGCTTATACCAAGTTGAATATGCCATGGA<br>AGCTATTGGACATGCAGGCACCTGTTTGGGAATTTTAGCAAATGATGGTGTTTTGCTTGCAGCAGAGAGACGCAAC<br>ATCCACAAGCTTCTTGATGAAGTCTTTTTTTCTGAAAAAATTTATAAACTCAATGAGGACATGGCTTGCAGTGTGGC<br>AGGCATAACTTCTGATGCTAATGTTCTGACTAATGAACTAAGGCTCATTGCTCAAAGGTATTTATTACAGTATCAGGA<br>GCCAATACCTTGTGAGCAGTTGGTTACAGCGCTGTGTGATATCAAACAAGCTTATACACAATTTGGAGGAAAACGT<br>CCCTTTGGTGTTTCATTGCTGTACATTGGCTGGGATAAGCACTATGGCTTTCAGCTCTATCAGAGTGACCCTAGTGG<br>AAATTACGGGGGATGGAAGGCCACATGCATTGGAAATAATAGCGCTGCAGCTGTGTCAATGTTGAAACAAGACTAT<br>AAAGAAGGAGAAATGACCTTGAAGTCAGCACTTGCTTTAGCTATCAAAGTACTAAATAAGACCATGGATGTTAGTA<br>AACTCTCTGCTGAAAAAGTGGAAATTGCAACACTAACAAGAGAGAATGGAAAGACAGTAATCAGAGTTCTCAAA<br>CAAAAAGAAGTGGAGCAGTTGATCAAAAAACATGAGGAAGAAGAAGCCAAAGCTGAGCGTGAGAAGAAAGAAA<br>AAGAACAGAAAGAAAAGGATAAACGATCCATGGAAGTGCAGCTGCAGCAGAGCGGAGCCGAGCTGGTGCGGAG<br>CGGGGCCTCTGTGAAACTGTCTTGCACCGCTTCTGGCTTCAACATCAAGGACTACTACATGCACTGGGTCAAGCAG<br>AGACCTGAGCAGGGCCTGGAATGGATCGGCTGGATGGACCCCAAGAACGGCGATACCGAGTACGCCCCTAAGTTC<br>CAGGGCAAGGCCACCATGACCGCCGACACCAGCAGCAATACCGCCTACCTGCAGCTGAGCAGCCTTACAAGCGAC<br>GACACAGCCGTGTACTACTGCCACGACGCCATCACCACGCCCACCAGAAACTTCGACGTGTGGGGCGCCGGCACA<br>ACCGTGACCGTGTCTTCTGGCGGCGGAGGCAGCGGCGGAGGCGGCAGCGGCGGCGGCGGCAGCGGAGGCGGCT<br>CTGATGTGGTGATGACCCAGACACCTCTGAGCCTGCCTGTGTCCCTGGGCGACCAGGCCAGCATCAGCTGTAGAA<br>GCTCTCAGAGCCTGGTGCACAGCAACGGCAACACCTACCTGCACTGGTACCTGCAAAAGCCCGGCCAGTCTCCAA<br>AGCTGCTGATGTACAAGGTGTCCAATAGATTCAGCGGTGTTCCTGATAGATTTAGCGGCAGCGGGAGCGGAACCGA<br>CTTCACCCTGAAGATCAGCAGAGTGGAAGCCGAAGATCTGGGAGTGTACTTCTGCAGCCAGTCAACACACGTGCC<br>TTTCACCTTCGGCTCCGGCACCAAGCTCGAGATCAAG |
| PSMA4-<br>$(GGGGS)_3$-<br>scFv(c-Fos) | ATGTCTCGAAGATATGACTCCAGGACCACTATATTTTCTCCAGAAGGTCGCTTATACCAAGTTGAATATGCCATGGA<br>AGCTATTGGACATGCAGGCACCTGTTTGGGAATTTTAGCAAATGATGGTGTTTTGCTTGCAGCAGAGAGACGCAAC<br>ATCCACAAGCTTCTTGATGAAGTCTTTTTTTCTGAAAAAATTTATAAACTCAATGAGGACATGGCTTGCAGTGTGGC<br>AGGCATAACTTCTGATGCTAATGTTCTGACTAATGAACTAAGGCTCATTGCTCAAAGGTATTTATTACAGTATCAGGA<br>GCCAATACCTTGTGAGCAGTTGGTTACAGCGCTGTGTGATATCAAACAAGCTTATACACAATTTGGAGGAAAACGT<br>CCCTTTGGTGTTTCATTGCTGTACATTGGCTGGGATAAGCACTATGGCTTTCAGCTCTATCAGAGTGACCCTAGTGG<br>AAATTACGGGGGATGGAAGGCCACATGCATTGGAAATAATAGCGCTGCAGCTGTGTCAATGTTGAAACAAGACTAT<br>AAAGAAGGAGAAATGACCTTGAAGTCAGCACTTGCTTTAGCTATCAAAGTACTAAATAAGACCATGGATGTTAGTA<br>AACTCTCTGCTGAAAAAGTGGAAATTGCAACACTAACAAGAGAGAATGGAAAGACAGTAATCAGAGTTCTCAAA<br>CAAAAAGAAGTGGAGCAGTTGATCAAAAAACATGAGGAAGAAGAAGCCAAAGCTGAGCGTGAGAAGAAAGAAA<br>AAGAACAGAAAGAAAAGGATAAACGATCCGGAGGAGGAGGGTCCGGTGGGGGAGGCTCTGGAGGGGGTGGGTC<br>TCGATCCATGGAAGTGCAGCTGCAGCAGAGCGGAGCCGAGCTGGTGCGGAGCGGGGCCTCTGTGAAACTGTCTT<br>GCACCGCTTCTGGCTTCAACATCAAGGACTACTACATGCACTGGGTCAAGCAGAGACCTGAGCAGGGCCTGGAAT<br>GGATCGGCTGGATGGACCCCAAGAACGGCGATACCGAGTACGCCCCTAAGTTCCAGGGCAAGGCCACCATGACCG<br>CCGACACCAGCAGCAATACCGCCTACCTGCAGCTGAGCAGCCTTACAAGCGACGACACAGCCGTGTACTACTGCC<br>ACGACGCCATCACCACGCCCACCAGAAACTTCGACGTGTGGGGCGCCGGCACAACCGTGACCGTGTCTTCTGGCG<br>GCGGAGGCAGCGGCGGAGGCGGCAGCGGCGGCGGCGGCAGCGGAGGCGGCTCTGATGTGGTGATGACCCAGAC<br>ACCTCTGAGCCTGCCTGTGTCCCTGGGCGACCAGGCCAGCATCAGCTGTAGAAGCTCTCAGAGCCTGGTGCACAG<br>CAACGGCAACACCTACCTGCACTGGTACCTGCAAAAGCCCGGCCAGTCTCCAAAGCTGCTGATGTACAAGGTGTC<br>CAATAGATTCAGCGGTGTTCCTGATAGATTTAGCGGCAGCGGGAGCGGAACCGACTTCACCCTGAAGATCAGCAG<br>AGTGGAAGCCGAAGATCTGGGAGTGTACTTCTGCAGCCAGTCAACACACGTGCCTTTCACCTTCGGCTCCGGCAC<br>CAAGCTCGAGATCAAG |

| | |
|---|---|
| PSMA4-<br>$(GGGGS)_6$-<br>scFv(c-Fos) | ATGTCTCGAAGATATGACTCCAGGACCACTATATTTTCTCCAGAAGGTCGCTTATACCAAGTTGAATATGCCATGGA<br>AGCTATTGGACATGCAGGCACCTGTTTGGGAATTTTAGCAAATGATGGTGTTTTGCTTGCAGCAGAGAGACGCAAC<br>ATCCACAAGCTTCTTGATGAAGTCTTTTTTTCTGAAAAAATTTATAAACTCAATGAGGACATGGCTTGCAGTGTGGC<br>AGGCATAACTTCTGATGCTAATGTTCTGACTAATGAACTAAGGCTCATTGCTCAAAGGTATTTATTACAGTATCAGGA<br>GCCAATACCTTGTGAGCAGTTGGTTACAGCGCTGTGTGATATCAAACAAGCTTATACACAATTTGGAGGAAAACGT<br>CCCTTTGGTGTTTCATTGCTGTACATTGGCTGGGATAAGCACTATGGCTTTCAGCTCTATCAGAGTGACCCTAGTGG<br>AAATTACGGGGGATGGAAGGCCACATGCATTGGAAATAATAGCGCTGCAGCTGTGTCAATGTTGAAACAAGACTAT<br>AAAGAAGGAGAAATGACCTTGAAGTCAGCACTTGCTTTAGCTATCAAAGTACTAAATAAGACCATGGATGTTAGTA<br>AACTCTCTGCTGAAAAAGTGGAAATTGCAACACTAACAAGAGAGAATGGAAAGACAGTAATCAGAGTTCTCAAA<br>CAAAAAGAAGTGGAGCAGTTGATCAAAAAACATGAGGAAGAAGAAGCCAAAGCTGAGCGTGAGAAGAAAGAAA<br>AAGAACAGAAAGAAAAGGATAAACGATCCGGCGGAGGTGGCAGCGGTGGTGGCGGATCTGGCGGAGGTGGCTCT<br>GGAGGAGGCGGTTCTGGCGGAGGCGGCTCAGGTGGAGGTGGCAGCCGATCCATGGAAGTGCAGCTGCAGCAGAG<br>CGGAGCCGAGCTGGTGCGGAGCGGGGCCTCTGTGAAACTGTCTTGCACCGCTTCTGGCTTCAACATCAAGGACTA<br>CTACATGCACTGGGTCAAGCAGAGACCTGAGCAGGGCCTGGAATGGATCGGCTGGATGGACCCCAAGAACGGCG<br>ATACCGAGTACGCCCCTAAGTTCCAGGGCAAGGCCACCATGACCGCCGACACCAGCAGCAATACCGCCTACCTGC<br>AGCTGAGCAGCCTTACAAGCGACGACACAGCCGTGTACTACTGCCACGACGCCATCACCACGCCCACCAGAAACT<br>TCGACGTGTGGGGCGCCGGCACAACCGTGACCGTGTCTTCTGGCGGCGGAGGCAGCGGCGGAGGCGGCAGCGGC<br>GGCGGCGGCAGCGGAGGCGGCTCTGATGTGGTGATGACCCAGACACCTCTGAGCCTGCCTGTGTCCCTGGGCGAC<br>CAGGCCAGCATCAGCTGTAGAAGCTCTCAGAGCCTGGTGCACAGCAACGGCAACACCTACCTGCACTGGTACCTG<br>CAAAAGCCCGGCCAGTCTCCAAAGCTGCTGATGTACAAGGTGTCCAATAGATTCAGCGGTGTTCCTGATAGATTTA<br>GCGGCAGCGGGAGCGGAACCGACTTCACCCTGAAGATCAGCAGAGTGGAAGCCGAAGATCTGGGAGTGTACTTC<br>TGCAGCCAGTCAACACACGTGCCTTTCACCTTCGGCTCCGGCACCAAGCTCGAGATCAAG |
| PSMB2-<br>scFv(c-Fos) | ATGGAGTACCTCATCGGTATCCAAGGCCCCGACTATGTTCTTGTCGCCTCCGACCGGGTGGCCGCCAGCAATATTGT<br>CCAGATGAAGGACGATCATGACAAGATGTTTAAGATGAGTGAAAAGATATTACTCCTGTGTGTTGGAGAGGCTGGA<br>GACACTGTACAGTTTGCAGAATATATTCAGAAAAACGTGCAACTTTATAAGATGCGAAATGGATATGAATTGTCTCC<br>CACGGCAGCAGCTAACTTCACACGCCGAAACCTGGCTGACTGTCTTCGGAGTCGGACCCCATATCATGTGAACCTC<br>CTCCTGGCTGGCTATGATGAGCATGAAGGGCCAGCGCTGTATTACATGGACTACCTGGCAGCCTTGGCCAAGGCCC<br>CTTTTGCAGCCCACGGCTATGGTGCCTTCCTGACTCTCAGTATCCTCGACCGATACTACACACCGACTATCTCACGT<br>GAGAGGGCAGTGGAACTCCTTAGGAAATGTCTGGAGGAGCTCCAGAAACGCTTCATCCTGAATCTGCCAACCTTC<br>AGTGTTCGAATCATTGACAAAAATGGCATCCATGACCTGGATAACATTTCCTTCCCCAAACAGGGCTCCAAGCTTG<br>GTACCGAGCTCGGATCCATGGAAGTGCAGCTGCAGCAGAGCGGAGCCGAGCTGGTGCGGAGCGGGGCCTCTGTG<br>AAACTGTCTTGCACCGCTTCTGGCTTCAACATCAAGGACTACTACATGCACTGGGTCAAGCAGAGACCTGAGCAG<br>GGCCTGGAATGGATCGGCTGGATGGACCCCAAGAACGGCGATACCGAGTACGCCCCTAAGTTCCAGGGCAAGGCC<br>ACCATGACCGCCGACACCAGCAGCAATACCGCCTACCTGCAGCTGAGCAGCCTTACAAGCGACGACACAGCCGTG<br>TACTACTGCCACGACGCCATCACCACGCCCACCAGAAACTTCGACGTGTGGGGCGCCGGCACAACCGTGACCGTG<br>TCTTCTGGCGGCGGAGGCAGCGGCGGAGGCGGCAGCGGCGGCGGCGGCAGCGGAGGCGGCTCTGATGTGGTGAT<br>GACCCAGACACCTCTGAGCCTGCCTGTGTCCCTGGGCGACCAGGCCAGCATCAGCTGTAGAAGCTCTCAGAGCCT<br>GGTGCACAGCAACGGCAACACCTACCTGCACTGGTACCTGCAAAAGCCCGGCCAGTCTCCAAAGCTGCTGATGTA<br>CAAGGTGTCCAATAGATTCAGCGGTGTTCCTGATAGATTTAGCGGCAGCGGGAGCGGAACCGACTTCACCCTGAA<br>GATCAGCAGAGTGGAAGCCGAAGATCTGGGAGTGTACTTCTGCAGCCAGTCAACACACGTGCCTTTCACCTTCGG<br>CTCCGGCACCAAGCTCGAGATCAAG |
| PSMB2-<br>$(GGGGS)_3$-<br>scFv(c-Fos) | ATGGAGTACCTCATCGGTATCCAAGGCCCCGACTATGTTCTTGTCGCCTCCGACCGGGTGGCCGCCAGCAATATTGT<br>CCAGATGAAGGACGATCATGACAAGATGTTTAAGATGAGTGAAAAGATATTACTCCTGTGTGTTGGAGAGGCTGGA<br>GACACTGTACAGTTTGCAGAATATATTCAGAAAAACGTGCAACTTTATAAGATGCGAAATGGATATGAATTGTCTCC<br>CACGGCAGCAGCTAACTTCACACGCCGAAACCTGGCTGACTGTCTTCGGAGTCGGACCCCATATCATGTGAACCTC<br>CTCCTGGCTGGCTATGATGAGCATGAAGGGCCAGCGCTGTATTACATGGACTACCTGGCAGCCTTGGCCAAGGCCC<br>CTTTTGCAGCCCACGGCTATGGTGCCTTCCTGACTCTCAGTATCCTCGACCGATACTACACACCGACTATCTCACGT<br>GAGAGGGCAGTGGAACTCCTTAGGAAATGTCTGGAGGAGCTCCAGAAACGCTTCATCCTGAATCTGCCAACCTTC<br>AGTGTTCGAATCATTGACAAAAATGGCATCCATGACCTGGATAACATTTCCTTCCCCAAACAGGGCTCCAAGCTTG<br>GTACCGAGCTCGGATCCGGAGGAGGAGGGTCCGGTGGGGGAGGCTCTGGAGGGGGTGGGTCTGGATCCATGGAA<br>GTGCAGCTGCAGCAGAGCGGAGCCGAGCTGGTGCGGAGCGGGGCCTCTGTGAAACTGTCTTGCACCGCTTCTGG<br>CTTCAACATCAAGGACTACTACATGCACTGGGTCAAGCAGAGACCTGAGCAGGGCCTGGAATGGATCGGCTGGAT<br>GGACCCCAAGAACGGCGATACCGAGTACGCCCCTAAGTTCCAGGGCAAGGCCACCATGACCGCCGACACCAGCA<br>GCAATACCGCCTACCTGCAGCTGAGCAGCCTTACAAGCGACGACACAGCCGTGTACTACTGCCACGACGCCATCA<br>CCACGCCCACCAGAAACTTCGACGTGTGGGGCGCCGGCACAACCGTGACCGTGTCTTCTGGCGGCGGAGGCAGC<br>GGCGGAGGCGGCAGCGGCGGCGGCGGCAGCGGAGGCGGCTCTGATGTGGTGATGACCCAGACACCTCTGAGCCT<br>GCCTGTGTCCCTGGGCGACCAGGCCAGCATCAGCTGTAGAAGCTCTCAGAGCCTGGTGCACAGCAACGGCAACA<br>CCTACCTGCACTGGTACCTGCAAAAGCCCGGCCAGTCTCCAAAGCTGCTGATGTACAAGGTGTCCAATAGATTCAG<br>CGGTGTTCCTGATAGATTTAGCGGCAGCGGGAGCGGAACCGACTTCACCCTGAAGATCAGCAGAGTGGAAGCCGA<br>AGATCTGGGAGTGTACTTCTGCAGCCAGTCAACACACGTGCCTTTCACCTTCGGCTCCGGCACCAAGCTCGAGATC<br>AAG |
| PSMB2-<br>$(GGGGS)_6$-<br>scFv(c-Fos) | ATGGAGTACCTCATCGGTATCCAAGGCCCCGACTATGTTCTTGTCGCCTCCGACCGGGTGGCCGCCAGCAATATTGT<br>CCAGATGAAGGACGATCATGACAAGATGTTTAAGATGAGTGAAAAGATATTACTCCTGTGTGTTGGAGAGGCTGGA<br>GACACTGTACAGTTTGCAGAATATATTCAGAAAAACGTGCAACTTTATAAGATGCGAAATGGATATGAATTGTCTCC<br>CACGGCAGCAGCTAACTTCACACGCCGAAACCTGGCTGACTGTCTTCGGAGTCGGACCCCATATCATGTGAACCTC<br>CTCCTGGCTGGCTATGATGAGCATGAAGGGCCAGCGCTGTATTACATGGACTACCTGGCAGCCTTGGCCAAGGCCC<br>CTTTTGCAGCCCACGGCTATGGTGCCTTCCTGACTCTCAGTATCCTCGACCGATACTACACACCGACTATCTCACGT<br>GAGAGGGCAGTGGAACTCCTTAGGAAATGTCTGGAGGAGCTCCAGAAACGCTTCATCCTGAATCTGCCAACCTTC |

| | |
|---|---|
| | AGTGTTCGAATCATTGACAAAAATGGCATCCATGACCTGGATAACATTTCCTTCCCCAAACAGGGCTCCAAGCTTG<br>GTACCGAGCTCGGATCCGGCGGAGGTGGCAGCGGTGGTGGCGGATCTGGCGGAGGTGGCTCTGGAGGAGGCGGT<br>TCTGGCGGAGGCGGCTCAGGTGGAGGTGGCAGCGGATCCATGGAAGTGCAGCTGCAGCAGAGCGGAGCCGAGCT<br>GGTGCGGAGCGGGGCCTCTGTGAAACTGTCTTGCACCGCTTCTGGCTTCAACATCAAGGACTACTACATGCACTG<br>GGTCAAGCAGAGACCTGAGCAGGGCCTGGAATGGATCGGCTGGATGGACCCCAAGAACGGCGATACCGAGTACG<br>CCCCTAAGTTCCAGGGCAAGGCCACCATGACCGCCGACACCAGCAGCAATACCGCCTACCTGCAGCTGAGCAGCC<br>TTACAAGCGACGACACAGCCGTGTACTACTGCCACGACGCCATCACCACGCCCACCAGAAACTTCGACGTGTGGG<br>GCGCCGGCACAACCGTGACCGTGTCTTCTGGCGGCGGAGGCAGCGGCGGAGGCGGCAGCGGCGGCGGCGGCAG<br>CGGAGGCGGCTCTGATGTGGTGATGACCCAGACACCTCTGAGCCTGCCTGTGTCCCTGGGCGACCAGGCCAGCAT<br>CAGCTGTAGAAGCTCTCAGAGCCTGGTGCACAGCAACGGCAACACCTACCTGCACTGGTACCTGCAAAAGCCCGG<br>CCAGTCTCCAAAGCTGCTGATGTACAAGGTGTCCAATAGATTCAGCGGTGTTCCTGATAGATTTAGCGGCAGCGGG<br>AGCGGAACCGACTTCACCCTGAAGATCAGCAGAGTGGAAGCCGAAGATCTGGGAGTGTACTTCTGCAGCCAGTC<br>AACACACGTGCCTTTCACCTTCGGCTCCGGCACCAAGCTCGAGATCAAG |
| scFv(c-Fos)-<br>PSMC5 | ATGGAAGTGCAGCTGCAGCAGAGCGGAGCCGAGCTGGTGCGGAGCGGGGCCTCTGTGAAACTGTCTTGCACCGC<br>TTCTGGCTTCAACATCAAGGACTACTACATGCACTGGGTCAAGCAGAGACCTGAGCAGGGCCTGGAATGGATCGG<br>CTGGATGGACCCCAAGAACGGCGATACCGAGTACGCCCCTAAGTTCCAGGGCAAGGCCACCATGACCGCCGACAC<br>CAGCAGCAATACCGCCTACCTGCAGCTGAGCAGCCTTACAAGCGACGACACAGCCGTGTACTACTGCCACGACGC<br>CATCACCACGCCCACCAGAAACTTCGACGTGTGGGGCGCCGGCACAACCGTGACCGTGTCTTCTGGCGGCGGAG<br>GCAGCGGCGGAGGCGGCAGCGGCGGCGGCGGCAGCGGAGGCGGCTCTGATGTGGTGATGACCCAGACACCTCTG<br>AGCCTGCCTGTGTCCCTGGGCGACCAGGCCAGCATCAGCTGTAGAAGCTCTCAGAGCCTGGTGCACAGCAACGGC<br>AACACCTACCTGCACTGGTACCTGCAAAAGCCCGGCCAGTCTCCAAAGCTGCTGATGTACAAGGTGTCCAATAGAT<br>TCAGCGGTGTTCCTGATAGATTTAGCGGCAGCGGGAGCGGAACCGACTTCACCCTGAAGATCAGCAGAGTGGAAG<br>CCGAAGATCTGGGAGTGTACTTCTGCAGCCAGTCAACACACGTGCCTTTCACCTTCGGCTCCGGCACCAAGCTCG<br>AGATCAAGGAATTCATGGCGCTTGACGGACCAGAGCAGATGGAGCTGGAGGAGGGGAAGGCAGGCAGCGGACTC<br>CGCCAATATTATCTGTCCAAGATTGAAGAACTCCAGCTGATTGTGAATGATAAGAGCCAAAACCTCCGGAGGCTGC<br>AGGCACAGAGGAACGAACTAAATGCTAAAGTTCGCCTATTGCGGGAGGAGCTACAGCTGCTGCAGGAGCAGGGC<br>TCCTATGTGGGGGAAGTAGTCCGGGCCATGGATAAGAAGAAAGTGTTGGTCAAGGTACATCCTGAAGGTAAATTTG<br>TTGTAGACGTGGACAAAAACATTGACATCAATGATGTGACACCCAATTGCCGGGTGGCTCTAAGGAATGACAGCTA<br>CACTCTGCACAAGATCCTGCCCAACAAGGTAGACCCATTAGTGTCACTGATGATGGTGGAGAAAGTACCAGATTCA<br>ACTTATGAGATGATTGGTGGACTGGACAAACAGATCAAGGAGATCAAAGAAGTGATCGAGCTGCCTGTTAAGCAT<br>CCTGAGCTCTTCGAAGCACTGGGCATTGCTCAGCCCAAGGGAGTGCTGCTGTATGGACCTCCAGGCACTGGGAAG<br>ACACTGTTGGCCCGGGCTGTGGCTCATCATACGGACTGTACCTTTATTCGTGTCTCTGGCTCTGAACTGGTACAGAA<br>ATTCATAGGGGAAGGGGCAAGAATGGTGAGGGAGCTGTTTGTCATGGCACGGGAACATGCTCCATCTATCATCTTC<br>ATGGACGAAATCGACTCCATCGGCTCCTCGCGGCTGGAGGGGGGTTCTGGAGGGGACAGTGAAGTGCAGCGCAC<br>GATGCTGGAGTTGCTCAACCAGCTCGACGGCTTTGAGGCCACCAAGAACATCAAGGTTATCATGGCTACTAATAGG<br>ATTGATATCCTGGACTCGGCACTGCTTCGCCCAGGGCGCATTGACAGAAAAATTGAATTCCCACCCCCCAATGAGG<br>AGGCCCGGCTGGACATTTTGAAGATTCATTCTCGGAAGATGAACCTGACCCGGGGGATCAACCTGAGAAAAATTG<br>CTGAGCTCATGCCAGGAGCATCAGGGGCTGAAGTGAAGGGCGTGTGCACAGAAGCTGGCATGTATGCCCTGCGAG<br>AACGGCGAGTCCATGTCACTCAGGAGGACTTTGAGATGGCAGTAGCCAAGGTCATGCAGAAGGACAGTGAGAAA<br>AACATGTCCATCAAGAAATTATGGAAG |
| scFv(c-Fos)-<br>(GGGGS)$_3$-<br>PSMC5 | ATGGAAGTGCAGCTGCAGCAGAGCGGAGCCGAGCTGGTGCGGAGCGGGGCCTCTGTGAAACTGTCTTGCACCGC<br>TTCTGGCTTCAACATCAAGGACTACTACATGCACTGGGTCAAGCAGAGACCTGAGCAGGGCCTGGAATGGATCGG<br>CTGGATGGACCCCAAGAACGGCGATACCGAGTACGCCCCTAAGTTCCAGGGCAAGGCCACCATGACCGCCGACAC<br>CAGCAGCAATACCGCCTACCTGCAGCTGAGCAGCCTTACAAGCGACGACACAGCCGTGTACTACTGCCACGACGC<br>CATCACCACGCCCACCAGAAACTTCGACGTGTGGGGCGCCGGCACAACCGTGACCGTGTCTTCTGGCGGCGGAG<br>GCAGCGGCGGAGGCGGCAGCGGCGGCGGCGGCAGCGGAGGCGGCTCTGATGTGGTGATGACCCAGACACCTCTG<br>AGCCTGCCTGTGTCCCTGGGCGACCAGGCCAGCATCAGCTGTAGAAGCTCTCAGAGCCTGGTGCACAGCAACGGC<br>AACACCTACCTGCACTGGTACCTGCAAAAGCCCGGCCAGTCTCCAAAGCTGCTGATGTACAAGGTGTCCAATAGAT<br>TCAGCGGTGTTCCTGATAGATTTAGCGGCAGCGGGAGCGGAACCGACTTCACCCTGAAGATCAGCAGAGTGGAAG<br>CCGAAGATCTGGGAGTGTACTTCTGCAGCCAGTCAACACACGTGCCTTTCACCTTCGGCTCCGGCACCAAGCTCG<br>AGATCAAGGAATTCGGAGGAGGAGGGTCCGGTGGGGGAGGCTCTGGAGGGGGTGGGTCTGAATTCATGGCGCTT<br>GACGGACCAGAGCAGATGGAGCTGGAGGAGGGGAAGGCAGGCAGCGGACTCCGCCAATATTATCTGTCCAAGAT<br>TGAAGAACTCCAGCTGATTGTGAATGATAAGAGCCAAAACCTCCGGAGGCTGCAGGCACAGAGGAACGAACTAA<br>ATGCTAAAGTTCGCCTATTGCGGGAGGAGCTACAGCTGCTGCAGGAGCAGGGCTCCTATGTGGGGGAAGTAGTCC<br>GGGCCATGGATAAGAAGAAAGTGTTGGTCAAGGTACATCCTGAAGGTAAATTTGTTGTAGACGTGGACAAAAACA<br>TTGACATCAATGATGTGACACCCAATTGCCGGGTGGCTCTAAGGAATGACAGCTACACTCTGCACAAGATCCTGCC<br>CAACAAGGTAGACCCATTAGTGTCACTGATGATGGTGGAGAAAGTACCAGATTCAACTTATGAGATGATTGGTGGA<br>CTGGACAAACAGATCAAGGAGATCAAAGAAGTGATCGAGCTGCCTGTTAAGCATCCTGAGCTCTTCGAAGCACTG<br>GGCATTGCTCAGCCCAAGGGAGTGCTGCTGTATGGACCTCCAGGCACTGGGAAGACACTGTTGGCCCGGGCTGTG<br>GCTCATCATACGGACTGTACCTTTATTCGTGTCTCTGGCTCTGAACTGGTACAGAAATTCATAGGGGAAGGGGCAA<br>GAATGGTGAGGGAGCTGTTTGTCATGGCACGGGAACATGCTCCATCTATCATCTTCATGGACGAAATCGACTCCATC<br>GGCTCCTCGCGGCTGGAGGGGGGTTCTGGAGGGGACAGTGAAGTGCAGCGCACGATGCTGGAGTTGCTCAACCA<br>GCTCGACGGCTTTGAGGCCACCAAGAACATCAAGGTTATCATGGCTACTAATAGGATTGATATCCTGGACTCGGCA<br>CTGCTTCGCCCAGGGCGCATTGACAGAAAAATTGAATTCCCACCCCCCAATGAGGAGGCCCGGCTGGACATTTTG<br>AAGATTCATTCTCGGAAGATGAACCTGACCCGGGGGATCAACCTGAGAAAAATTGCTGAGCTCATGCCAGGAGCA<br>TCAGGGGCTGAAGTGAAGGGCGTGTGCACAGAAGCTGGCATGTATGCCCTGCGAGAACGGCGAGTCCATGTCACT<br>CAGGAGGACTTTGAGATGGCAGTAGCCAAGGTCATGCAGAAGGACAGTGAGAAAAACATGTCCATCAAGAAATT<br>ATGGAAG |

| | |
|---|---|
| scFv(c-Fos)-<br>(GGGGS)$_6$-<br>PSMC5 | ATGGAAGTGCAGCTGCAGCAGAGCGGAGCCGAGCTGGTGCGGAGCGGGGCCTCTGTGAAACTGTCTTGCACCGC<br>TTCTGGCTTCAACATCAAGGACTACTACATGCACTGGGTCAAGCAGAGACCTGAGCAGGGCCTGGAATGGATCGG<br>CTGGATGGACCCCAAGAACGGCGATACCGAGTACGCCCCTAAGTTCCAGGGCAAGGCCACCATGACCGCCGACAC<br>CAGCAGCAATACCGCCTACCTGCAGCTGAGCAGCCTTACAAGCGACGACACAGCCGTGTACTACTGCCACGACGC<br>CATCACCACGCCCACCAGAAACTTCGACGTGTGGGGCGCCGGCACAACCGTGACCGTGTCTTCTGGCGGCGGAG<br>GCAGCGGCGGAGGCGGCAGCGGCGGCGGCGGCAGCGGAGGCGGCTCTGATGTGGTGATGACCCAGACACCTCTG<br>AGCCTGCCTGTGTCCCTGGGCGACCAGGCCAGCATCAGCTGTAGAAGCTCTCAGAGCCTGGTGCACAGCAACGGC<br>AACACCTACCTGCACTGGTACCTGCAAAAGCCCGGCCAGTCTCCAAAGCTGCTGATGTACAAGGTGTCCAATAGAT<br>TCAGCGGTGTTCCTGATAGATTTAGCGGCAGCGGGAGCGGAACCGACTTCACCCTGAAGATCAGCAGAGTGGAAG<br>CCGAAGATCTGGGAGTGTACTTCTGCAGCCAGTCAACACACGTGCCTTTCACCTTCGGCTCCGGCACCAAGCTCG<br>AGATCAAGGAATTCGGCGGAGGTGGCAGCGGTGGTGGCGGATCTGGCGGAGGTGGCTCTGGAGGAGGCGGTTCT<br>GGCGGAGGCGGCTCAGGTGGAGGTGGCAGCGAATTCATGGCGCTTGACGGACCAGAGCAGATGGAGCTGGAGGA<br>GGGGAAGGCAGGCAGCGGACTCCGCCAATATTATCTGTCCAAGATTGAAGAACTCCAGCTGATTGTGAATGATAAG<br>AGCCAAAACCTCCGGAGGCTGCAGGCACAGAGGAACGAACTAAATGCTAAAGTTCGCCTATTGCGGGAGGAGCT<br>ACAGCTGCTGCAGGAGCAGGGCTCCTATGTGGGGGAAGTAGTCCGGGCCATGGATAAGAAGAAAGTGTTGGTCAA<br>GGTACATCCTGAAGGTAAATTTGTTGTAGACGTGGACAAAAACATTGACATCAATGATGTGACACCCAATTGCCGG<br>GTGGCTCTAAGGAATGACAGCTACACTCTGCACAAGATCCTGCCCAACAAGGTAGACCCATTAGTGTCACTGATGA<br>TGGTGGAGAAAGTACCAGATTCAACTTATGAGATGATTGGTGGACTGGACAAACAGATCAAGGAGATCAAAGAAG<br>TGATCGAGCTGCCTGTTAAGCATCCTGAGCTCTTCGAAGCACTGGGCATTGCTCAGCCCAAGGGAGTGCTGCTGTA<br>TGGACCTCCAGGCACTGGGAAGACACTGTTGGCCCGGGCTGTGGCTCATCATACGGACTGTACCTTTATTCGTGTC<br>TCTGGCTCTGAACTGGTACAGAAATTCATAGGGGAAGGGGCAAGAATGGTGAGGGAGCTGTTTGTCATGGCACGG<br>GAACATGCTCCATCTATCATCTTCATGGACGAAATCGACTCCATCGGCTCCTCGCGGCTGGAGGGGGGTTCTGGAG<br>GGGACAGTGAAGTGCAGCGCACGATGCTGGAGTTGCTCAACCAGCTCGACGGCTTTGAGGCCACCAAGAACATC<br>AAGGTTATCATGGCTACTAATAGGATTGATATCCTGGACTCGGCACTGCTTCGCCCAGGGCGCATTGACAGAAAAAT<br>TGAATTCCCACCCCCCAATGAGGAGGCCCGGCTGGACATTTTGAAGATTCATTCTCGGAAGATGAACCTGACCCGG<br>GGGATCAACCTGAGAAAAATTGCTGAGCTCATGCCAGGAGCATCAGGGGCTGAAGTGAAGGGCGTGTGCACAGA<br>AGCTGGCATGTATGCCCTGCGAGAACGGCGAGTCCATGTCACTCAGGAGGACTTTGAGATGGCAGTAGCCAAGGT<br>CATGCAGAAGGACAGTGAGAAAAACATGTCCATCAAGAAATTATGGAAG |
| PSMD4-<br>scFv(Flag) | ATGGTGTTGGAAAGCACTATGGTGTGTGTGGACAACAGTGAGTATATGCGGAATGGAGACTTCTTACCCACCAGGC<br>TGCAGGCCCAGCAGGATGCTGTCAACATAGTTTGTCATTCAAAGACCCGCAGCAACCCTGAGAACAACGTGGGCC<br>TTATCACACTGGCTAATGACTGTGAAGTGCTGACCACACTCACCCCAGACACTGGCCGTATCCTGTCCAAGCTACA<br>TACTGTCCAACCCAAGGGCAAGATCACCTTCTGCACGGGCATCCGCGTGGCCCATCTGGCTCTGAAGCACCGACA<br>AGGCAAGAATCACAAGATGCGCATCATTGCCTTTGTGGGAAGCCCAGTGGAGGACAATGAGAAGGATCTGGTGAA<br>ACTGGCTAAACGCCTCAAGAAGGAGAAAGTAAATGTTGACATTATCAATTTTGGGGAAGAGGAGGTGAACACAGA<br>AAAGCTGACAGCCTTTGTAAACACGTTGAATGGCAAAGATGGAACCGGTTCTCATCTGGTGACAGTGCCTCCTGG<br>GCCCAGTTTGGCTGATGCTCTCATCAGTTCTCCGATTTTGGCTGGTGAAGGTGGTGCCATGCTGGGTCTTGGTGCC<br>AGTGACTTTGAATTTGGAGTAGATCCCAGTGCTGATCCTGAGCTGGCCTTGGCCCTTCGTGTATCTATGGAAGAGC<br>AGCGGCAGCGGCAGGAGGAGGAGGCCCGGCGGGCAGCTGCAGCTTCTGCTGCTGAGGCCGGGATTGCTACGACT<br>GGGACTGAAGACTCAGACGATGCCCTGCTGAAGATGACCATCAGCCAGCAAGAGTTTGGCCGCACTGGGCTTCCT<br>GACCTAAGCAGTATGACTGAGGAAGAGCAGATTGCTTATGCCATGCAGATGTCCCTGCAGGGAGCAGAGTTTGGC<br>CAGGCGGAATCAGCAGACATTGATGCCAGCTCAGCTATGGACACATCTGAGCCAGCCAAGGAGGAGGATGATTAC<br>GACGTGATGCAGGACCCCGAGTTCCTTCAGAGTGTCCTAGAGAACCTCCCAGGTGTGGATCCCAACAATGAAGCC<br>ATTCGAAATGCTATGGGCTCCCTGGCCTCCCAGGCCACCAAGGACGGCAAGAAGGACAAGAAGGAGGAAGACAA<br>GAAGCGACCCTTAATTAACGAGGTGAAGCTGGTGGAGTCTGGGGGAGGCTTAGTGAAGCCTGGAGGGTCCCTGA<br>AACTCTCCTGTGCAGCCTCTGGATTCACTTTCAGTAGCTTTGGAATGCACTGGGTTCGCCAGACTCCGGAGAAGAG<br>GCTGGAGTGGGTCGCATACATAAGTAGTGGCAGTAGTACCATCTACTATGCAGACACAGTGAAGGGCCGATTCACC<br>ATCTCCAGAGACAATGCCAAGAACACCCTGTACCTGCAAATGAGCAGTCTGAGGTCTGAGGACACGGCCATATATT<br>ACTGTGCAAGATCCCTTGCTACGGCCGCTTTTGCTTACTGGGGCCAAGGCACCACTCTCACAGTCTCCTCAGGTGG<br>AGGCGGTTCAGGCGGAGGTGGCTCTGGCGGTGGCGGATCGGACATCGTTCTGACACAGAGTCCCGCCAGCCTTAC<br>TGTCTCACTCGGCCAGAGGGCTACCATTTCCTGCAGATCCTCTCAAAGCATAGTGTACAGTAATGGAAACACCTATT<br>TAGAATGGTACCAACAGAAACCAGGACAGCCACCCAAACTCCTCATCTATAAAGTTTCCAACCGATTTTCTGGGAT<br>CCCAGCCAGGTTTAGTGGCAGTGGGTCTGGGACAGACTTCACCCTCAACATCCATCCTGTGGAGGAGGAGGATGC<br>TGCAACCTATTACTGTTTTCAAGGTTCACATGTTCCGTACACGTTCGGTGCTGGGACCAAGCTGGAAATAAAA |
| scFv(HA)-<br>ADRM1 | ATGGAGGTGAAGCTGGTGGAGTCTGGGGGAGGCTTAGTGAAGCCTGGAGGGTCCCTGAAACTCTCCTGTGCAGC<br>CTCTGGATTCACTTTCAGTAGTTATGGCATGTCTTGGGTTCGCCAGACTCCGGAGAAGAGGCTGGAGTGGGTCGCA<br>ACCATTAGTAGAGGTGGTAGTTATACCTACTATCCAGACAGTGTGAAGGGGCGATTCACCATCTCCAGAGACAATG<br>CCAAGAACACCCTGTACCTGCAAATGAGCAGTCTGAGGTCTGAGGACACGGCCATATATTACTGTGCAAGACGGG<br>AAACATACGATGAAAAGGGCTTTGCCTACTGGGGCCAAGGCACCACTCTCACAGTCTCCTCAGGTGGAGGCGGTT<br>CAGGCGGAGGTGGCTCTGGCGGTGGCGGATCGGACATTGTGCTGACCCAATCTCCAGCTTCTTTGACTGTGTCTCT<br>AGGGCAGAGGGCCACCATCTCCTGCAAGTCCAGCCAAAGTCTTCTGAATAGTGGTAATCAGAAGAACTACCTCAC<br>CTGGTACCAACAGAAACCAGGACAGCCACCCAAACTCCTCATCTATTGGGCATCCACTAGGGAATCTGGGATCCCA<br>GCCAGGTTTAGTGGCAGTGGGTCTGGGACAGACTTCACCCTCAACATCCATCCTGTGGAGGAGGAGGATGCTGCA<br>ACCTATTACTGTCAGAACGATAATTCCCATCCGCTCACGTTCGGTGCTGGGACCAAGCTGGAAATAAAAGGCGCGC<br>CAATGACGACCTCAGGCGCGCTCTTTCCAAGCCTGGTGCCAGGCTCTCGGGGCGCCTCCAACAAGTACTTGGTGG<br>AGTTTCGGGCGGGAAAGATGTCCCTGAAGGGGACCACCGTGACTCCGGATAAGCGGAAAGGGCTGGTGTACATTC<br>AGCAGACGGACGACTCGCTTATTCACTTCTGCTGGAAGGACAGGACGTCCGGGAACGTGGAAGACGACTTGATCA<br>TCTTCCCTGACGACTGTGAGTTCAAGCGGGTGCCGCAGTGCCCCAGCGGGAGGGTCTACGTGCTGAAGTTCAAGG<br>CAGGGTCCAAGCGGCTTTTCTTCTGGATGCAGGAACCCAAGACAGACCAGGATGAGGAGCATTGCCGGAAAGTC |

| | |
|---|---|
| | AACGAGTATCTGAACAACCCCCCGATGCCTGGGGCGCTGGGGGCCAGCGGAAGCAGCGGCCACGAACTCTCTGC<br>GCTAGGCGGTGAGGGTGGCCTGCAGAGCCTGCTGGGAAACATGAGCCACAGCCAGCTCATGCAGCTCATCGGACC<br>AGCCGGCCTTGGAGGACTGGGTGGGCTGGGGGCCCTGACTGGACCTGGCCTGGCCAGCTTACTGGGGAGCAGTG<br>GGCCTCCAGGGAGCAGCTCCTCCTCCAGCTCCCGGAGCCAGTCGGCAGCGGTCACCCCGTCATCCACCACCTCTT<br>CCACCCGTGCCACCCCAGCCCCTTCTGCTCCAGCAGCTGCCTCAGCAACTAGCCCGAGCCCCGCGCCCAGTTCCG<br>GGAATGGAGCCAGCACAGCAGCCAGCCCGACCCAGCCCATCCAGCTGAGCGACCTCCAGAGCATCCTGGCCACG<br>ATGAACGTACCAGCCGGGCCAGCAGGCGGCCAGCAAGTGGACCTGGCCAGTGTGCTGACGCCGGAGATAATGGC<br>TCCCATCCTCGCCAACGCGGATGTCCAGGAGCGCCTGCTTCCCTACTTGCCATCTGGGGAGTCGCTGCCGCAGACC<br>GCGGATGAGATCCAGAATACCCTGACCTCGCCCCAGTTCCAGCAGGCCCTGGGCATGTTCAGCGCAGCCTTGGCCT<br>CGGGGCAGCTGGGCCCCCTCATGTGCCAGTTCGGTCTGCCTGCAGAGGCTGTGGAGGCCGCCAACAAGGGCGATG<br>TGGAAGCGTTTGCCAAAGCCATGCAGAACAACGCCAAGCCCGAGCAGAAAGAGGGCGACACGAAGGACAAGAA<br>GGACGAAGAGGAGGACATGAGCCTGGAC |
| VHH(GFP)-<br>ADRM1 | ATGGTCCAACTGGTGGAGTCTGGTGGCGCTTTGGTGCAGCCAGGTGGCTCTCTGCGTTTGTCCTGTGCCGCTTCTG<br>GCTTCCCAGTGAACCGCTATTCCATGCGCTGGTATCGCCAGGCTCCAGGCAAAGAGCGTGAGTGGGTAGCCGGTAT<br>GTCCAGCGCGGGTGATCGTAGCTCCTATGAAGACTCCGTGAAGGGCCGTTTCACCATCAGCCGTGACGATGCCCGT<br>AACACGGTGTATCTGCAAATGAACAGCTTGAAACCTGAAGATACGGCCGTGTATTACTGTAATGTGAACGTGGGCT<br>TCGAGTATTGGGGCCAAGGCACCCAGGTCACCGTCTCCAGCGGCGCGCCAATGACGACCTCAGGCGCGCTCTTTC<br>CAAGCCTGGTGCCAGGCTCTCGGGGCGCCTCCAACAAGTACTTGGTGGAGTTTCGGGCGGGAAAGATGTCCCTGA<br>AGGGGACCACCGTGACTCCGGATAAGCGGAAAGGGCTGGTGTACATTCAGCAGACGGACGACTCGCTTATTCACT<br>TCTGCTGGAAGGACAGGACGTCCGGGAACGTGGAAGACGACTTGATCATCTTCCCTGACGACTGTGAGTTCAAGC<br>GGGTGCCGCAGTGCCCCAGCGGGAGGGTCTACGTGCTGAAGTTCAAGGCAGGGTCCAAGCGGCTTTTCTTCTGGA<br>TGCAGGAACCCAAGACAGACCAGGATGAGGAGCATTGCCGGAAAGTCAACGAGTATCTGAACAACCCCCCGATG<br>CCTGGGGCGCTGGGGGCCAGCGGAAGCAGCGGCCACGAACTCTCTGCGCTAGGCGGTGAGGGTGGCCTGCAGAG<br>CCTGCTGGGAAACATGAGCCACAGCCAGCTCATGCAGCTCATCGGACCAGCCGGCCTTGGAGGACTGGGTGGGCT<br>GGGGGCCCTGACTGGACCTGGCCTGGCCAGCTTACTGGGGAGCAGTGGGCCTCCAGGGAGCAGCTCCTCCTCCA<br>GCTCCCGGAGCCAGTCGGCAGCGGTCACCCCGTCATCCACCACCTCTTCCACCCGTGCCACCCCAGCCCCTTCTGC<br>TCCAGCAGCTGCCTCAGCAACTAGCCCGAGCCCCGCGCCCAGTTCCGGGAATGGAGCCAGCACAGCAGCCAGCC<br>CGACCCAGCCCATCCAGCTGAGCGACCTCCAGAGCATCCTGGCCACGATGAACGTACCAGCCGGGCCAGCAGGCG<br>GCCAGCAAGTGGACCTGGCCAGTGTGCTGACGCCGGAGATAATGGCTCCCATCCTCGCCAACGCGGATGTCCAGG<br>AGCGCCTGCTTCCCTACTTGCCATCTGGGGAGTCGCTGCCGCAGACCGCGGATGAGATCCAGAATACCCTGACCTC<br>GCCCCAGTTCCAGCAGGCCCTGGGCATGTTCAGCGCAGCCTTGGCCTCGGGGCAGCTGGGCCCCCTCATGTGCCA<br>GTTCGGTCTGCCTGCAGAGGCTGTGGAGGCCGCCAACAAGGGCGATGTGGAAGCGTTTGCCAAAGCCATGCAGA<br>ACAACGCCAAGCCCGAGCAGAAAGAGGGCGACACGAAGGACAAGAAGGACGAAGAGGAGGACATGAGCCTGG<br>AC |
| PSMD4-<br>VHH(GFP) | ATGGTGTTGGAAAGCACTATGGTGTGTGTGGACAACAGTGAGTATATGCGGAATGGAGACTTCTTACCCACCAGGC<br>TGCAGGCCCAGCAGGATGCTGTCAACATAGTTTGTCATTCAAAGACCCGCAGCAACCCTGAGAACAACGTGGGCC<br>TTATCACACTGGCTAATGACTGTGAAGTGCTGACCACACTCACCCCAGACACTGGCCGTATCCTGTCCAAGCTACA<br>TACTGTCCAACCCAAGGGCAAGATCACCTTCTGCACGGGCATCCGCGTGGCCCATCTGGCTCTGAAGCACCGACA<br>AGGCAAGAATCACAAGATGCGCATCATTGCCTTTGTGGGAAGCCCAGTGGAGGACAATGAGAAGGATCTGGTGAA<br>ACTGGCTAAACGCCTCAAGAAGGAGAAAGTAAATGTTGACATTATCAATTTTGGGGAAGAGGAGGTGAACACAGA<br>AAAGCTGACAGCCTTTGTAAACACGTTGAATGGCAAAGATGGAACCGGTTCTCATCTGGTGACAGTGCCTCCTGG<br>GCCCAGTTTGGCTGATGCTCTCATCAGTTCTCCGATTTTGGCTGGTGAAGGTGGTGCCATGCTGGGTCTTGGTGCC<br>AGTGACTTTGAATTTGGAGTAGATCCCAGTGCTGATCCTGAGCTGGCCTTGGCCCTTCGTGTATCTATGGAAGAGC<br>AGCGGCAGCGGCAGGAGGAGGAGGCCCGGCGGGCAGCTGCAGCTTCTGCTGCTGAGGCCGGGATTGCTACGACT<br>GGGACTGAAGACTCAGACGATGCCCTGCTGAAGATGACCATCAGCCAGCAAGAGTTTGGCCGCACTGGGCTTCCT<br>GACCTAAGCAGTATGACTGAGGAAGAGCAGATTGCTTATGCCATGCAGATGTCCCTGCAGGGAGCAGAGTTTGGC<br>CAGGCGGAATCAGCAGACATTGATGCCAGCTCAGCTATGGACACATCTGAGCCAGCCAAGGAGGAGGATGATTAC<br>GACGTGATGCAGGACCCCGAGTTCCTTCAGAGTGTCCTAGAGAACCTCCCAGGTGTGGATCCCAACAATGAAGCC<br>ATTCGAAATGCTATGGGCTCCCTGGCCTCCCAGGCCACCAAGGACGGCAAGAAGGACAAGAAGGAGGAAGACAA<br>GAAGCGACCCTTAATTAACGTCCAACTGGTGGAGTCTGGTGGCGCTTTGGTGCAGCCAGGTGGCTCTCTGCGTTTG<br>TCCTGTGCCGCTTCTGGCTTCCCAGTGAACCGCTATTCCATGCGCTGGTATCGCCAGGCTCCAGGCAAAGAGCGTG<br>AGTGGGTAGCCGGTATGTCCAGCGCGGGTGATCGTAGCTCCTATGAAGACTCCGTGAAGGGCCGTTTCACCATCAG<br>CCGTGACGATGCCCGTAACACGGTGTATCTGCAAATGAACAGCTTGAAACCTGAAGATACGGCCGTGTATTACTGT<br>AATGTGAACGTGGGCTTCGAGTATTGGGGCCAAGGCACCCAGGTCACCGTCTCCAGC |
| ADRM1-<br>scFv(BRD4) | ATGACGACCTCAGGCGCGCTCTTTCCAAGCCTGGTGCCAGGCTCTCGGGGCGCCTCCAACAAGTACTTGGTGGAG<br>TTTCGGGCGGGAAAGATGTCCCTGAAGGGGACCACCGTGACTCCGGATAAGCGGAAAGGGCTGGTGTACATTCAG<br>CAGACGGACGACTCGCTTATTCACTTCTGCTGGAAGGACAGGACGTCCGGGAACGTGGAAGACGACTTGATCATC<br>TTCCCTGACGACTGTGAGTTCAAGCGGGTGCCGCAGTGCCCCAGCGGGAGGGTCTACGTGCTGAAGTTCAAGGCA<br>GGGTCCAAGCGGCTTTTCTTCTGGATGCAGGAACCCAAGACAGACCAGGATGAGGAGCATTGCCGGAAAGTCAA<br>CGAGTATCTGAACAACCCCCCGATGCCTGGGGCGCTGGGGGCCAGCGGAAGCAGCGGCCACGAACTCTCTGCGCT<br>AGGCGGTGAGGGTGGCCTGCAGAGCCTGCTGGGAAACATGAGCCACAGCCAGCTCATGCAGCTCATCGGACCAG<br>CCGGCCTTGGAGGACTGGGTGGGCTGGGGGCCCTGACTGGACCTGGCCTGGCCAGCTTACTGGGGAGCAGTGGG<br>CCTCCAGGGAGCAGCTCCTCCTCCAGCTCCCGGAGCCAGTCGGCAGCGGTCACCCCGTCATCCACCACCTCTTCC<br>ACCCGTGCCACCCCAGCCCCTTCTGCTCCAGCAGCTGCCTCAGCAACTAGCCCGAGCCCCGCGCCCAGTTCCGGG<br>AATGGAGCCAGCACAGCAGCCAGCCCGACCCAGCCCATCCAGCTGAGCGACCTCCAGAGCATCCTGGCCACGATG<br>AACGTACCAGCCGGGCCAGCAGGCGGCCAGCAAGTGGACCTGGCCAGTGTGCTGACGCCGGAGATAATGGCTCC<br>CATCCTCGCCAACGCGGATGTCCAGGAGCGCCTGCTTCCCTACTTGCCATCTGGGGAGTCGCTGCCGCAGACCGCG<br>GATGAGATCCAGAATACCCTGACCTCGCCCCAGTTCCAGCAGGCCCTGGGCATGTTCAGCGCAGCCTTGGCCTCGG |

| | |
|---|---|
| | GGCAGCTGGGCCCCCTCATGTGCCAGTTCGGTCTGCCTGCAGAGGCTGTGGAGGCCGCCAACAAGGGCGATGTGG<br>AAGCGTTTGCCAAAGCCATGCAGAACAACGCCAAGCCCGAGCAGAAAGAGGGCGACACGAAGGACAAGAAGGA<br>CGAAGAGGAGGACATGAGCCTGGACTTAATTAACGAGGTGCAATTGTTGGAGAGCGGGGGAGGCTTGGTACAGC<br>CTGGGGGGTCCCTGCGCCTCTCCTGTGCAGCCAGCGGATTCACCTTTTACGGTTATGGTATGTCTTGGGTCCGCCAG<br>GCTCCAGGGAAGGGGCTGGAGTGGGTCTCATACATTGGTGGTTACGGTAGTTCTACATCTTATGCAGACTCCGTGA<br>AGGGCCGGTTCACCATCTCCCGTGACAATTCCAAGAACACGCTGTATCTGCAAATGAACAGCCTGCGTGCCGAGG<br>ACACGGCTGTATATTATTGTGCGCGCGGTTCTGCTTACATTGACTATTGGGGCCAAGGAACCCTGGTCACCGTCTCC<br>TCAGGTGGAGGCGGTTCAGGCGGAGGTGGATCCGGCGGTGGCGGATCGGACATCCAGATGACCCAGTCTCCATCC<br>TCCCTGAGCGCATCTGTAGGAGACCGCGTCACCATCACCTGCAGGGCAAGTCAGAGCATTAGCAGCTATTTAAATT<br>GGTATCAGCAGAAACCAGGGAAAGCCCCTAAGCTCCTGATCTATGCTGCATCCAGTTTGCAAAGTGGGGTCCCATC<br>ACGTTTCAGTGGCAGTGGAAGCGGGACAGATTTCACTCTCACCATCAGCAGTCTGCAACCTGAAGATTTTGCAAC<br>TTATTACTGTCAACAGTACAACTCTCTGTACACTTTTGGCCAGGGGACCAAGCTGGAGATCAAA |
| mutPSMD4(6) | ATGGTGTTGGAAAGCACTATGGTGTGTGTGGCTAACGCTGCTGCTATGCGGAATGGAGACTTCTTACCCACCAGGC<br>TGCAGGCCCAGCAGGATGCTGTCAACATAGTTTGTCATTCAAAGACCCGCAGCAACCCTGAGAACAACGTGGGCC<br>TTATCACACTGGCTAATGACTGTGAAGTGCTGACCACACTCACCCCAGACACTGGCCGTATCCTGTCCAAGCTACA<br>TACTGTCCAACCCAAGGGCAAGATCACCTTCTGCACGGGCATCCGCGTGGCCCATCTGGCTCTGAAGCACCGACA<br>AGGCAAGAATCACAAGATGCGCATCATTGCCTTTGTGGCTGCTCCAGTGGAGGACAATGAGAAGGATCTGGTGAA<br>ACTGGCTAAACGCCTCAAGAAGGAGAAAGTAAATGTTGACATTATCAATTTTGGGGAAGAGGAGGTGAACACAGA<br>AAAGCTGACAGCCTTTGTAAACACGTTGAATGGCAAAGATGGAACCGGTTCTCATCTGGTGACAGTGCCTCCTGG<br>GCCCAGTTTGGCTGATGCTCTCATCAGTTCTCCGATTTTGGCTGGTGAAGGTGGTGCCATGCTGGGTCTTGGTGCC<br>AGTGACTTTGAATTTGGAGTAGATCCCAGTGCTGATCCTGAGCTGGCCTTGGCCCTTCGTGTATCTATGGAAGAGC<br>AGCGGCAGCGGCAGGAGGAGGAGGCCCGGCGGGCAGCTGCAGCTTCTGCTGCTGAGGCCGGGATTGCTACGACT<br>GGGACTGAAGACTCAGACGATGCCCTGCTGAAGATGACCATCAGCCAGCAAGAGTTTGGCCGCACTGGGCTTCCT<br>GACCTAAGCAGTATGACTGAGGAAGAGCAGATTGCTTATGCCATGCAGATGTCCCTGCAGGGAGCAGAGTTTGGC<br>CAGGCGGAATCAGCAGACATTGATGCCAGCTCAGCTATGGACACATCTGAGCCAGCCAAGGAGGAGGATGATTAC<br>GACGTGATGCAGGACCCCGAGTTCCTTCAGAGTGTCCTAGAGAACCTCCCAGGTGTGGATCCCAACAATGAAGCC<br>ATTCGAAATGCTATGGGCTCCCTGGCCTCCCAGGCCACCAAGGACGGCAAGAAGGACAAGAAGGAGGAAGACAA<br>GAAGCGACCC |
| ADRM1(ΔPru) | ATGCCGCCTATGCCCGGAGCACTCGGAGCATCTGGGAGTAGTGGGCATGAGTTGTCCGCTCTCGGTGGAGAAGGC<br>GGGTTACAATCTCTTCTCGGGAATATGTCTCATTCTCAACTGATGCAATTAATTGGGCCGGCTGGACTGGGCGGTCT<br>CGGCGGCCTTGGAGCATTGACCGGGCCAGGGCTCGCTTCACTCTTAGGATCTTCTGGTCCCCCGGGTAGTTCATCT<br>TCATCATCAAGCCGATCACAAAGTGCTGCTGTGACACCCTCTAGCACTACGAGCAGCACTAGGGCAACGCCTGCA<br>CCTTCCGCACCTGCGGCAGCATCCGCCACATCACCTAGTCCTGCTCCTAGCAGTGGCAACGGCGCAAGTACTGCCG<br>CTTCTCCTACTCAACCAATTCAATTGAGTGATCTGCAAAGTATTTTGGCAACCATGAATGTTCCCGCTGGACCTGCG<br>GGCGGACAACAGGTCGATCTCGCGAGCGTATTGACCCCCGAAATCATGGCCCCAATATTAGCTAATGCTGACGTGC<br>AAGAACGGCTCCTGCCATATCTGCCTTCCGGCGAATCTTTACCTCAAACTGCCGACGAAATTCAAAACACTCTTAC<br>TTCACCGCAATTTCAACAAGCTTTAGGAATGTTTTCCGCGGCACTCGCTTCTGGACAATTAGGTCCTTTGATGTGTC<br>AATTTGGCCTTCCAGCCGAAGCAGTAGAAGCGGCTAATAAAGGAGACGTAGAGGCCTTCGCAAAGGCTATGCAAA<br>ATAATGCAAAACCAGAACAAAAGGAAGGGGACACTAAAGATAAGAAAGATGAGGAAGAAGATATGAGTCTTGAT |
| scFv(IgG1) | ATGGAAGTACAATTACAACAATCCGGAGCGGAATTGGTGAGGTCTGGCGCGTCCGTGAAGTTAAGCTGTACCGCG<br>AGTGGCTTCAACATAAAGCGCTATGGAATTAAATGGGTTAAACAAAGGCCAGAGCAGGGTCTCGAATGGATCGGC<br>TGGATCAACACTAGGAGTGGCGTGCCGGCGTATGCTCAAGGTTTCACCGGAAAGGCGACCATGACTGCAGATACT<br>TCTTCAAACACTGCTTACCTTCAACTTTCATCATTGACCAGTGACGATACCGCCGTATACTACTGCCACGATAAAAC<br>AGAATACTGGGAAGATGGATTTGACGTCTGGGGAGCAGGTACGACCGTAACCGTAAGCTCTGGTGGTGGTGGCTC<br>AGGTGGCGGTGGCTCTGGCGGCGGTGGTTCTGGTGGTGGTTCCGATGTCGTAATGACACAAACACCCCTCTCTCTC<br>CCTGTATCCCTTGGGGATCAAGCATCAATAAGCTGTTCCGGAAGTAGTTCAAATATTGGGAATTCATACGTTTACTG<br>GTACCTGCAAAAGCCCGGCCAATCCCCTAAATTGCTGATGTACAGGAACAATAGACGCCCATCTGGGGTGCCAGAC<br>AGATTCAGCGGAAGCGGGAGCGGGACAGATTTTACGTTGAAGATTTCAAGGGTCGAGGCAGAAGATCTTGGAGTG<br>TACTTTTGCGCTACCTGGGACGACAGTTTGAGCGGGCGCCTGTTTGGGTCCGGCACCAAGCTCGAGATCAAACAT<br>CATCACCACCATCACGACTACGATATACCCACAACCGCCAGCGAGAATCTGTACTTTCAAGGAGAGTTAGGGATGA<br>GGGGTTCAGCTGGAAAGGCCGGTGAAGGTGAAATCCCTGCCCCTCTTGCTGGTACCGTTTCTAAGATACTGGTAAA<br>AGAAGGTGACACTGTTAAAGCTGGTCAAACAGTTCTGGTGCTGGAGGCTATGAAAATGGAGACCGAAATTAACGC<br>TCCTACTGACGGAAAAGTTGAAAAGGTGTTAGTTAAGGAAAGAAATGCTGTTCAAGGTGGTCAAGGGTTAATCAA<br>GATCGGCGTTTGA |
